# Posterior marginalization accelerates Bayesian inference for dynamical systems

**DOI:** 10.1101/2022.12.02.518841

**Authors:** Elba Raimúndez, Michael Fedders, Jan Hasenauer

**Affiliations:** Life and Medical Sciences (LIMES) Institute, University of Bonn, Bonn, Germany; Technische Universität München, Center for Mathematics, Garching, Germany; Helmholtz Zentrum München - German Research Center for Environmental Health, Computational Health Center, Neuherberg, Germany

## Abstract

Bayesian inference is an important method in the life and natural sciences for learning from data. It provides information about parameter uncertainties, and thereby the reliability of models and their predictions. Yet, generating representative samples from the Bayesian posterior distribution is often computationally challenging. Here, we present an approach that lowers the computational complexity of sample generation for problems with scaling, offset and noise parameters. The proposed method is based on the marginalization of the posterior distribution, which reduces the dimensionality of the sampling problem. We provide analytical results for a broad class of problems and show that the method is suitable for a large number of applications. Subsequently, we demonstrate the benefit of the approach for various application examples from the field of systems biology. We report a substantial improvement up to 50 times in the effective sample size per unit of time, in particular when applied to multi-modal posterior problems. As the scheme is broadly applicable, it will facilitate Bayesian inference in different research fields.

## Introduction

Mathematical models are important tools for understanding and predicting the dynamics of many processes, such as signaling processing in biological systems [1–3], patient progression [4, 5] and epidemics [6, 7]. However, the parameters of mathematical models are in general unknown and need to be inferred from experimental data. This is an inherently challenging problem and complicated by the fact that, in addition to the dynamical properties of interest (e.g. rate constants and initial conditions), also characteristics of the measurement process may be unknown. In systems biology, most measurement techniques, including Western blotting [8], fluorescence microscopy [9] and mass spectrometry [10], are not fully quantitative but provide only relative information. Moreover, there is often an unknown offset and/or noise level [11]. Accordingly, unknown observation parameters, such as scaling factors but also offsets and noise levels, have to be estimated along with parameters of the mathematical models [12–14].

Bayesian inference is often used to determine unknown parameters [15–17]. A particularly common approach is to employ Markov chain Monte Carlo (MCMC) algorithms, such as (adaptive) Metropolis Hastings [18], Hamiltonian Monte Carlo methods [19, 20] and parallel tempering [21], to generate representative samples from the posterior distribution. Yet, with increasing number of unknown parameters, the application of MCMC algorithms becomes challenging [22]. This is a bottleneck that leaves sampling methods on the edge of computational feasibility. In principle, the challenge can be addressed by reducing the dimensionality of the sampling problem, e.g., by marginalizing over nuisance parameters (as e.g. demonstrated in cosmology [23]). However, there is no generic and broadly applicable framework.

In frequentist inference, a template for the reduction of the dimensionality of parameter estimation problems has been provided [14, 24, 25]. Here, hierarchical optimization approaches have been developed to determine the maximum likelihood estimate. These methods exploit that the observation parameters can be computed analytically for a given set of model parameters. It has been shown that this benefits the convergence of optimization methods and the computational efficiency, while providing the same results (see, e.g. [24]). Yet, these concepts cannot be directly translated to Bayesian inference as we are not interested in only optimal point estimates, but in (marginal) posterior distributions over parameters.

In this manuscript, we introduce a generic method for improving sampling efficiency by marginalizing over observation parameters. We provide analytical results for the marginalization over complex posterior distributions for a broad class of observation models. The marginalization yields a lower dimensional posterior for MCMC sampling. Samples of the original posterior can be obtained by subsequent sampling of the observation parameters conditioned on the remaining parameters. To illustrate the properties of the proposed approach, we benchmark its performance with a collection of published models, including models for which current available sampling strategies are computationally infeasible. We demonstrate that the proposed method achieves higher sampling efficiencies by reducing the auto-correlation of the samples and increasing the transition probabilities between posterior modes. Indeed, it turns a computationally infeasible sampling problems feasible, increasing the set of problems which can be tackled using Bayesian inference.

## Results

### Many model structures allow for analytical marginalization of parameters and sampling in lower dimensional space

To facilitate Bayesian inference for mathematical models with observation parameters, we developed and implemented a marginalization-based sampling approach (Figure 1). The approach allows for inferring the parameters of mathematical models, such as ordinary differential equation (ODEs) and partial differential equation models, from data via observation models with scaling, offset and noise parameters. For the case of a mathematical model with parameter *θ* and time- and parameter-dependent states *x*(*t, θ*), we consider for the case of a one-dimensional observable with additive Gaussian measurement noise the observation model

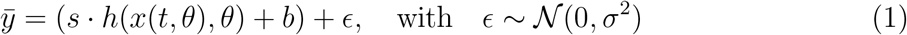

in which *h*(*x*, *θ*) is the observable map, *s* is the scaling factor, *b* is the offset and *σ*^2^ is the variance of the measurement noise. Following Bayes’ theorem, the posterior distribution is given by

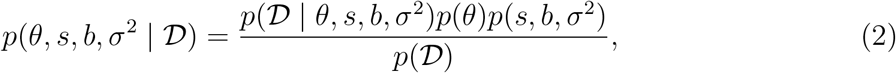

in which 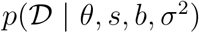 denotes the likelihood of the data 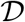, *p*(*θ, s, b, σ*^2^) = *p*(*θ*)*p*(*s, b, σ*^2^) denotes the prior distribution, and 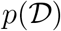 denotes the marginal probability of the data.

**Figure 1:**
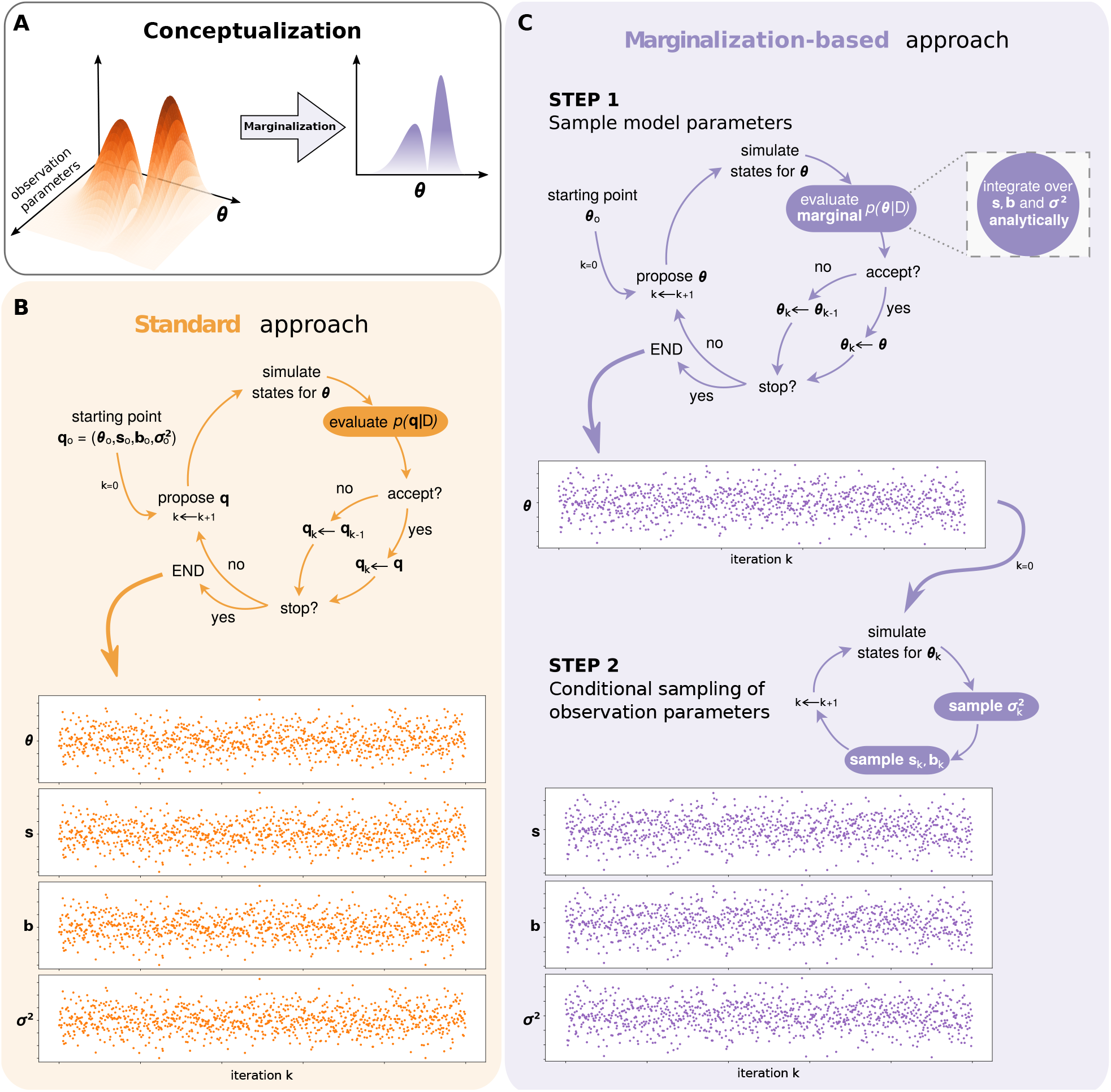
Standard and marginalization-based Markov chain Monte-Carlo sampling. (A) Illustration of the general marginalization concept. (B) Standard approach. (C) Marginalization-based approach depicting: (Step 1) the sequential integration of the observation parameters *s, b* and σ^2^ to evaluate 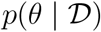, and (Step 2) the (optional) conditional sampling of the marginalized observation parameters.

The **standard approach** is to use MCMC methods to obtain representative samples from the joint posterior distribution for model parameters *θ* and observation parameters *s*, *b* and *σ*^2^ (2) for subsequent analysis (Figure 1B). All parameters are sampled jointly, disregarding their nature (Figure 1B), in particular note that the state *x*(*t*, *θ*) and the value of the observation map *h*(*x*(*t*, *θ*), *θ*) only depends on *θ* but not on *s, b* or *σ*^2^. This approach is often challenging and even infeasible for models with large datasets, since the number of observation parameters can easily exceed the number of model parameters (see e.g. [26, 27]).

To simplify the sampling process, we propose a **marginalization-based approach**, which exploits a decomposition of the sampling problem in two steps (Figure 1C). In Step 1, we consider the marginalization of the posterior distribution (2) with respect to the observation parameters *s, b* and *σ*^2^, yielding

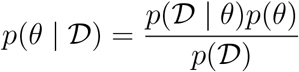

with 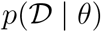 as the marginal likelihood given by

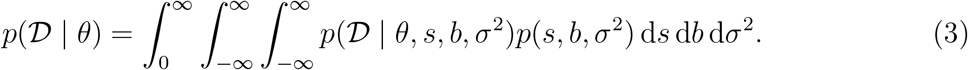

For various choices of noise models and prior distributions (in particular conjugate priors), this marginal likelihood can be computed in closed-form. This is for instance the case for the combination of additive Gaussian noise with a joint prior distribution for *s*, *b* and *σ*^2^,

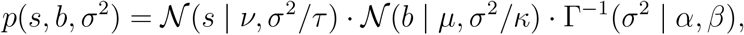

in which 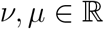 and 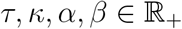 denote hyperparameters of the Normal-Inverse-Gamma-distributed joint prior, and Γ^-1^(·) the Inverse-Gamma function. Here, we obtain for observations *ȳ_i_* with *i* = 1,…, *n_t_* the closed-form expression for the marginal likelihood as

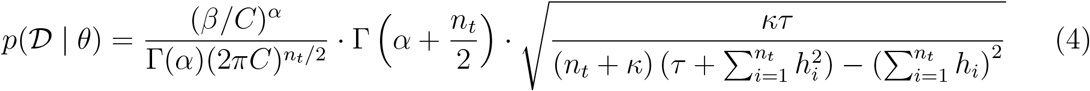

with *h_i_*:= *h*(*x*(*t_i_, θ*), *θ*) and parameter-dependent constant

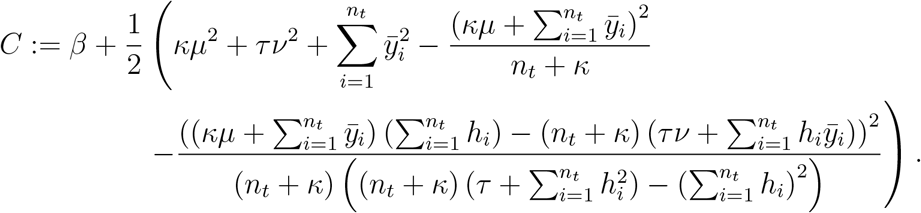

The combination of additive Gaussian noise and Normal-Inverse-Gamma prior is a common choice of conjugate distributions, which allow for an analytically tractable marginal likelihood. There are various other cases, including multiplicative Gaussian noise and even distributions with outliers. For the latter, Laplacian noise has shown to be more robust against measurement outliers [28]. Supplementary Tables S1–S2 summarize ten practically relevant cases for which we obtained closed-form expressions, and we are certain that many more are possible. For details on the derivation of all individual results (including two cases for Laplace distributed noise), we refer to the *Supplementary Material.*

Given the marginalized likelihood function 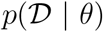 and the prior *p*(*θ*), the posterior distribution 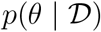 of the parameters of the mathematical model can be sampled using MCMC and related methods. The sampling can be performed in the space of *θ*, as the observation parameters are implicitly considered (Figure 1C).

The samples of model parameters *θ* from 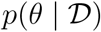 allow for the assessment of the model properties and its uncertainties. In this regard, there is no difference of sampling the marginalized posterior distribution 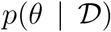 compared to projecting the full posterior distribution 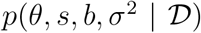 onto the *θ* component. However, tasks like the assessment and plotting of the model-data mismatch also require the posterior of the observation parameters. These can be obtained by sampling from the conditional distribution 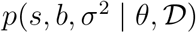. As the observation parameters only influence the observation model (1) and not the calculation of state *x*(*t,θ*) and observable map *h*(*x,θ*), the conditional distribution can be expressed in closed-form and sampled efficiently. For the aforementioned case, a matching sample of observation parameters for a given model parameter *θ* can be obtained by drawing from Gamma and Normal distributions:

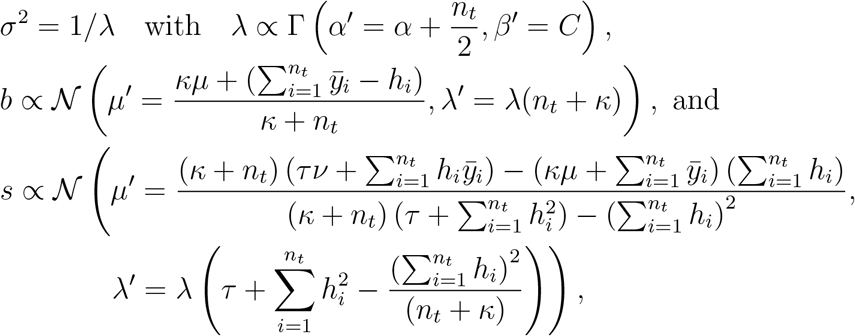

with *h_i_* and *C* being evaluated for model parameter *θ*. This conditional sampling can be proven to provide the same correlation structure as directly sampling the full posterior distribution. For details on the derivation of the conditional sampling for the observation parameters we refer to the *Supplementary Material*. As the conditional sampling can be performed independently and does not require model simulation, it is computationally efficient. For additional observation models see Supplementary Tables S1–S2.

In summary, a broad spectrum of sampling problems occurring in scientific disciplines, such as systems and computational biology, can be reformulated by performing an analytically tractable marginalization of their observation parameters. Sampling of this lower dimensional posterior distribution for the model parameters *θ* in combination with conditional sampling for the observation parameters allows the construction of samples from the full posterior distribution. Accordingly, the original sampling problem is decomposed in two sub-problems, of which the conditional sampling is optional.

### Marginalization-based approach yields same results at lower computational cost

To compare the performance for the standard and marginalization-based approach, we performed a range of studies using (i) a simple test problem and (ii) published models and datasets.

As a simple test problem we considered a model of a conversion reaction process, *A* ⇌ *B*. This process was considered in various other publications [28, 32] and can be described using a two-dimensional system of ODEs, with the concentrations of *A* and *B* as state variables. Here, we considered that the abundance of *B* is measured up to an unknown scaling, offset and noise level. Accordingly, the mathematical model possesses two model parameters: the forward rate *A* to *B*, *θ*_1_, and the backward rate *B* to *A*, *θ*_2_; and three observation parameters: the scaling *s*, the offset *b* and the noise variance *σ*^2^ (Table 1). A detailed description of the model is provided in the *Methods* section.

**Table 1:**
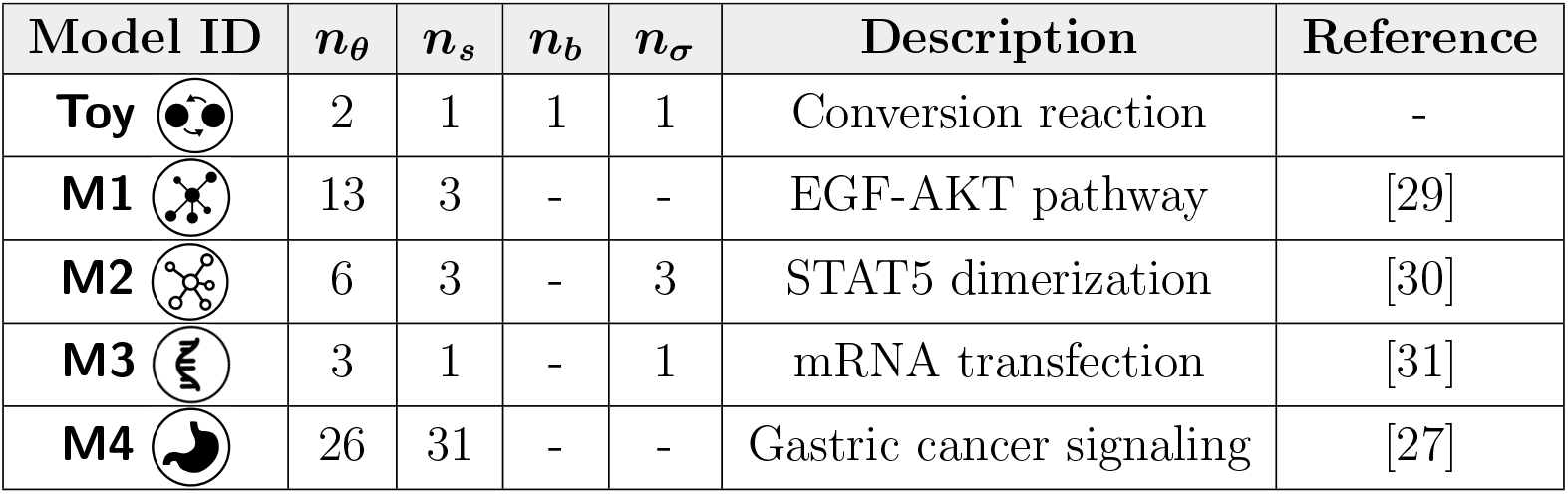
Key numbers and features of the considered toy and benchmark models. The number of unknown model parameters *n_θ_*, unknown scaling parameters *n_s_*, unknown offset parameters *n_b_* and unknown noise parameters *n_σ_*, which are effectively sampled, are reported.

| Model ID | $n_\theta$ | $n_s$ | $n_b$ | $n_\sigma$ | Description | Reference |
| --- | --- | --- | --- | --- | --- | --- |
| **Toy** | 2 | 1 | 1 | 1 | Conversion reaction | - |
| **M1** | 13 | 3 | - | - | EGF-AKT pathway | [29] |
| **M2** | 6 | 3 | - | 3 | STAT5 dimerization | [30] |
| **M3** | 3 | 1 | - | 1 | mRNA transfection | [31] |
| **M4** | 26 | 31 | - | - | Gastric cancer signaling | [27] |

In the first step, we used the model to assess the correctness of the analytical marginalized likelihood (4) by comparing its agreement with numerical integration of (3). The results show a perfect match for a range of different parameter values (Figure 2A). Yet, the evaluation of the analytical marginalized likelihood was five orders of magnitude faster than the numerical integration (Figure 2B), which highlights the importance of the analytical derivations. In the second step, we performed 100 independent MCMC sampling runs for the standard and marginalization-based approach. The runs employed a state-of-the-art adaptive Metropolis Hasting method [18]. We found a superior performance of the marginalization-based approach, as the observed effective sample size per unit of time was twice as high as for the standard approach (Figure 2C). This indicates that the marginalization-based approach facilitates already for simple problems the mixing of the MCMC chains and, hence, provides a more efficient exploration of the posterior. Moreover, the model fit for the best sample found (i.e. maximizing the posterior) coincided for both approaches (Figure 2D) as well as the marginal distributions for the model parameters *θ*_1_ and *θ*_2_ (Figure 2E–F), and the conditionally sampled observation parameters (Figure 2G–I).

**Figure 2:**
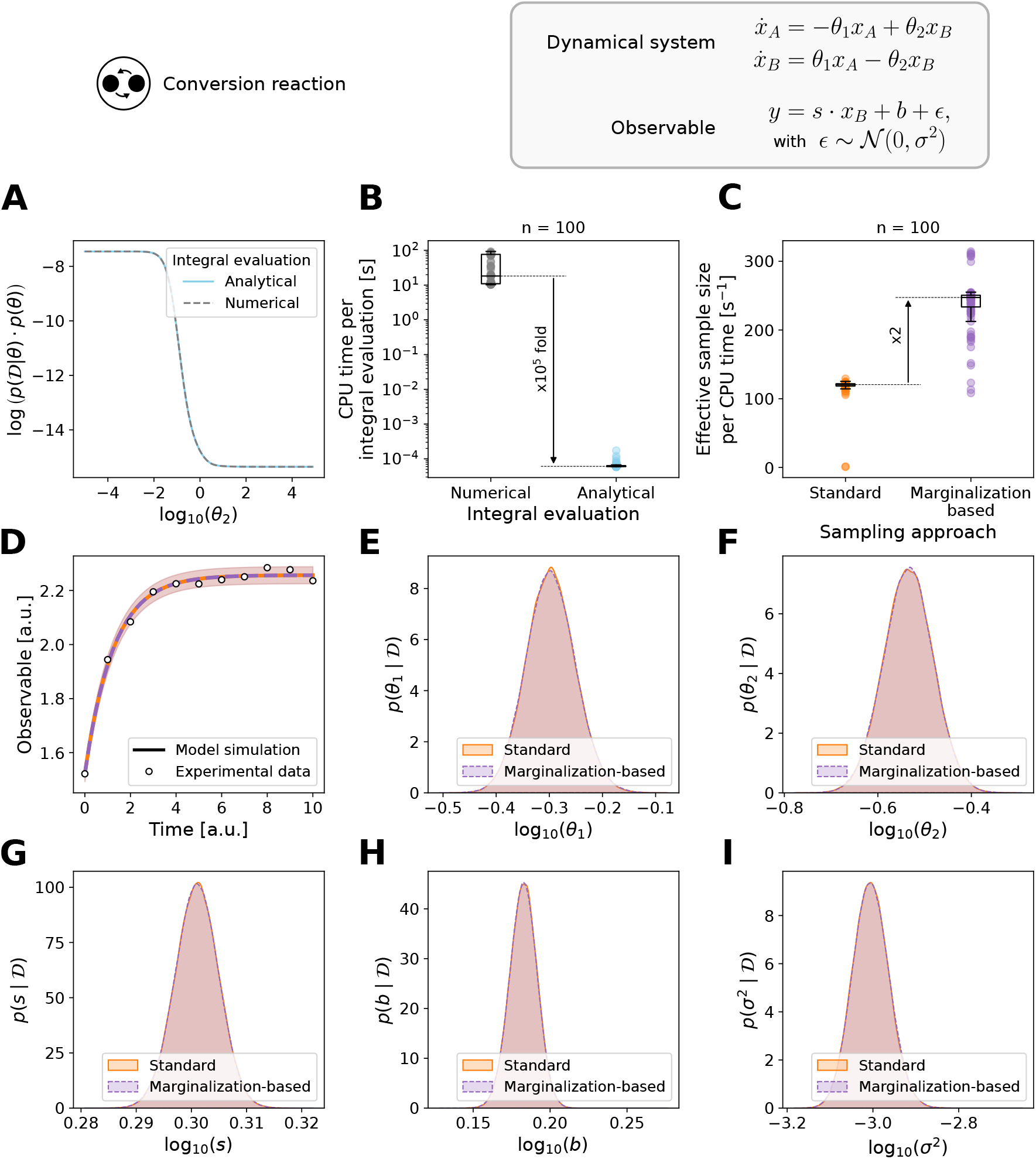
Evaluation of the standard and marginalization-based approach for the toy model. (A) Comparison of analytical vs. numerical integration. (B) Time comparison of analytical vs. numerical integration. (C) Effective sample size per unit of time for 100 independent runs. (D) Model fit of the best sample found during sampling from the standard (orange) and marginalization-based (purple) approach. (E–I) Parameter marginal posterior distributions computed using a kernel density estimate for the model parameters (E) *θ*_1_ and (F) *θ*_2_, and the conditionally sampled observation parameters: (G) scaling factor *s*, (H) offset *b*, and (I) noise variance *σ*^2^.

Following the promising results for the test problem, we evaluated the performance of the proposed marginalization-based approach for three already published models and datasets (Table 1 and *Methods* section). The models M1 to M3 describe cellular processes: (M1) EGF-induced AKT signalling; (M2) phosphorylation-dependent STAT5 dimerization; and (M3) mRNA transfection. The numbers of model and observation parameters differ, and so do the observation functions. Accordingly, different closed-form expressions for the marginalized likelihood function are used (Supplementary Tables S1 – S2). More importantly, the full posterior distributions exhibit different characteristics, ranging for instance from uni-to bi-modal.

For the considered application problems, the marginalization of the observation parameters reduced the dimensionality of the sampling problems by up to 50% (Figure 3A). To evaluate the impact of this reduction on the sampling efficiency, we performed 50 independent MCMC sampling runs using the parallel tempering algorithm with 10 temperatures [21] after assessing the correctness of the analytical marginalized likelihood for models M1–M3 (Supplementary Figure S9). All the runs were initialized at the local optima found during multi-start optimization [12], and run for 10^6^ iterations. Further details are provided in the *Methods*section. The high number of iterations allowed all MCMC runs of the standard and marginalized problem to converge according to the Geweke test [33]. Yet, the marginalization-based approach achieved a higher effective sample size per unit of computation time than the standard approach (Figure 3B). The improvement was problem dependent and ranged from 2 (M1 and M2) to nearly 50 (M3) times higher efficiency in the marginalization-based approach. As the computation time was similar, the core reasons for this is a reduction in the auto-correlation length (Figure 3C). The model fits for the best sample found were identical for both approaches (Figure 3D) as well as the parameter marginal distributions (Supplementary Figures S1, S3 and S5).

**Figure 3:**
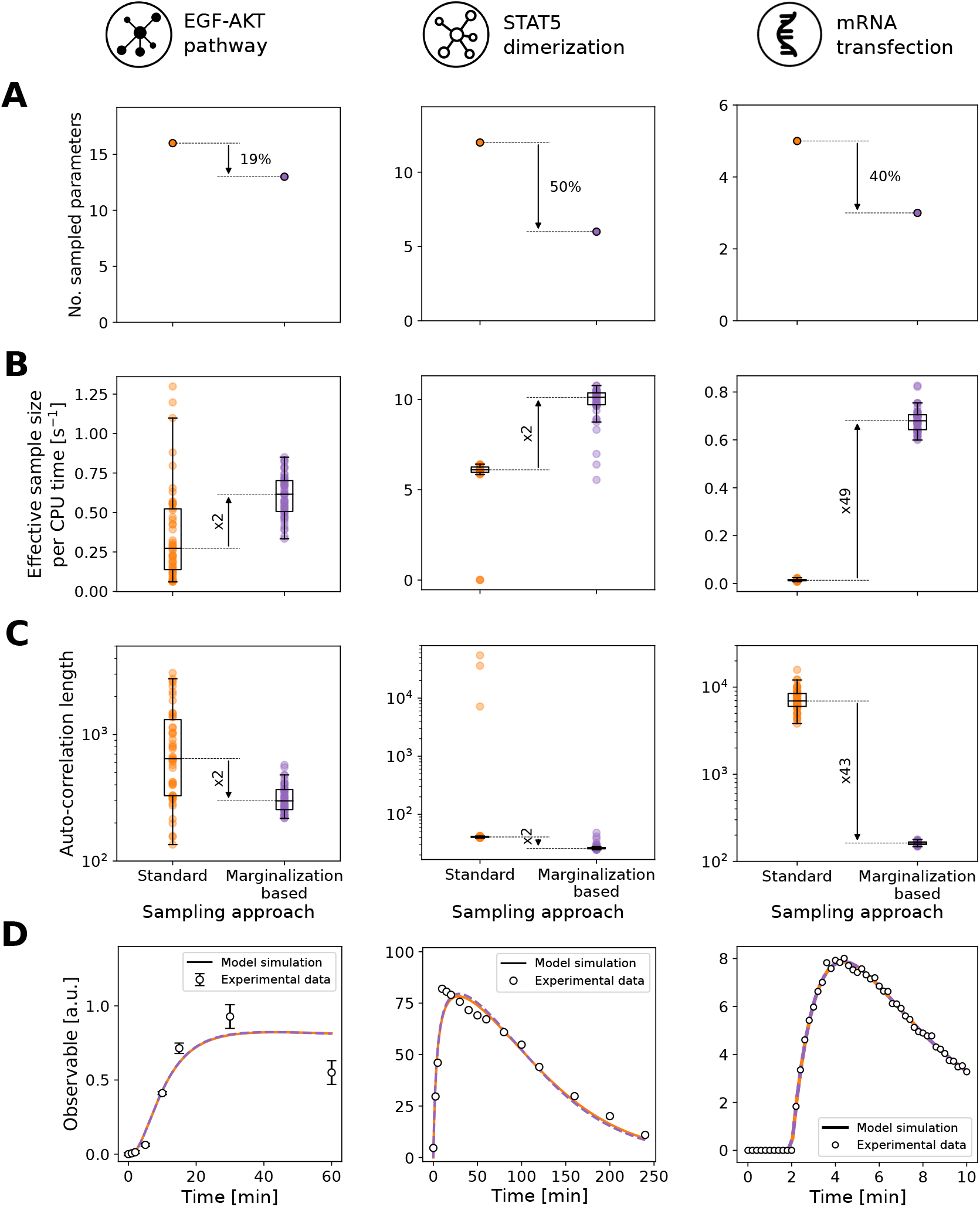
Evaluation of the standard and marginalization-based approach for the benchmark models. Models M1–M3 are shown from left to right. (A) Number of sampled parameters. (B) Effective sample size per unit of time. (C) Auto-correlation length. (D) Model fit of the best sample found during sampling. A subset of the experimental data is shown for M1 and M2. Complete datasets are depicted in Supplementary Figures S2 and S4.

In summary, test and application problems demonstrates the acceleration potential of the marginalization-based approach. The improvement was problem specific, with no clear dependence on the degree of dimensionality reduction, but in all cases substantial.

### Marginalization-based approach improves transition rates between posterior modes

To understand for which problems the marginalization-based approach is expected to achieve a large acceleration, we considered the model M3. The posterior distribution for M3 is bi-modal and a simple explanation for the acceleration would have been that the bimodality is eliminated. Yet, this is not the case as the bimodality is related to a symmetry in model parameters. Numerical simulations as well as analytical results reveal that the observable trajectory remains unchanged when the mRNA and protein degradation rates are interchanged. As long as the optimal point is not located on the line of equal degradation rates, standard and marginalized posterior are bimodal.

We hypothesized that the large efficiency improvement is related to a lower minimum energy path for the transitions in the marginalized posterior. To assess this, we computed the minimum energy paths [34] for the standard (Figure 4A,B) and marginalized posterior (Figure 4C,D) (see details in the *Methods* section). To our surprise, the minimum energy path is almost identical for both approaches (Figure 4E). Hence, there is at least no difference in the minimum energy path.

**Figure 4:**
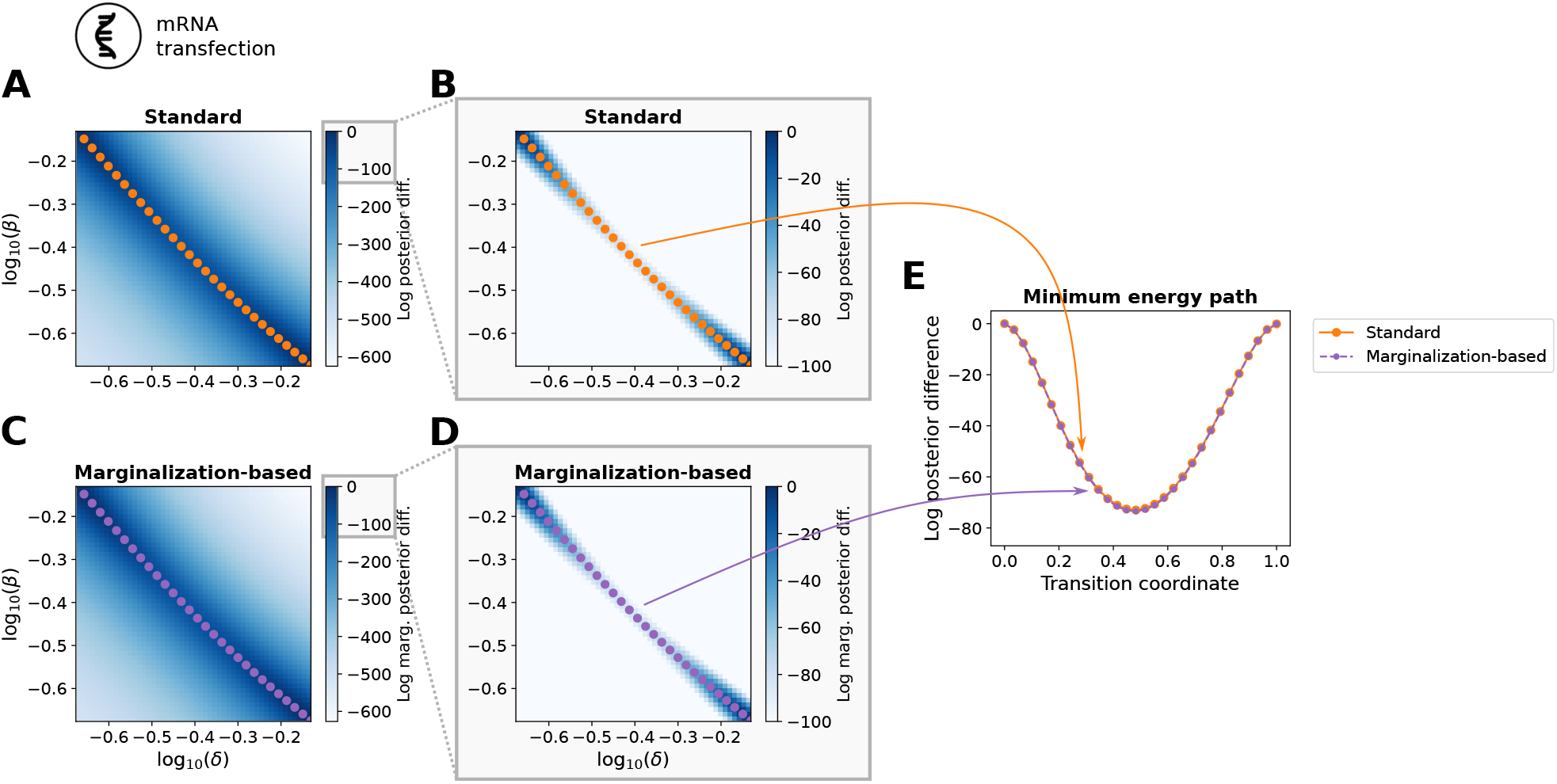
Comparison of the minimum energy path for model M3. Landscape of the optimized (A,B) posterior and (C,D) marginalized posterior for different fixed values of the model parameters *β* and *δ*. The difference with respect to the maximal posterior value is depicted. (E) Transition coordinates for the minimum energy path.

In order to understand the improvement observed for runs of adaptive parallel tempering methods, we performed 10 runs of a single-chain adaptive Metropolis algorithm [18] with 10^6^ iterations. This simplified the interpretation as it excludes the possibility of chain swaps. Yet, we found that for the given number of iterations this single-chain algorithm does essentially not transition between the modes (see *T* = 1 in Figure 5A). To assess the relative complexity of the sampling problem for standard and marginalization-based approach, we repeated the evaluation for the tempered posterior. We found that the marginalization-based approach allows already at lower temperatures for transitions between the modes unlike the standard sampling approach (Figure 5A and Supplementary Figures S7–S8). For temperatures such as *T* = 16, the standard approach showed an average number of only 5 transitions between the modes with many runs only sampling from a single mode (Figure 5B,C), while for the marginalization-based approach on average 1.6 × 10^4^ transitions occurred (Figure 5D,E). As the minimum barrier energy is conserved also for higher temperatures (Supplementary Figure S6), this increase in the transition rate by four orders of magnitude for the same algorithm implies a lower overall complexity of the marginalization-based sampling problem.

**Figure 5:**
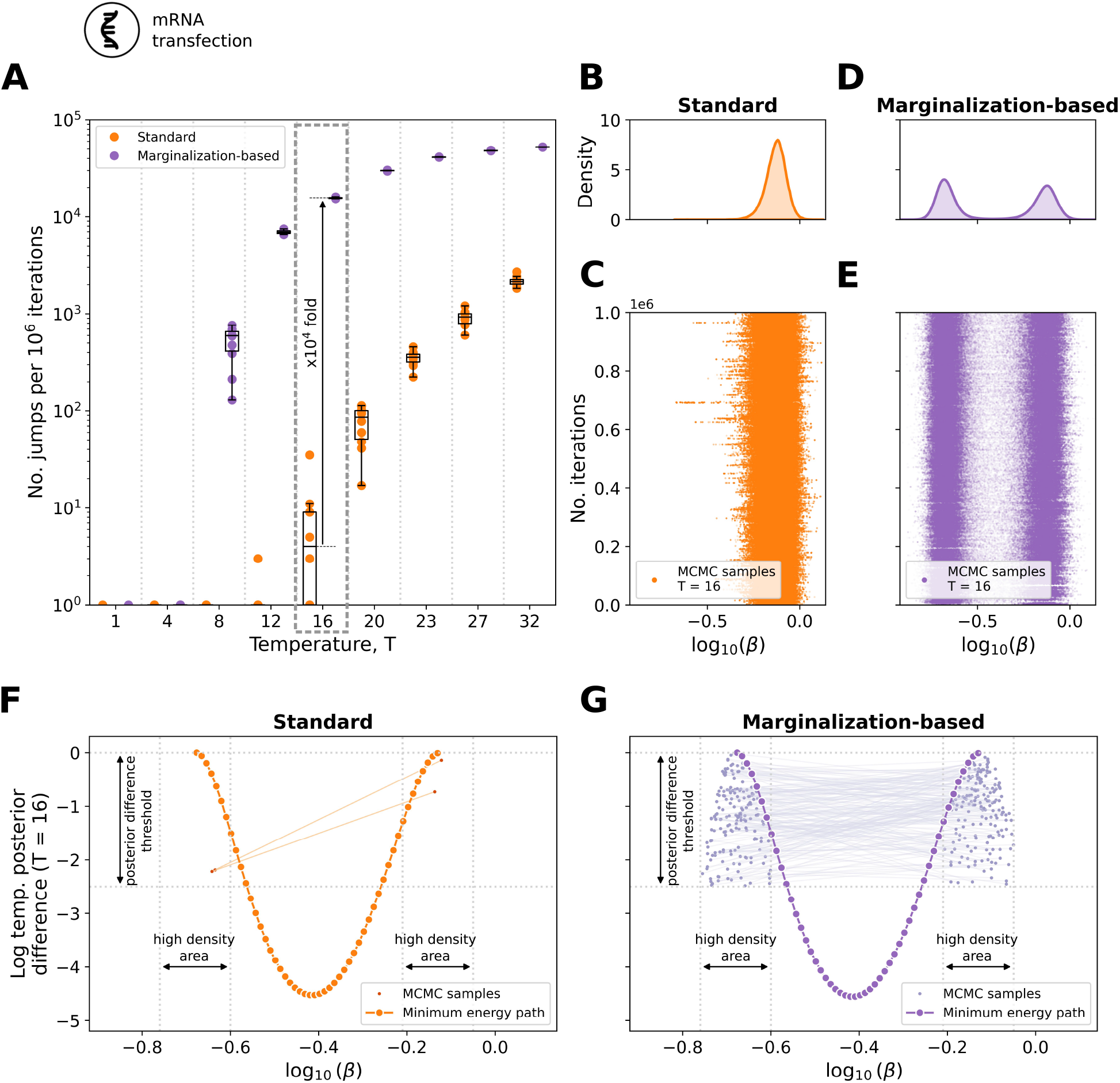
Quantification of the transitions between the posterior modes for different temperatures *T* for model M3. (A) Number of transitions per 10^6^ iterations for a range of temperatures for the standard (orange) and marginalization-based (purple) approach. A total of 10 chains per temperature value are depicted. (B,D) Marginal distribution computed using a kernel density estimate and (C,E) parameter trace for the model parameter *β* of a representative chain obtained with the (B,C) standard and (D,E) marginalization-based approach for *T* = 16. (F,G) Direct transitions between the posterior modes of a representative chain along with the minimum energy path obtained with the (F) standard and (G) marginalization-based approach for *T* = 16.

As the increased transition rate is not caused by an altered energy path, we studied the transition paths. This revealed that the employed single-chain algorithm facilitates jumps over the valley in the objective function (Figure 5F,G), meaning that it transitions between high-probability regions around the local optima. These direct transitions appear at a high rate for the marginalization-based approach (Figure 5G), while they rarely happen for the standard approach (Figure 5F). For the latter, most transitions are along low-energy paths with posterior probabilities dropping below the minimum energy path. Accordingly, the transition behaviour is for the marginalization-based approach more efficient than for the standard approach.

In summary, the in-depth study of the mRNA transfection model (M3) showed that the marginalization-based approach can achieve substantial accelerations as the structure of the sampling problem is simplified, e.g. by facilitating transitions between modes. The improvements are related to the interplay of sampling approach and problem geometry. In particular for challenging (e.g. multi-modal) problems a much greater improvement could be observed.

### Marginalization-based approach enables Bayesian inference for large models

As the marginalization-based approach appeared beneficial for challenging problems, we assessed in a next step whether it enables Bayesian inference for problems for which standard approaches did not provide reproducible results in a reasonable time-frame. Specifically, we considered an ODE model for signal transduction in gastric cancer cells (cell line MKN1) that was developed to unravel response and resistance markers [27]. This model possesses in total 57 unknown parameters, of which 26 are model parameters and 31 are observation parameters (Table 1, M4).

The application of the marginalization-based approach resulted in a reduction of the dimensionality of the sampling problem by over 50% (Figure 6A). For the 26 model parameters which remain to be sampled, we compared the marginal likelihoods as computed using the previously derived analytical formulas and numerical integration (Figure 6B). The agreement of the results (Pearson correlation *r* ≈ 1.0) confirmed the correctness of our analytical integration.

**Figure 6:**
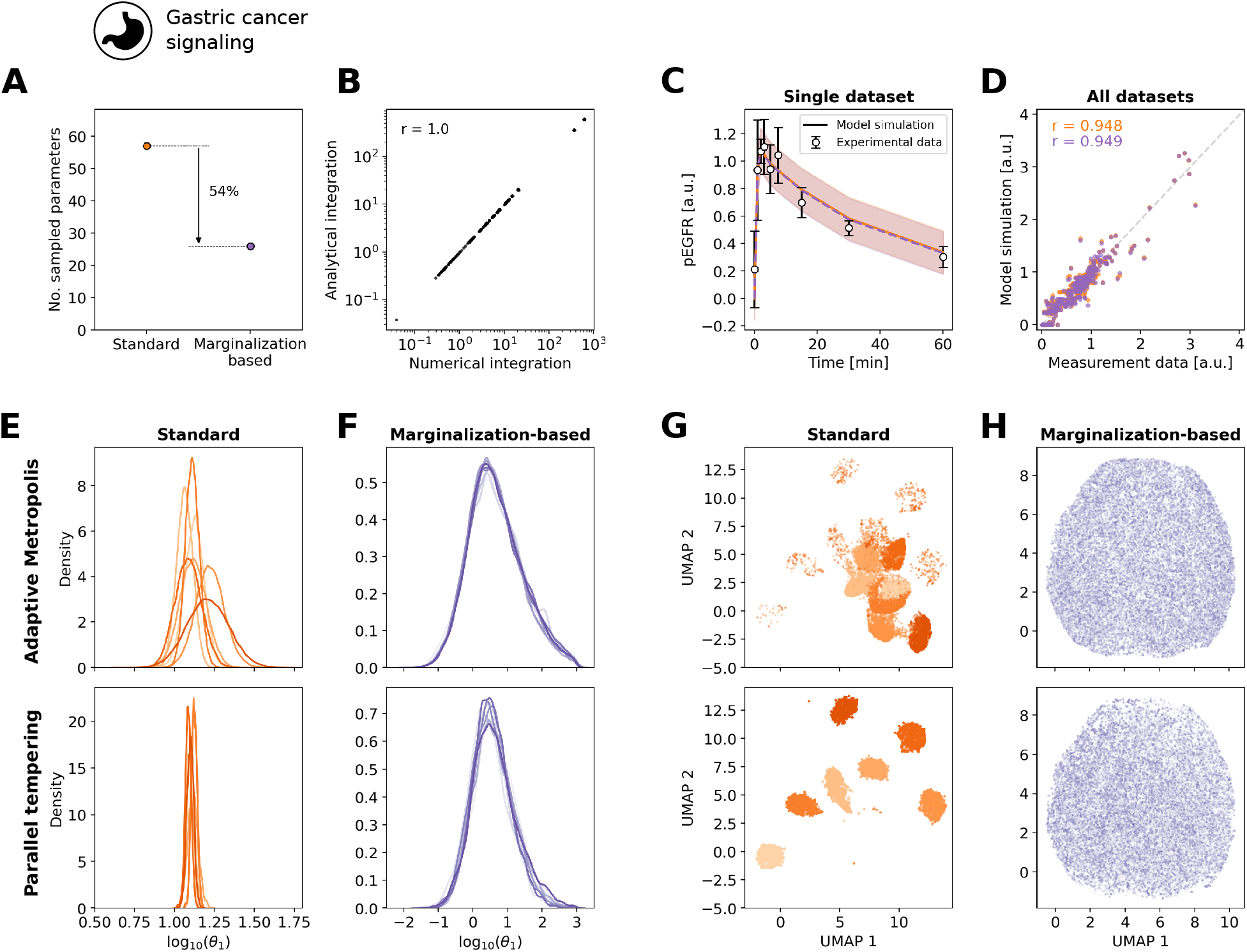
Convergence of the marginalization-based approach for model M4. (A) Number of sampled parameters. (B) Scatter plot for the agreement of analytical and numerical integration. (C,D) Model fit of the best sample found during sampling for, (C) a subset of the experimental data and (D) the complete dataset in form of a scatter plot, the standard (orange) and marginalization-based approach (purple). (E–H) Results from adaptive Metropolis (top) and parallel tempering (bottom) are shown. (E,F) Parameter marginal posterior distribution obtained using the (E) standard and (F) marginalization-based approach computed using a kernel density estimate for model parameter *θ*_1_. (G,H) Dimensionality reduction for all samples from all runs for the (G) standard and (H) marginalization-based approach using the UMAP representation. Different shades correspond to individual runs. The UMAPs were constructed using the Python package `umap` [35].

To determine the parameters of the model, we performed sampling using standard and marginalization-based approach. The adaptive Metropolis-Hastings algorithm [18] and the adaptive parallel tempering algorithm [21] employed in the previous sections were run 10 times with different starting points and random seeds for 10^6^ iterations for the adaptive Metropolis-Hastings and 10^5^ iterations for the adaptive parallel tempering algorithm. The maximum a posteriori estimates observed in the different runs provided similar fits (Figure 6C,D). In contrast, the marginal distributions of the model parameters differed, with the marginalization-based approach mostly providing broader parameter distributions than the standard approach (Figure 6E,F). The assessment of the reproducibility of the marginal distributions revealed a high variability between different runs performed using the standard approach (Figure 6E and Supplementary Figure S10). On the contrary, for the marginalization-based approach a good agreement between runs was observed (Figure 6F and Supplementary Figure S11), indicating reproducibility. To verify that the behavior observed for the individual parameters is maintained in the full parameter space, we analyzed the overall agreement of all parameter samples across all runs for the standard and marginalization-based approach by visualizing the samples using the uniform manifold approximation and projection (UMAP) representation [35]. We found that the individual runs of the standard approach represent individual clusters in the UMAP (Figure 6G), while the individual runs of the marginalization-based approach were indistinguishable (Figure 6H). This revealed that: (i) in the marginalization-based approach all the individual runs sample from the same distribution, and (ii) the standard approach failed for both algorithms considered here.

The study of the model of signal processing in gastric cancer cells revealed that marginalization-based approach allows for reproducible sampling in problems, where the standard approach failed. While for the marginalization-based approach all runs provided consistent results, the standard approach failed to converge within an average CPU time of 150 hours rendering its application impracticable. Furthermore, our study provides improved estimates for the parameters (Supplementary Figure S12) of important processes of a drug used in clinical practice.

In summary, the application of our marginalization-based approach to Bayesian inference for models with relative measurement data shows consistently that our approach yields the same marginal distributions for the parameters as the standard approach, while being highly more efficient in exploring the parameter space and enabling Bayesian inference of larger models, which was not possible before with the standard approach.

## Discussion

Bayesian inference for models of biological processes requires the consideration of parameters of the dynamical systems as well as the measurement process. The unknown scaling factors, offsets and noise levels often resemble large fraction of the overall parameters [12]. This complicates sampling and can render the generation of representative samples practically infeasible. Here, we address this challenge by introducing a framework which employs (analytical) marginalization. This approach allows for the construction of a sample from the full posterior by (i) sampling a marginalized posterior for the parameters of the dynamical systems and (ii) conditional sampling of the observation parameters.

We evaluated the performance of our marginalization-based approach and compared it to the standard approach for four published models, with differences in their complexity. This revealed an increased effective sample size per unit of time, and increased transition probabilities between posterior modes. The marginalization-based approach was for all considered problems more efficient than the standard approach, but – more importantly – it also enabled the assessment of the posterior distribution for larger models for which the standard approach failed to converge in the considered time-frame. Interestingly, there was no strong relation between the reduction of the problem dimensionality and the improvement in efficiency. This is consistent with previous finding for hierarchical optimization [25]. Based on our observations we expect the sampling behavior to benefit substantially even from the removal of a small number of parameters, as (i) the likelihood value is often very sensitive to them, which produces narrow rims in the posterior distribution, and as (ii) the removal of a small number of parameters can result in a substantially increased probability to jump between modes. The latter was observed for the model of mRNA transfection.

The approach presented here is not limited to relative measurement data, but also applicable to absolute measurements. As for these, the noise parameters would still have to be inferred (Supplementary Tables S1 and S2). We provide the detailed derivation in the *Supplementary Material*. Accordingly, our approach can be used for combinations of relative and absolute data. Also, it is applicable to different measurement process functions and noise models to the ones considered here. We hypothesize that also an extension to correlated noise is possible, but this remains to be assessed.

The choice of conjugate priors for the marginalized parameters eased the analytical derivation of the marginal posterior. This implies in our case that observable and noise parameters are not independent under the prior. Mostly, this is not a problem since both parameters are related to the measurement process. However, in some cases, there might be known to be independent, therefore other prior distribution assumptions must be considered. It should be noted that the concept of marginalization is not restricted to integrals that are analytically solvable, but also numerical integration schemes can be considered. However, this would increase the required computation time (as observed in Figure 2B), but very likely the improved mixing properties would be maintained.

The proposed method was beneficial in combination with adaptive Metropolis-Hastings and adaptive parallel tempering algorithms. We expect that the same will hold true for sampling algorithms exploiting gradient information, such as Hamilton Monte Carlo sampling [19, 20]. As the marginal likelihood is differentiable, merely the derivation and implementation of the gradient is required. The usage of methods which exploit the Riemann geometry of the parameter space of statistical models, e.g., Metropolis-adjusted Langevin algorithm [36], might be slightly more involved. This requires the derivation of the marginalized Fisher information matrix. While we assume that this can be derived in closed-form or at least be accurately approximated, the corresponding results are not yet available. Alternatively, automatic differentiation could be employed to obtain gradients [37].

In this study, we focused on the assessment of parameter uncertainties for ODE models. Yet, as the marginalization-based approach provides a complete parameter sample, it facilitates also the evaluation of prediction uncertainties [16]. Accordingly, we expect that it might contribute to resolving reliability problems of Bayes prediction uncertainty analysis encountered in recent studies [38]. Furthermore, the proposed approach is not limited to ODEs, but directly applicable for other deterministic models, e.g. partial differential equations. As well, the idea might be incorporated in likelihood-free inference schemes used for stochastic and multi-scale models [39, 40]. Among other things, it might be used in exact Approximate Bayesian Computation schemes [41] by reformulating the acceptance probability.

In summary, the marginalization-based approach provides a new tool for Bayesian inference for models with observation-related parameters. It substantially benefits the efficiency of sampling-based approaches, and renders the generation of representative posterior samples for large models possible. As it is agnostic to the structure of the underlying dynamical model, it is widely applicable to mathematical models from different research fields, such as engineering, physics and ecology.

## Methods

### Mechanistic modeling of biological systems

We consider models based on ODEs of the form

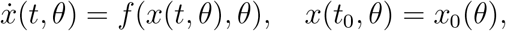

in which the vector field 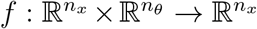 determines the temporal evolution of the states 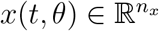. The unknown model parameters, which are estimated from the measurements, are denoted by 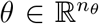. Usually, *θ* includes reaction rate constants and initial amounts of species. Here, *n_x_* is the total number of modeled species, and *n_θ_* the total number of model parameters. The states *x*(*t*, *θ*) and model parameters θ are linked to the observables via the observation map 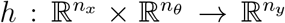, where *n_y_* is the total number of observables. The observables are the measured properties of the model. Most measurement techniques only provide relative information about the absolute concentrations of interest [8, 9] and, frequently, measurements are noise corrupted. Hence, to obtain the measurements *ˌ* (i) the model observables must be rescaled by introducing scaling factors and offsets, and (ii) the model also must capture experimental errors by defining a noise model. Most commonly, independent and additive Gaussian distributed noise models are assumed

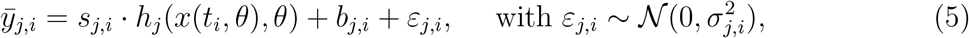

with observable index *j*, time index *i*, scaling factors 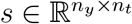, offsets 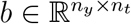, and noise parameters 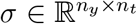. Here, *n_t_* denotes the total number of time points. These parameters are often unknown and, therefore, also need to be estimated along with the unknown model parameters. Other usual noise assumptions include log-normal distributed noise models [11] and Laplace distributed noise models [28]. In this study, we focus on the case of additive Gaussian noise (5), but implementations for log-normal and Laplace distributed noise models are provided in Supplementary Tables S1–S2 and *Supplementary Material*.

We denoted the group of all measurements as 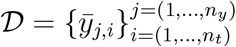

### Benchmark models

For the evaluation of the marginalization-based approach, we employed in total five models (one toy model and four published M1–M4) and their corresponding datasets (Table 1).

#### Toy: Model of a conversion reaction 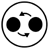

The conversion reaction model was introduced in [28] and describes a reversible chemical reaction, which converts a biochemical species *A* to a species *B* with rate *θ*_1_, and *B* to *A* with rate *θ*_2_ (Figure 2). We modified the observation model to include scaling and offsets. For the evaluation of the proposed method, we generated one artificial dataset which is depicted in Figure 2D. For details on the model structure and synthetic data generation we refer to the Supplementary Material.

#### M1: Model of EGF-dependent AKT pathway 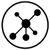

The model of EGF-dependent AKT pathway has been introduced in [29] and possesses in total 16 unknown parameters: 13 model parameters and 3 scaling factors (Table 1, M1). The available experimental data are a total of 144 data points under 6 different experimental conditions for 3 observables. For each data point, the corresponding variance of the measurement noise is provided, therefore it does not need to be estimated. The complete dataset is depicted in Supplementary Figure S2.

#### M2: Model of STAT5 dimerization 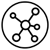

The model of STAT5 dimerization has been introduced in [30] and possesses in total 9 unknown parameters: 6 model parameters and 3 noise parameters. To this model, we have added 3 scaling factors (Table 1, M2), one per observable, for the sake of testing the proposed method. The available experimental data are a total of 48 data points for 3 observables. The complete dataset is depicted in Supplementary Figure S4.

#### M3: Model of mRNA transfection 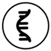

The model for mRNA transfection has been introduced in [31] and possesses in total 5 unknown parameters: 3 model parameters, 1 scaling factor, and 1 noise parameter (Table 1, M3). The complete dataset is depicted in Figure 3D. For further details of the model structure we refer to the Supplementary Material.

#### M4: Model of gastric cancer signaling 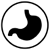

The model for gastric cancer signalling has been introduced in [27]. Here, we considered the Cetuximab responder cell line MKN1. The available experimental data for the responder cell line were a total of 303 data points under 106 different experimental conditions for 31 observables. For each data point, the corresponding variance of the measurement noise was provided, therefore it did not need to be estimated.

### Parameter optimization

To determine the maximum a posteriori (MAP) estimates, we minimized the negative log-posterior function. This minimization was performed using multi-start local optimization, an approach which was previously shown to be reliable [12, 42]. For local optimization, we used the trust-region optimizer `fides` [43]. Parameters were log_10_-transformed to improve numerical properties [42, 44, 45]. We generated 100 starting points for local optimization, except for model M4 for which we used 500 starting points.

### Bayesian parameter inference

To perform Bayesian parameter inference, we used MCMC sampling following the pipeline presented in [46]. The MAP estimates were used to initialize the MCMC chains [46]: the full optimal vector (*θ, s, b, σ*^2^)* to initialize the standard approach runs, while for the marginalization-based approach runs the corresponding subset *θ*\* from (*θ, s, b, σ*^2^)* was used.

The parameter posterior distribution was sampled using the adaptive Metropolis [18] and parallel tempering [47, 48] algorithms implemented in the Python toolbox pyPESTO [49]. For the parallel tempering algorithm, we used 10 chains initialized at the 10 best local optima found during multi-start optimization for both approaches.

Convergence after burn-in was assessed using the Geweke test [33] and auto-correlation length using Sokal’s adaptive truncated periodogram-estimator [50], both also available under pyPESTO. The effective sample size is given by

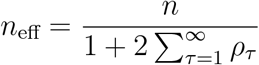

where *n* is the number of samples remaining after discarding burn-in period, and *ρ_τ_* is the estimated auto-correlation at lag *τ*.

For all models, the prior hyperparameters for both sampling approaches were the same as used for optimization.

### Tempering scheme for the posterior analysis

The posterior for standard and marginalization-based approach were tempered to assess transition characteristics (Figure 5). We used the tempered posteriors

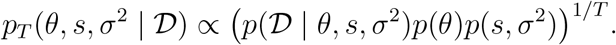

and

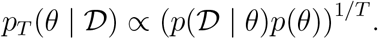

with temperature *T*.

### Implementation and data availability

Models M1, M2 and M4 were taken from the PEtab benchmark collection [51] which is based on [44] and available at https://github.com/Benchmarking-Initiative/Benchmark-Models-PEtab. As model M3 is analytically solvable, we implemented the solution in Python code. For ODE integration (models M1, M2 and M4) we used the Python toolbox AMICI [52]. For optimization and sampling, we used the Python toolbox pyPESTO [49]. pyPESTO already offers an interface to the fides optimizer [43]. For the UMAP visualizations and the minimum energy path calculation, we used respectively the Python packages umap [35] and mep https://github.com/chc273/mep.

All code and models used in this study are available from the Zenodo database at https://doi.org/10.5281/zenodo.7199473.

## Acknowledgements

This work was supported by the German Federal Ministry of Education and Research (Grant no. 031L0159C; J.H.), the University of Bonn (via the Schlegel Professorship; J.H.), the Deutsche Forschungsgemeinschaft (DFG, German Research Foundation) under Germany’s Excellence Strategy EXC 2047/1 - 390685813 (E.R., M.F., J.H); EXC 2151 - 390873048 (E.R., J.H.); TRR 333/1 - 450149205 (E.R., J.H.); SFB 1454 - 432325352 (M.F.); and 443187771 (J.H.).

## Author contributions

Conceptualization: J.H., E.R.; Methodology: J.H., E.R., M.F.; Software: E.R.; Formal analysis: E.R.; Investigation E.R., M.F.; Data curation: E.R.; Writing – original draft: J.H., E.R.; Writing – review and editing: all authors; Visualization: E.R.; Supervision: J.H., E.R.; Funding acquisition: J.H.

## Supplementary Figures and Tables

**Figure S1:**
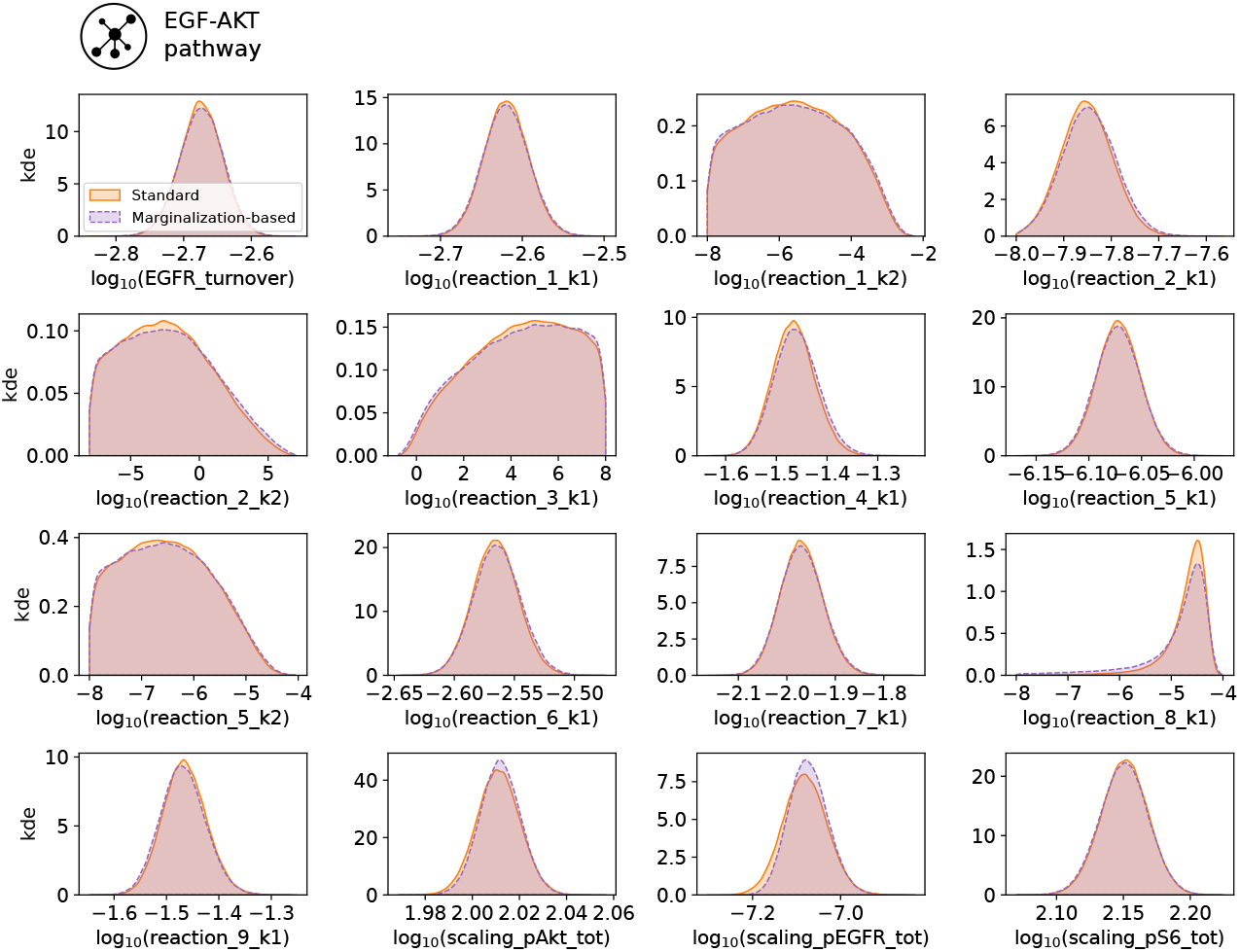
Parameter marginal posterior distributions computed using a kernel density estimate for model M1. The marginalized parameters, which are conditionally sampled, correspond to those denoted with *scaling_*\*.

**Figure S2:**
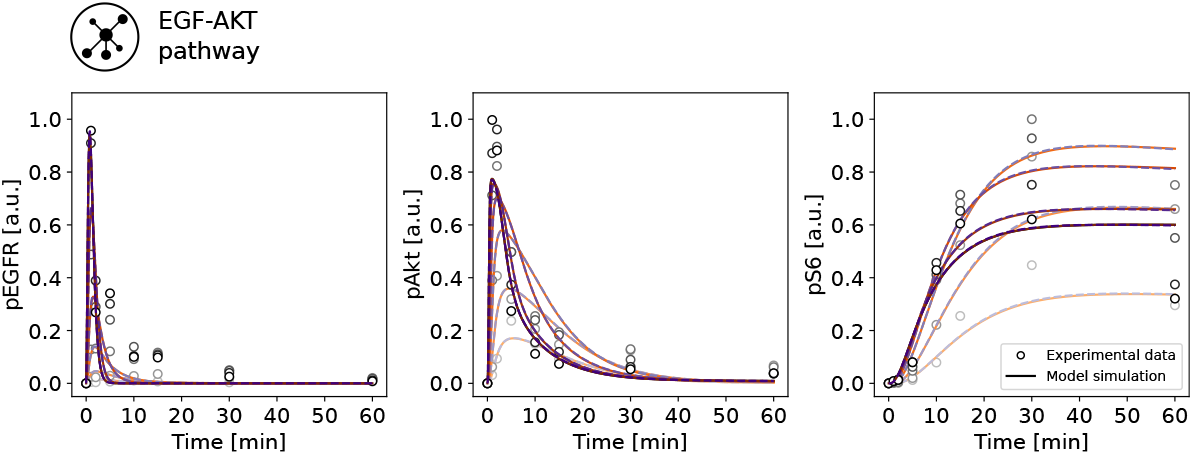
Complete dataset and model fit for model M1. Model simulation of the best sample found for the standard approach is depicted in orange and for the marginalization-based approach in purple. Different shades indicate different experimental conditions.

**Figure S3:**
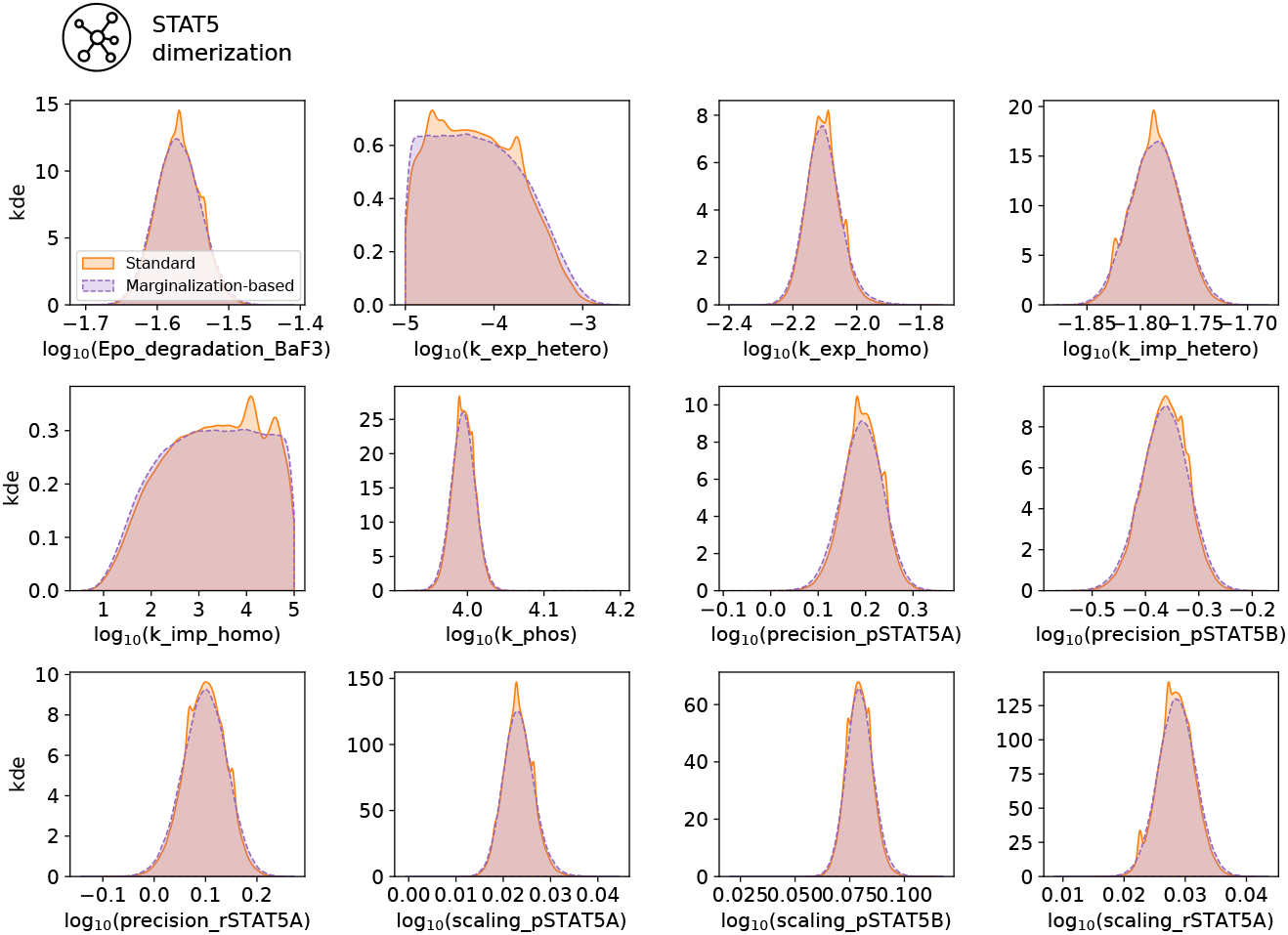
Parameter marginal posterior distributions computed using a kernel density estimate for model M2. The marginalized parameters, which are conditionally sampled, correspond to those denoted with *scaling_*\* and *precision_\**.

**Figure S4:**
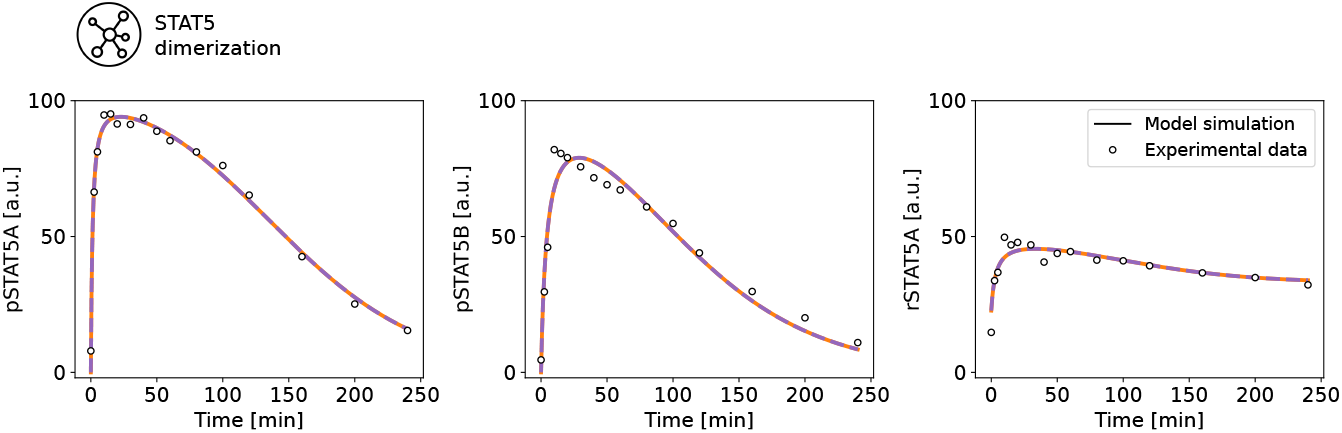
Complete dataset and model fit for model M2. Model simulation of the best sample found for the standard approach is depicted in orange and for the marginalization-based approach in purple.

**Figure S5:**
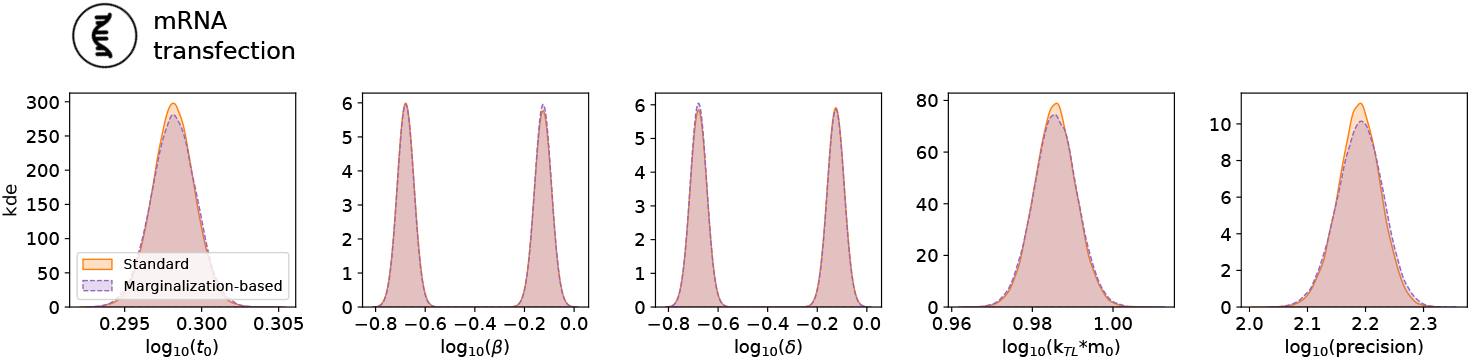
Parameter marginal posterior distributions computed using a kernel density estimate for model M3. The marginalized parameters, which are conditionally sampled, correspond to *k_TL_* * *m*_0_ and *precision*.

**Figure S6:**
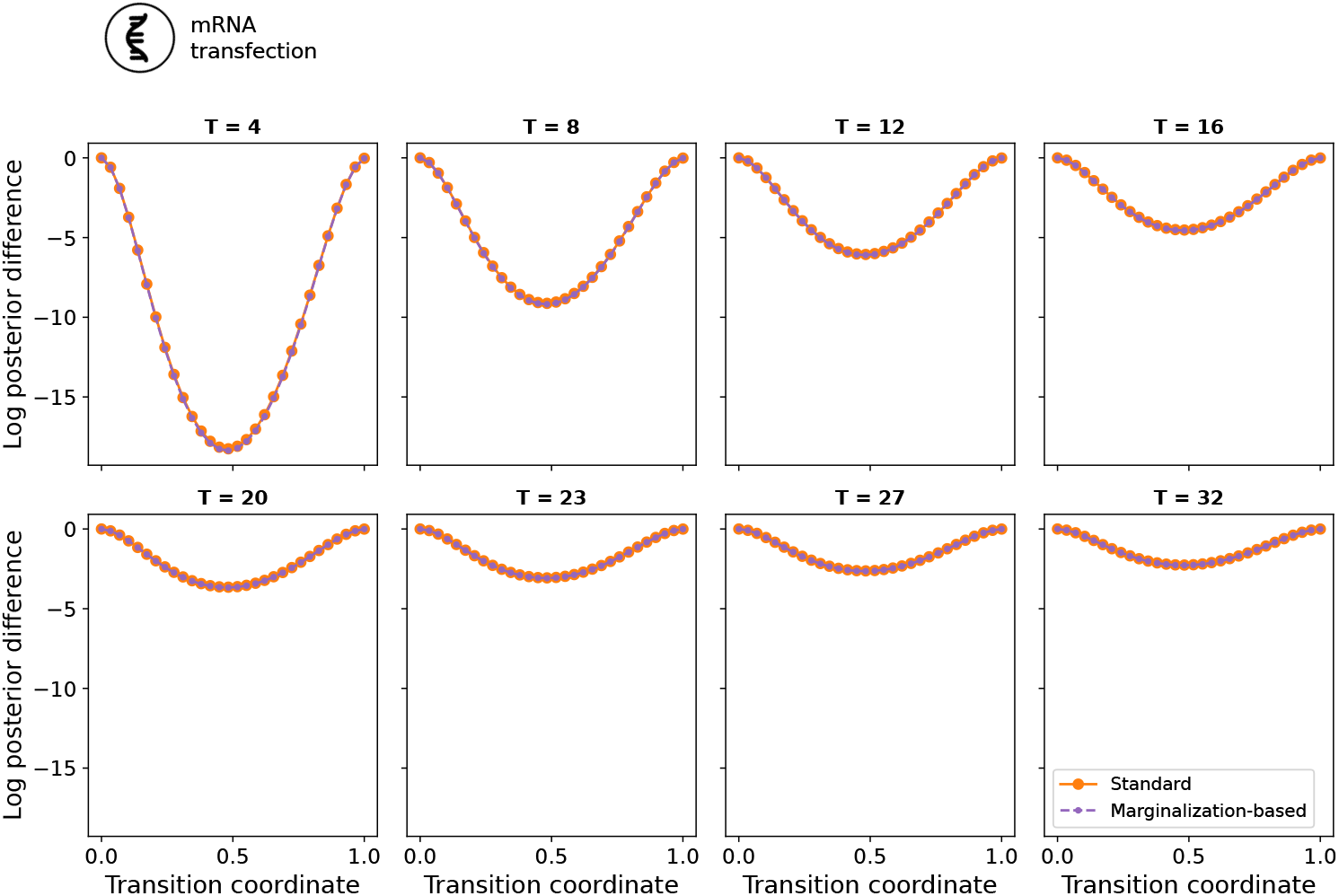
Minimum energy path of the tempered posteriors for a range of temperatures considered in Figure 5A for model M3.

**Figure S7:**
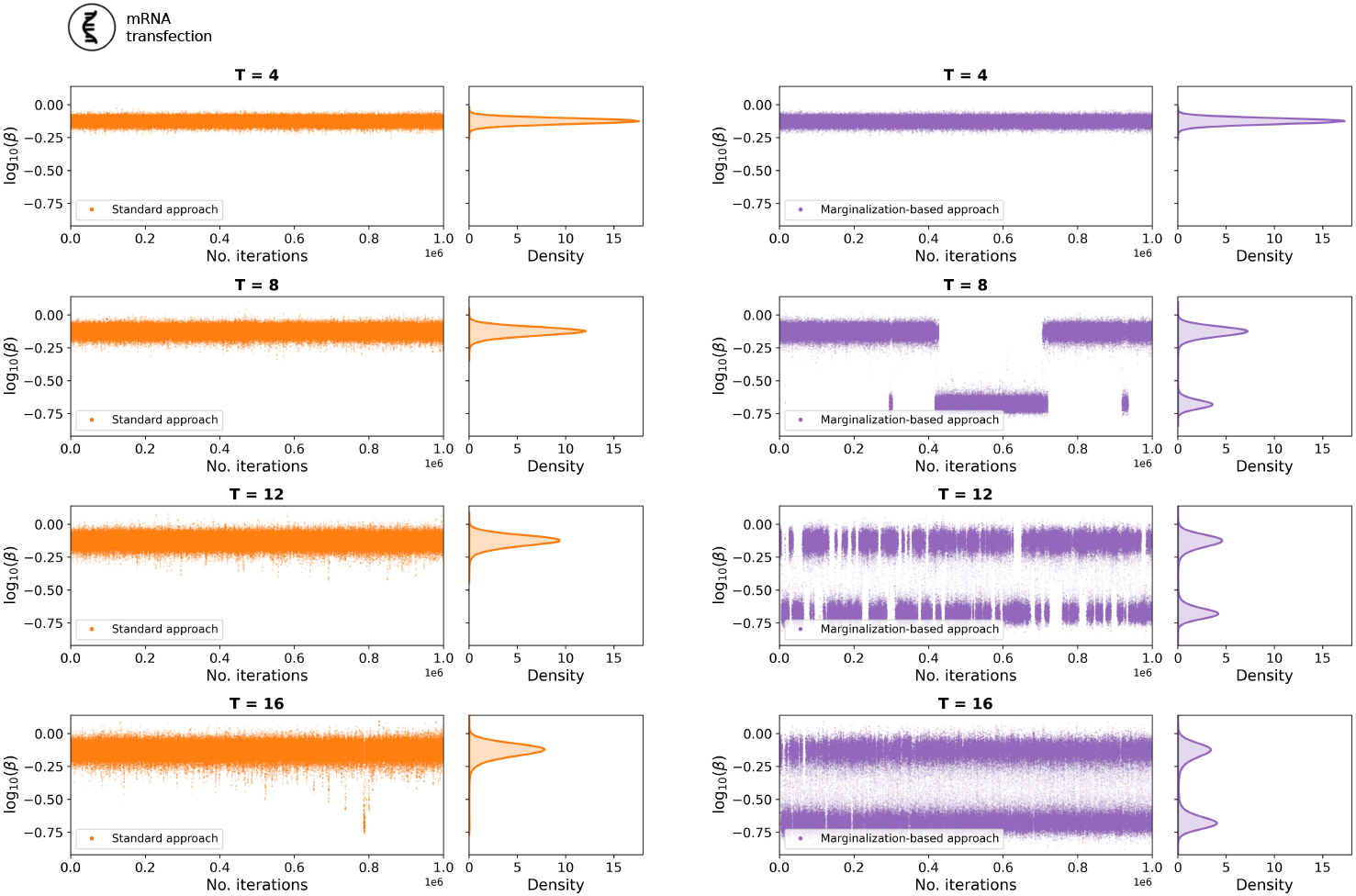
Representative parameter traces for the model parameter *β* for a range of temperatures considered in Figure 5A for model M3.

**Figure S8:**
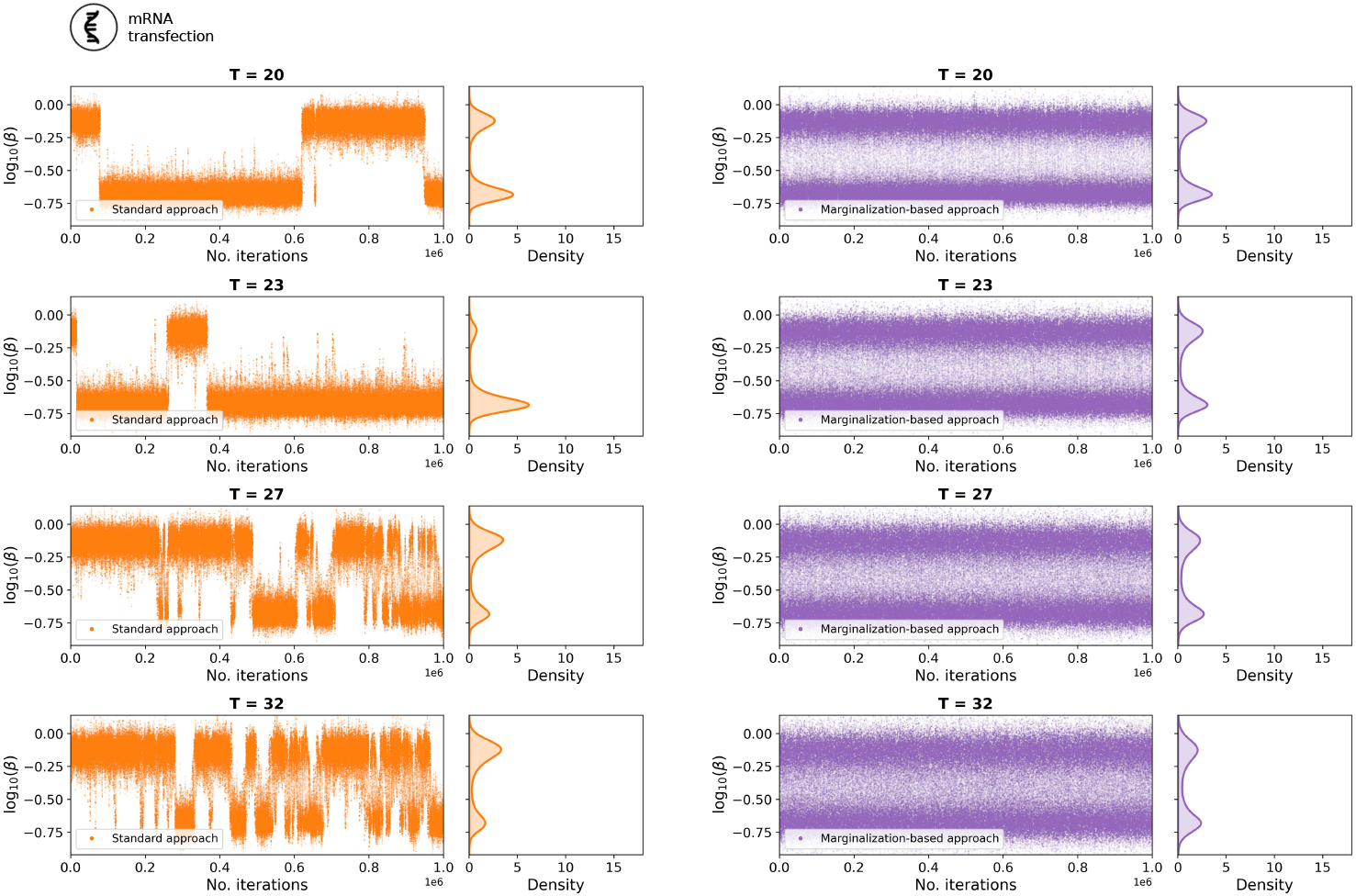
Representative parameter traces for the model parameter *β* for a range of temperatures considered in Figure 5A for model M3.

**Figure S9:**
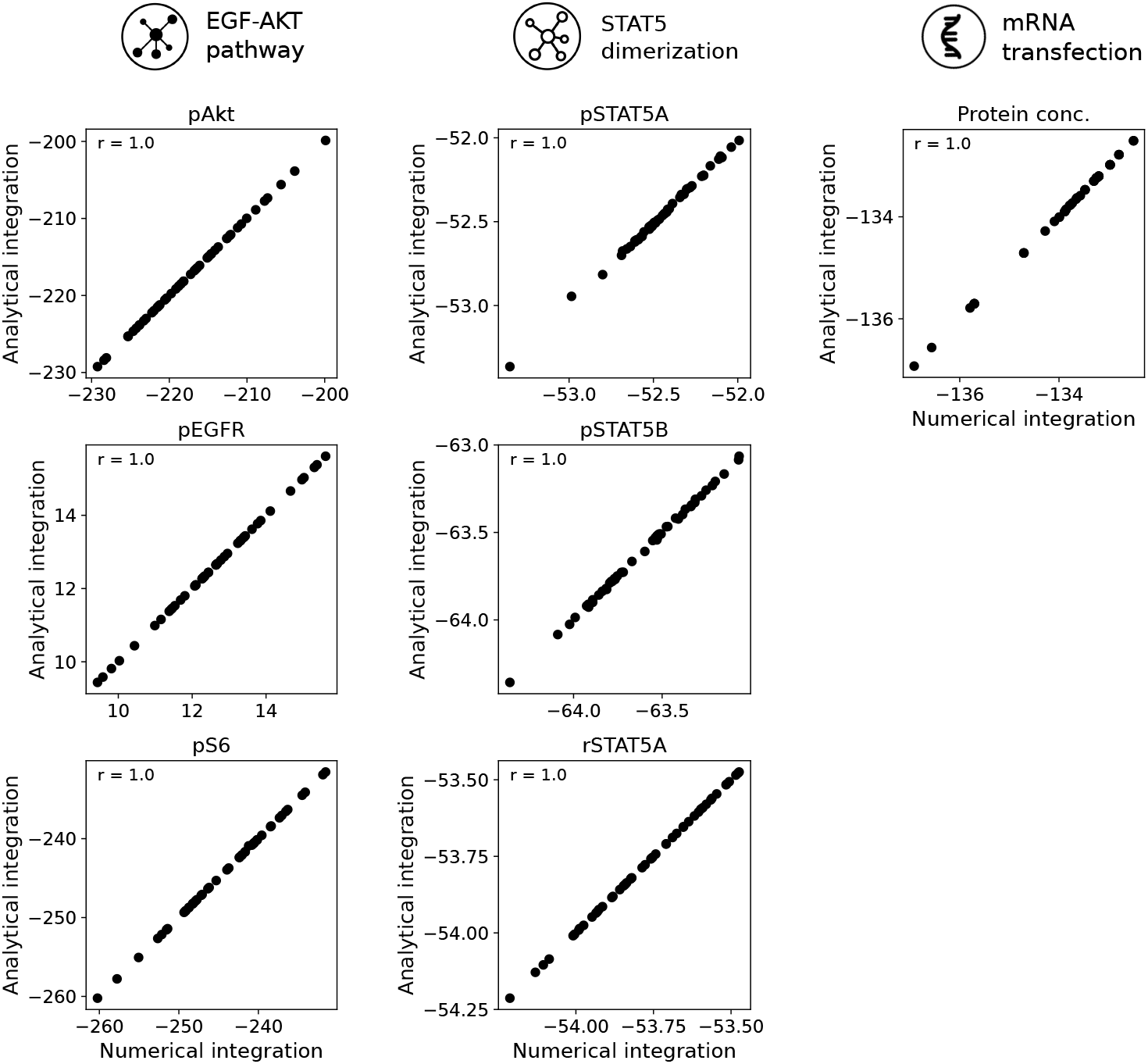
Correctness of the analytical integration for model M1, M2 and M3. Scatter plot for the agreement of analytical and numerical integration for 50 different parameter vectors. The integration results are shown for each model observable.

**Figure S10:**
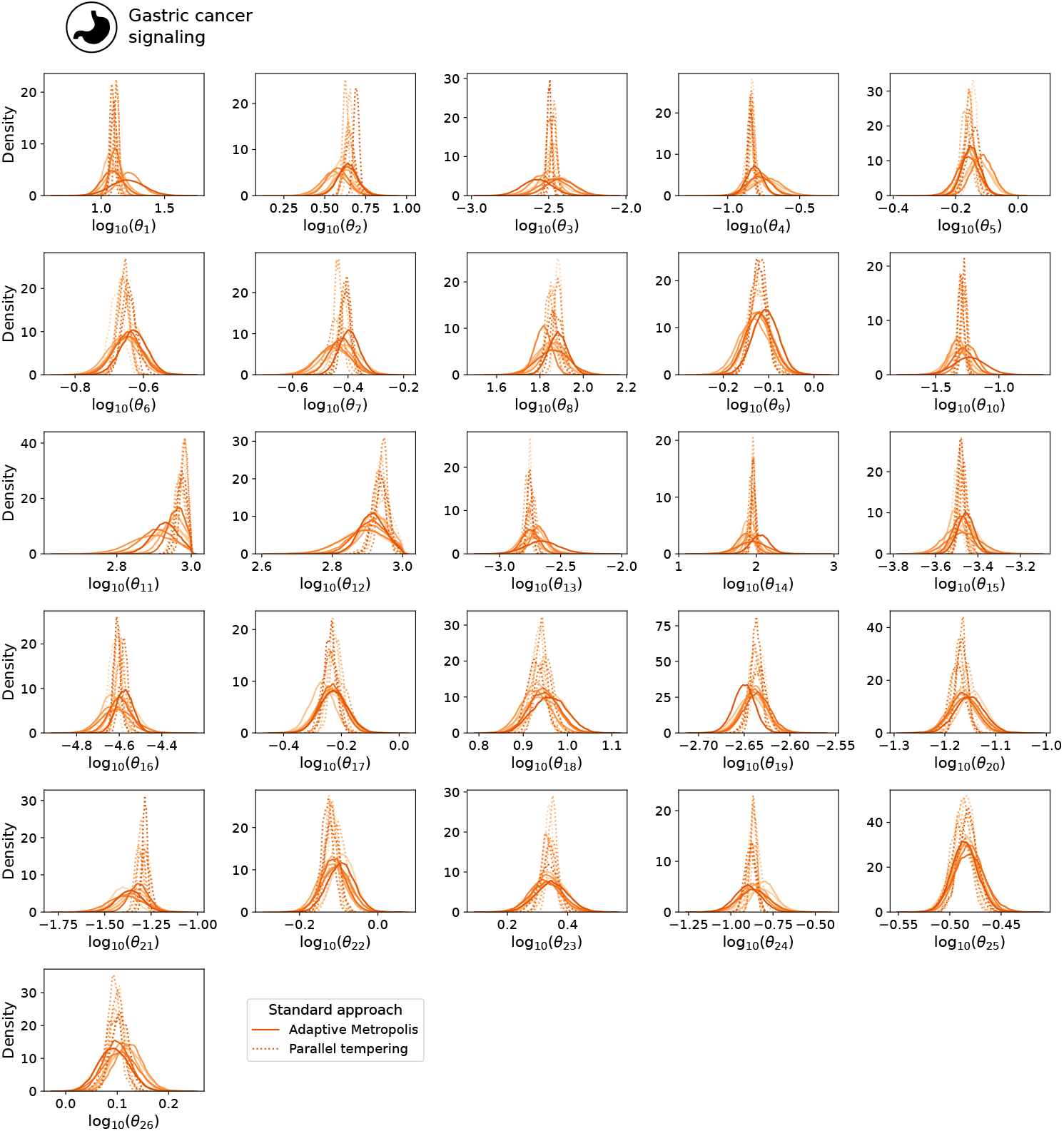
Parameter marginal posterior distributions using the standard approach for model M4. Results from two sampling algorithms (adaptive Metropolis and parallel tempering) and only the subset of model parameters are shown. The marginals were computed using a kernel density estimate.

**Figure S11:**
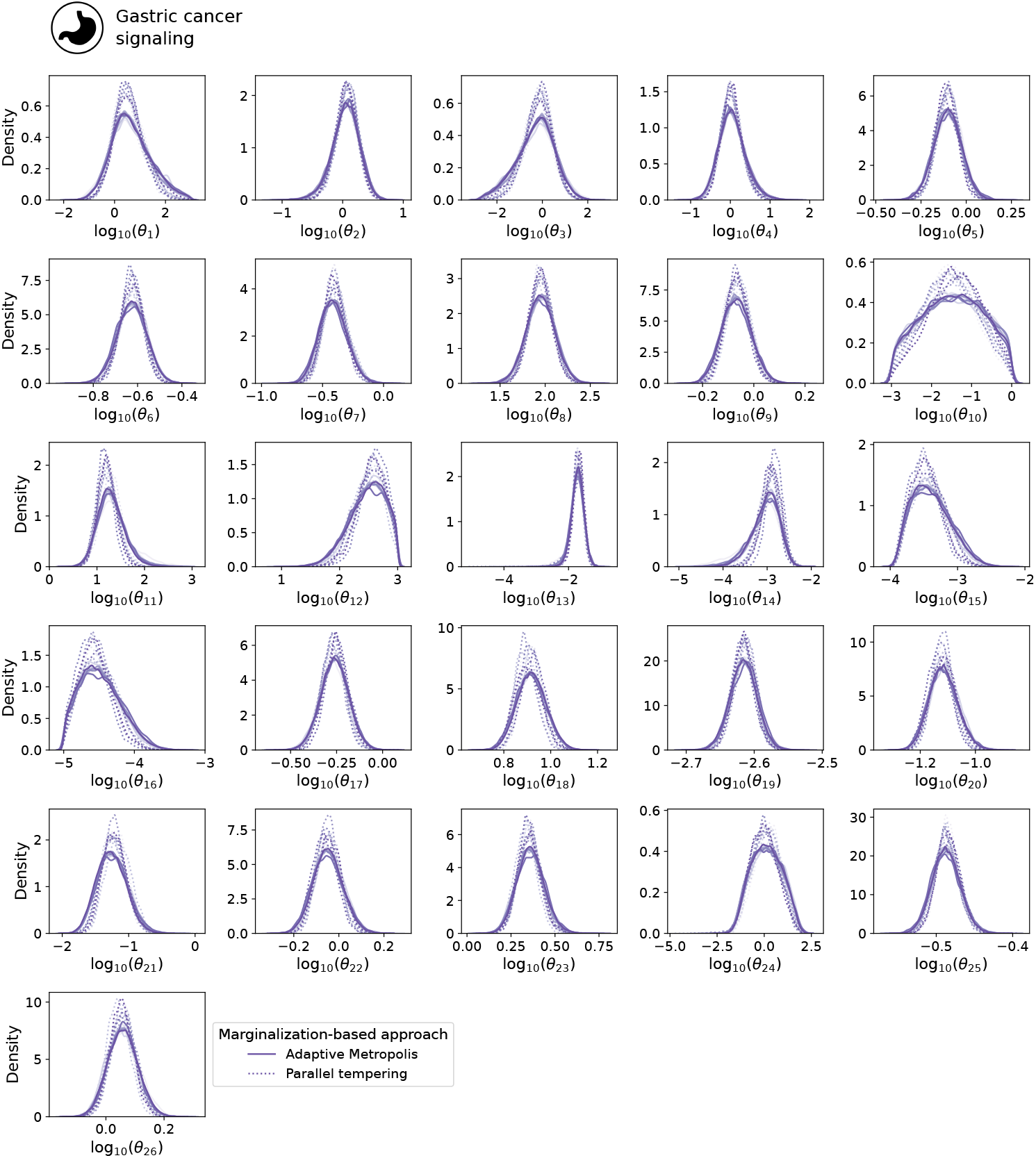
Parameter marginal posterior distributions using the marginalization-based approach for model M4. Results from two sampling algorithms (adaptive Metropolis and parallel tempering) and only the subset of model parameters are shown. The marginals were computed using a kernel density estimate.

**Figure S12:**
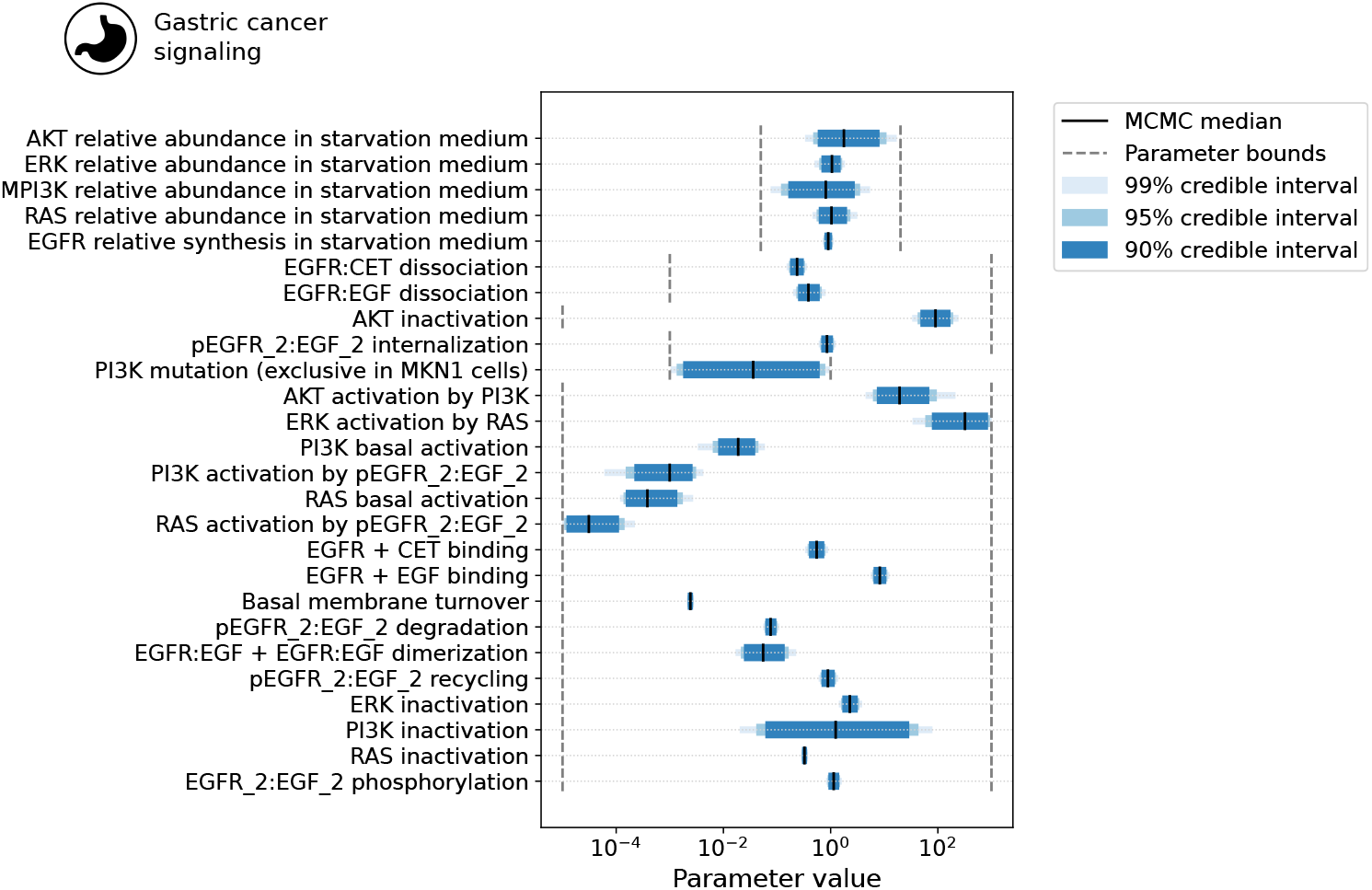
Credible intervals for the model parameters of model M4. The credible intervals were extracted from the MCMC samples obtained with the marginalization-based approach. The credible levels 90%, 95% and 99% are shown. Parameter bounds used for sampling are indicated in black dashed lines. Only the subset of model parameters are shown.

**Table S1:**
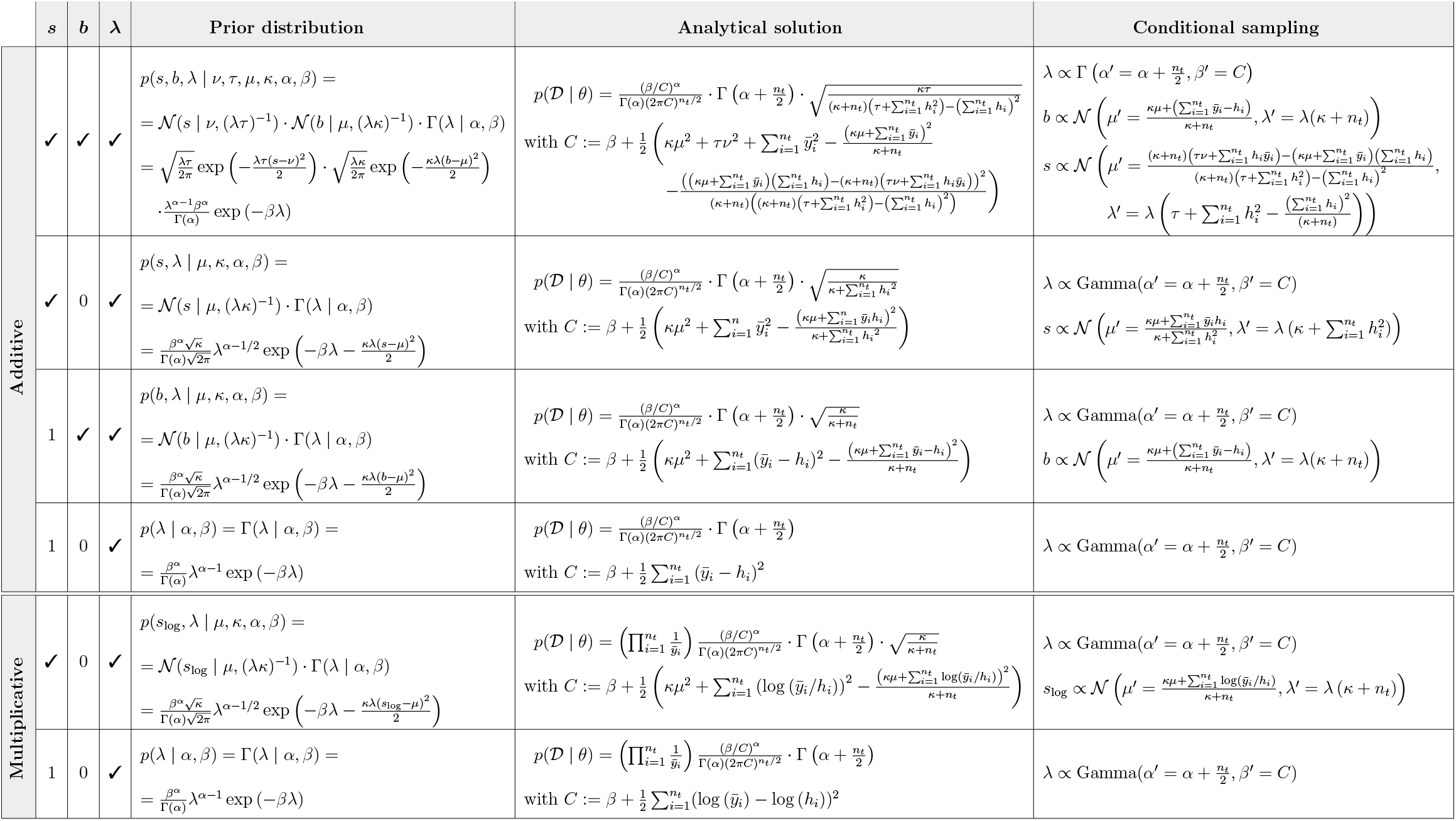
Overview of the marginalization-based approach applied to different observable combinations under unknown additive and multiplicative Gaussian measurement noise. Observation parameters considered are scaling factors (s) and offsets (b). The noise is denoted as precision *λ*:= 1/*σ*^2^. For multiplicative noise, the logarithm of the scaling factor (*s*_log_) is used. Unknown/estimated observation parameters are denoted by ⌔, otherwise the fixed numerical value is shown. Further details for each case are in the *Supplementary Material*.

**Table S2:**
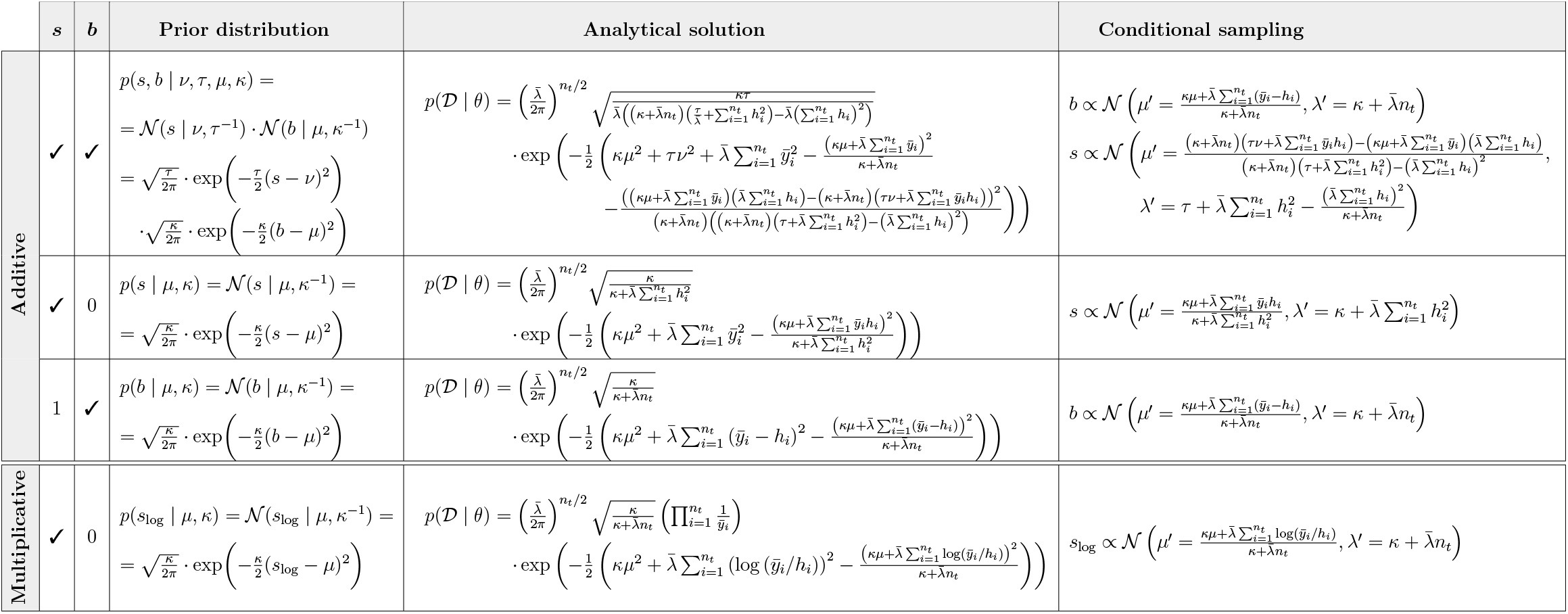
Overview of the marginalization-based approach applied to different observable combinations under experimentally measured additive and multiplicative Gaussian measurement noise. Observation parameters considered are scaling factors (*s*) and offsets (*b*). The experimentally measured noise is denoted as precision 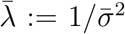. For multiplicative noise, the logarithm of the scaling factor (*s*_log_) is used. Unknown/estimated observation parameters are denoted by ✔, otherwise the fixed numerical value is shown. Further details for each case are in the *Supplementary Material*.

