## Supplementary Material for "Posterior marginalization accelerates Bayesian inference for dynamical systems"

**5 Benchmark models** **36**

### 1 Mathematical model

We consider systems of ordinary differential equations (ODEs),

$$\frac{\mathrm{d}x(t,\theta)}{\mathrm{d}t} = f(x(t,\theta),\theta), \quad x(t_0,\theta) = x_0(\theta)$$

with state vector $x$, parameter vector $\theta$, and time $t$. The vector field is denoted by $f$ and the initial condition by $x_0$. The parameters $\theta$ are assumed to be unknown and need to be estimated from the experimental data.

We assume that experimental data $\mathcal{D}$ are obtained by noise-corrupted measurements of the observables

$$y(t,\theta,s,b) = \mathrm{diag}(s) \cdot h(x(t,\theta),\theta) + b,$$

with scaling vector $s$ and offset vector $b$. The measurements are time discrete and collected at time points $t_k$, $k = 1,\dots,N$. To shorten the notation, we define $h_k := h(x(t_k,\theta),\theta)$.

To simplify the notation, we consider in the following sections the case of a scalar observable. Yet, our results generalise to multiple observables which do not share observation and noise parameters. Furthermore, we only consider a single experimental condition without pre-equilibration, while the results generalize to multiple experimental conditions with and without pre-equilibration.

In the following, we present results for the analytical marginalization of the posterior distribution for different combinations of observation and noise models.

### 2 Marginalization-based approach for additive Gaussian noise

In this section, we consider *additive Gaussian* measurement noise. The data set $\mathcal{D}$ is a collection of measurements

$$\bar{y}_k = (s \cdot h_k + b) + \epsilon_k, \quad \epsilon_k \sim \mathcal{N}(0,\sigma^2), k = 1,\dots,N.$$

For simplicity of notation, we write the problem in terms of precision $\lambda$, which is the inverse of the variance $\sigma^2$.

Bayes' theorem for unknown model parameter vector $\theta$, unknown scaling factors $s$, unknown offsets $b$, and unknown noise parameters $\lambda$ is given by

$$p(\theta,s,b,\lambda \mid \mathcal{D}) = \frac{p(\mathcal{D} \mid \theta,s,b,\lambda)p(\theta,s,b,\lambda)}{p(\mathcal{D})},$$

with prior $p(\theta,s,b,\lambda)$, marginal $p(\mathcal{D})$ and likelihood

$$p(\mathcal{D} \mid \theta,s,b,\lambda) = \frac{1}{(2\pi)^{N/2}}\lambda^{N/2} \exp\left(-\frac{\lambda}{2}\sum_{k=1}^{N}(\bar{y}_k - (s \cdot h_k + b))^2\right)$$

In the following, we show different combinations of known and unknown parameters.

#### 2.1 Unknown noise and scaling parameters

We consider the observation and noise model

$$\bar{y}_k = s \cdot h_k + \epsilon_k, \quad \epsilon_k \sim \mathcal{N}(0, 1/\lambda),$$

with unknown scaling parameter $s$ and unknown noise parameter $\lambda$. For these unknown parameters we assume a Normal-Gamma prior,

$$\begin{aligned} p(s,\lambda) &= \mathcal{N}(s \mid \mu, 1/(\kappa\lambda)) \cdot \Gamma(\lambda \mid \alpha, \beta) \\ &= \frac{\beta^\alpha \sqrt{\kappa}}{\Gamma(\alpha)\sqrt{2\pi}} \lambda^{\alpha-1/2} \exp\left(-\beta\lambda - \frac{\kappa\lambda(s-\mu)^2}{2}\right) \end{aligned}$$

with $\mu$, $\alpha$, $\beta$, $\kappa$ as hyper-parameters. $\Gamma(\cdot)$ denotes the Gamma function. The marginalized likelihood is denoted by

$$\begin{aligned} p(\mathcal{D} \mid \theta) &= \int_0^\infty \int_{-\infty}^\infty p(\mathcal{D} \mid \theta, s, \lambda) p(s,\lambda) \, \mathrm{d}s \, \mathrm{d}\lambda \\ &= \int_0^\infty \int_{-\infty}^\infty \left(\frac{\lambda}{2\pi}\right)^{N/2} \exp\left(-\frac{\lambda}{2}\sum_{k=1}^N (\bar{y}_k - s\cdot h_k)^2\right) \cdot \frac{\beta^\alpha\sqrt{\kappa}}{\Gamma(\alpha)\sqrt{2\pi}} \lambda^{\alpha-1/2} \exp\left(-\beta\lambda - \frac{\kappa\lambda(s-\mu)^2}{2}\right) \, \mathrm{d}s \, \mathrm{d}\lambda. \end{aligned}$$

We reformulate the marginalized likelihood by removing constants from the integral and by reordering the terms under the integral. This yields

$$\begin{aligned} p(\mathcal{D} \mid \theta) &= \frac{\beta^\alpha\sqrt{\kappa}}{\Gamma(\alpha)\sqrt{2\pi}} \frac{1}{(2\pi)^{N/2}} \int_0^\infty \lambda^{\alpha-1/2}\lambda^{N/2} \int_{-\infty}^\infty \exp\left(-\frac{\lambda}{2}\sum_{k=1}^N (\bar{y}_k - s\cdot h_k)^2\right) \cdot \exp\left(-\beta\lambda - \frac{\kappa\lambda(s-\mu)^2}{2}\right) \, \mathrm{d}s \, \mathrm{d}\lambda \\ &= \frac{\beta^\alpha\sqrt{\kappa}}{\Gamma(\alpha)(2\pi)^{(N+1)/2}} \int_0^\infty \lambda^{\alpha+(N-1)/2} \exp(-\beta\lambda) \underbrace{\int_{-\infty}^\infty \exp\left(-\frac{\lambda}{2}\left(\sum_{k=1}^N (\bar{y}_k - s\cdot h_k)^2 + \kappa(s-\mu)^2\right)\right) \, \mathrm{d}s}_{(*)} \, \mathrm{d}\lambda. \end{aligned}$$

The integral $(*)$ with respect to $s$ can be written as

$$\begin{aligned} (*) &= \int_{-\infty}^\infty \exp\left(-\frac{\lambda}{2}\left(s^2\sum_{k=1}^N h_k^2 - 2s\sum_{k=1}^N \bar{y}_k h_k + \sum_{k=1}^N \bar{y}_k^2 + \kappa s^2 - 2\kappa\mu s + \kappa\mu^2\right)\right) \, \mathrm{d}s \\ &= \int_{-\infty}^\infty \exp\left(-\frac{\lambda}{2}\left(\left(\kappa + \sum_{k=1}^N h_k^2\right) s^2 - \left(2\sum_{k=1}^N \bar{y}_k h_k + 2\kappa\mu\right) s + \left(\sum_{k=1}^N \bar{y}_k^2 + \kappa\mu^2\right)\right)\right) \, \mathrm{d}s \\ &= \int_{-\infty}^\infty \exp\left(-\frac{\lambda}{2}\left(\kappa + \sum_{k=1}^N h_k^2\right) s^2 + \lambda\left(\sum_{k=1}^N \bar{y}_k h_k + \kappa\mu\right) s - \frac{\lambda}{2}\left(\sum_{k=1}^N \bar{y}_k^2 + \kappa\mu^2\right)\right) \, \mathrm{d}s. \end{aligned} \tag{1}$$

This is a Gaussian integral of the type

$$\int_{-\infty}^\infty \exp\left(-Ax^2 + Bx + C\right) \, \mathrm{d}x = \sqrt{\frac{\pi}{A}} \exp\left(\frac{B^2}{4A} + C\right) \tag{2}$$

with constants

$$A := \frac{\lambda}{2}\left(\kappa + \sum_{k=1}^{N} h_k^2\right), \qquad B := \lambda\left(\sum_{k=1}^{N} \bar{y}_k h_k + \kappa\mu\right), \qquad \text{and} \qquad C := -\frac{\lambda}{2}\left(\sum_{k=1}^{N} \bar{y}_k^2 + \kappa\mu^2\right).$$

As $\lambda, h_k^2, \kappa > 0$, $A > 0$ can be assured, we obtain

$$\begin{aligned}(*) &= \sqrt{\frac{\pi}{\frac{\lambda}{2}\left(\kappa + \sum_{k=1}^{N} h_k^2\right)}} \exp\left(\frac{\left(\lambda\left(\sum_{k=1}^{N} \bar{y}_k h_k + \kappa\mu\right)\right)^2}{4\frac{\lambda}{2}\left(\kappa + \sum_{k=1}^{N} h_k^2\right)} - \frac{\lambda}{2}\left(\sum_{k=1}^{N} \bar{y}_k^2 + \kappa\mu^2\right)\right) \\ &= \sqrt{\frac{2\pi}{\lambda\left(\kappa + \sum_{k=1}^{N} h_k^2\right)}} \exp\left(-\lambda\tilde{\omega}\right)\end{aligned}$$

with

$$\tilde{\omega} := -\frac{1}{2}\left(\frac{\left(\sum_{k=1}^{N} \bar{y}_k h_k + \kappa\mu\right)^2}{\left(\kappa + \sum_{k=1}^{N} h_k^2\right)} - \left(\sum_{k=1}^{N} \bar{y}_k^2 + \kappa\mu^2\right)\right).$$

The substitution of this result for the integral over $s$ in the integral formulation yields

$$\begin{aligned}p(\mathcal{D} \mid \theta) &= \frac{\beta^\alpha \sqrt{\kappa}}{\Gamma(\alpha)(2\pi)^{(N+1)/2}} \int_0^\infty \lambda^{\alpha+(N-1)/2} \exp\left(-\beta\lambda\right) \sqrt{\frac{2\pi}{\lambda\left(\kappa + \sum_{k=1}^{N} h_k^2\right)}} \exp\left(-\lambda\tilde{\omega}\right)\ \mathrm{d}\lambda \\ &= \frac{\beta^\alpha \sqrt{\kappa}}{\Gamma(\alpha)(2\pi)^{(N+1)/2}} \sqrt{\frac{2\pi}{\left(\kappa + \sum_{k=1}^{N} h_k^2\right)}} \int_0^\infty \lambda^{\alpha+N/2-1} \exp\left(-\lambda\omega\right)\ \mathrm{d}\lambda \qquad (3)\end{aligned}$$

with

$$\omega := \beta + \tilde{\omega} = \beta - \frac{1}{2}\left(\frac{\left(\sum_{k=1}^{N} \bar{y}_k h_k + \kappa\mu\right)^2}{\left(\kappa + \sum_{k=1}^{N} h_k^2\right)} - \left(\sum_{k=1}^{N} \bar{y}_k^2 + \kappa\mu^2\right)\right).$$

Using the transformation $z = \lambda\omega$, we obtain for the marginalized likelihood

$$\begin{aligned}p(\mathcal{D} \mid \theta) &= \frac{\beta^\alpha \sqrt{\kappa}}{\Gamma(\alpha)(2\pi)^{(N+1)/2}} \sqrt{\frac{2\pi}{\left(\kappa + \sum_{k=1}^{N} h_k^2\right)}} \int_0^\infty \left(\frac{z}{\omega}\right)^{\alpha+N/2-1} \exp\left(-z\right) \frac{1}{\omega}\,\mathrm{d}z \\ &= \left(\frac{1}{\omega}\right)^\delta \cdot \frac{\beta^\alpha \sqrt{\kappa}}{\Gamma(\alpha)(2\pi)^{(N+1)/2}} \sqrt{\frac{2\pi}{\left(\kappa + \sum_{k=1}^{N} h_k^2\right)}} \int_0^\infty z^{\delta-1} \exp\left(-z\right)\ \mathrm{d}z \qquad (4)\end{aligned}$$

with $\delta := \alpha + N/2$. The integral in (4) can be solved analytically since it follows the structure

$$\int_0^\infty z^{\delta-1} \exp\left(-z\right)\ \mathrm{d}z = \Gamma(\delta). \qquad (5)$$

This yields a closed-form solution for the marginalized likelihood,

$$
\begin{aligned}
p(\mathcal{D} \mid \theta) &= \left(\frac{1}{\omega}\right)^{\delta} \cdot \frac{\beta^{\alpha}\sqrt{\kappa}}{\Gamma(\alpha)(2\pi)^{(N+1)/2}} \sqrt{\frac{2\pi}{\left(\kappa + \sum_{k=1}^{N} h_k^2\right)}} \cdot \Gamma(\delta) \\
&= \left(\frac{1}{\omega}\right)^{\alpha+N/2} \cdot \frac{\beta^{\alpha}\sqrt{\kappa}}{\Gamma(\alpha)(2\pi)^{(N+1)/2}} \sqrt{\frac{2\pi}{\left(\kappa + \sum_{k=1}^{N} h_k^2\right)}} \cdot \Gamma\left(\alpha + \frac{N}{2}\right) \\
&= \frac{(\beta/\omega)^{\alpha}}{\Gamma(\alpha)(2\pi\omega)^{N/2}} \cdot \sqrt{\frac{\kappa}{\kappa + \sum_{k=1}^{N} h_k^2}} \cdot \Gamma\left(\alpha + \frac{N}{2}\right).
\end{aligned}
$$

##### 2.1.1 Retrieving the marginalized parameters

The conditional probability of the marginalized noise and scaling parameters can be derived after obtaining $p(\theta \mid \mathcal{D})$ given values of $\theta$.

#### Sampling of the noise parameter

The integrand in (3) shows that the precision $\lambda$ has (up to a constant) the distribution of a Gamma distribution, mainly

$$
\lambda \propto \text{Gamma}(\alpha' = \alpha + N/2, \beta' = \omega).
$$

Similarly

$$
\sigma^2 \propto \text{Inv-Gamma}(\alpha' = \alpha + N/2, \beta' = \omega).
$$

The distribution is well-defined if $\alpha', \beta' \in \mathbb{R}_+$, which holds for both parameters. While for $\alpha'$ this is clear by definition, for $\beta'$ we consider the following

$$
\omega \in \mathbb{R}_+ \Leftrightarrow \beta + \frac{1}{2}\left(\kappa\mu^2 + \sum_{k=1}^{N} \bar{y}_k^2\right) > \frac{1}{2}\frac{\left(\kappa\mu + \sum_{k=1}^{N} \bar{y}_k h_k\right)^2}{\left(\kappa + \sum_{k=1}^{N} h_k^2\right)}.
$$

As $\beta > 0$ it is sufficient to show that

$$
\left(\kappa\mu^2 + \sum_{k=1}^{N} \bar{y}_k^2\right) \geq \frac{\left(\kappa\mu + \sum_{k=1}^{N} \bar{y}_k h_k\right)^2}{\left(\kappa + \sum_{k=1}^{N} h_k^2\right)}. \tag{6}
$$

Then, following Cauchy–Schwarz inequality

$$
\left(\sum_{k=1}^{N} a_k \cdot \rho_k\right)^2 \leq \left(\sum_{k=1}^{N} a_k^2\right) \cdot \left(\sum_{k=1}^{N} \rho_k^2\right) \tag{7}
$$

with $a_k = \bar{y}_k,\ \rho_k = h_k$ for $1 \le k \le N$ and $a_{N+1} = \sqrt{\kappa}\mu,\ b_{N+1} = \sqrt{\kappa}$, we receive

$$\left(\kappa\mu + \sum_{k=1}^{N} \bar{y}_k h_k\right)^2 \le \left(\kappa\mu^2 + \sum_{k=1}^{N} \bar{y}_k^2\right) \cdot \left(\kappa + \sum_{k=1}^{N} h_k^2\right).$$

By dividing through $\left(\kappa + \sum_{k=1}^{N} h_k^2\right)$ we get the desired result (6).

##### Sampling of the scaling parameter

For $s$ the integrand in (1) can be considered and we get

$$\begin{aligned}
s &\propto \exp\left(-\frac{\lambda}{2}\left(\kappa + \sum_{k=1}^{N} h_k^2\right) s^2 + \lambda\left(\sum_{k=1}^{N} \bar{y}_k h_k + \kappa\mu\right) s\right) \\
&\propto \exp\left(-\frac{1}{2}\frac{\left(s - \frac{\left(\sum_{k=1}^{N} \bar{y}_k h_k + \kappa\mu\right)}{\left(\kappa + \sum_{k=1}^{N} h_k^2\right)}\right)^2}{\sigma^2 / \left(\kappa + \sum_{k=1}^{N} h_k^2\right)}\right) \\
&\propto \mathcal{N}\left(\mu' = \frac{\left(\sum_{k=1}^{N} \bar{y}_k h_k + \kappa\mu\right)}{\left(\kappa + \sum_{k=1}^{N} h_k^2\right)}, (\sigma')^2 = \frac{\sigma^2}{\left(\kappa + \sum_{k=1}^{N} h_k^2\right)}\right)
\end{aligned}$$

The distribution is well-defined if $\mu' \in \mathbb{R}$ and $(\sigma')^2 \in \mathbb{R}_+$, which holds for both and is clear by definition.

#### 2.2 Unknown noise and offset parameters

We consider the observation and noise model

$$\bar{y}_k = h_k + b + \epsilon_k, \quad \epsilon_k \sim \mathcal{N}(0, 1/\lambda),$$

with unknown offset parameter $b$ and unknown noise parameter $\lambda$. For these unknown parameters we assume a Normal-Gamma prior,

$$\begin{aligned}
p(b, \lambda) &= \mathcal{N}(b \mid \mu, 1/(\kappa\lambda)) \cdot \Gamma(\lambda \mid \alpha, \beta) \\
&= \frac{\beta^\alpha \sqrt{\kappa}}{\Gamma(\alpha)\sqrt{2\pi}} \lambda^{\alpha - 1/2} \exp\left(-\beta\lambda - \frac{\kappa\lambda(b-\mu)^2}{2}\right)
\end{aligned}$$

with $\mu$, $\alpha$, $\beta$, $\kappa$ as hyper-parameters. $\Gamma(\cdot)$ denotes the Gamma function. The marginalized likelihood is denoted by

$$\begin{aligned}
p(\mathcal{D} \mid \theta) &= \int_0^\infty \int_{-\infty}^\infty p(\mathcal{D} \mid \theta, b, \lambda) p(b, \lambda) \,\mathrm{d}b \,\mathrm{d}\lambda \\
&= \int_0^\infty \int_{-\infty}^\infty \left(\frac{\lambda}{2\pi}\right)^{N/2} \exp\left(-\frac{\lambda}{2}\sum_{k=1}^{N} (\bar{y}_k - (h_k + b))^2\right) \cdot \frac{\beta^\alpha \sqrt{\kappa}}{\Gamma(\alpha)\sqrt{2\pi}} \lambda^{\alpha - 1/2} \exp\left(-\beta\lambda - \frac{\kappa\lambda(b-\mu)^2}{2}\right) \mathrm{d}b \,\mathrm{d}\lambda
\end{aligned}$$

We reformulate the marginalized likelihood by removing constants from the integral and by reordering the terms under the integral. This yields

$$p(\mathcal{D} \mid \theta) = \frac{\beta^\alpha \sqrt{\kappa}}{\Gamma(\alpha)(2\pi)^{\frac{N+1}{2}}} \int_0^\infty \lambda^{\alpha+\frac{N-1}{2}} \underbrace{\int_{-\infty}^{\infty} \exp\left(-\frac{\lambda}{2}\left(\left(\sum_{k=1}^{N}((\bar{y}_k - h_k) - b)^2\right) + \kappa(b-\mu)^2 + 2\beta\right)\right) \mathrm{d}b}_{(*)} \,\mathrm{d}\lambda.$$

The integral $(*)$ with respect to $b$ can be written as

$$\begin{aligned}(*) &= \int_{-\infty}^{\infty} \exp\left(-\frac{\lambda}{2}\left(\left(\sum_{k=1}^{N}((\bar{y}_k - h_k) - b)^2\right) + \kappa(b-\mu)^2 + 2\beta\right)\right) \mathrm{d}b \\ &= \int_{-\infty}^{\infty} \exp\left(-\frac{\lambda}{2}\left((N+\kappa)b^2 - 2\left(\left(\sum_{k=1}^{N}\bar{y}_k - h_k\right) + \kappa\mu\right) b + \left(\sum_{k=1}^{N}(\bar{y}_k - h_k)^2\right) + \kappa\mu^2 + 2\beta\right)\right) \mathrm{d}b \end{aligned} \tag{8}$$

This is a Gaussian integral of the type (2) with constants

$$A := \frac{\lambda(N+\kappa)}{2}, \quad B := \lambda\left(\left(\sum_{k=1}^{N}\bar{y}_k - h_k\right) + \kappa\mu\right), \quad \text{and} \quad C := -\frac{\lambda}{2}\left(\left(\sum_{k=1}^{N}(\bar{y}_k - h_k)^2\right) + \kappa\mu^2 + 2\beta\right).$$

As $\lambda, N, \kappa > 0$, $A > 0$ can be assured, we obtain

$$(*) = \sqrt{\frac{2\pi}{\lambda(N+\kappa)}} \cdot \exp\left(\frac{\lambda}{2} \cdot \left(\frac{\left(\left(\sum_{k=1}^{N}\bar{y}_k - h_k\right) + \kappa\mu\right)^2}{(N+\kappa)} - \left(\left(\sum_{k=1}^{N}(\bar{y}_k - h_k)^2\right) + \kappa\mu^2 + 2\beta\right)\right)\right)$$

The substitution of this result for the integral over $b$ in the integral formulation yields

$$\begin{aligned} p(\mathcal{D} \mid \theta) &= \frac{\beta^\alpha \sqrt{\kappa}}{\Gamma(\alpha)(2\pi)^{\frac{N+1}{2}}} \cdot \sqrt{\frac{2\pi}{(N+\kappa)}} \\ &\quad \cdot \int_0^\infty \lambda^{\alpha+\frac{N-1}{2}} \cdot \sqrt{\frac{1}{\lambda}} \cdot \exp\left(\frac{\lambda}{2} \cdot \left(\frac{\left(\left(\sum_{k=1}^{N}\bar{y}_k - h_k\right) + \kappa\mu\right)^2}{(N+\kappa)} - \left(\left(\sum_{k=1}^{N}(\bar{y}_k - h_k)^2\right) + \kappa\mu^2 + 2\beta\right)\right)\right) \mathrm{d}\lambda \\ &= \frac{\beta^\alpha}{\Gamma(\alpha)(2\pi)^{\frac{N}{2}}} \cdot \sqrt{\frac{\kappa}{N+\kappa}} \int_0^\infty \lambda^{\alpha+\frac{N}{2}-1} \cdot \exp(-\lambda\omega)\,\mathrm{d}\lambda \end{aligned} \tag{9}$$

with

$$\omega := \frac{1}{2} \cdot \left(\kappa\mu^2 + 2\beta + \sum_{k=1}^{N}(\bar{y}_k - h_k)^2 - \frac{\left(\kappa\mu + \sum_{k=1}^{N}\bar{y}_k - h_k\right)^2}{N+\kappa}\right)$$

Using the transformation $z = \lambda\omega$ and using the definition of the Gamma function (5), we obtain a closed-form solution for the marginalized likelihood

$$\begin{aligned} p(\mathcal{D} \mid \theta) &= \frac{\beta^\alpha}{\Gamma(\alpha)(2\pi)^{\frac{N}{2}}} \cdot \sqrt{\frac{\kappa}{N+\kappa}} \cdot \omega^{-\left(\alpha+\frac{N}{2}\right)} \int_0^\infty z^{(\alpha+\frac{N}{2})-1} \cdot \exp(-z)\,\mathrm{d}z \\ &= \frac{(\beta/\omega)^\alpha}{\Gamma(\alpha)(2\pi\omega)^{\frac{N}{2}}} \cdot \sqrt{\frac{\kappa}{N+\kappa}} \cdot \Gamma\left(\alpha + \frac{N}{2}\right) \end{aligned}$$

##### 2.2.1 Retrieving the marginalized parameters

The conditional probability of the marginalized noise and offset parameters can be derived after obtaining $p(\theta \mid \mathcal{D})$ given values of $\theta$.

###### Sampling of the noise parameter

The integrand in (9) shows that the precision $\lambda$ has (up to a constant) the distribution of a Gamma distribution, mainly

$$\lambda \propto \mathrm{Gamma}(\alpha' = \alpha + N/2, \beta' = \omega).$$

This distribution is well defined if $\alpha', \beta' \in \mathbb{R}_+$ which holds for both parameters. For $\alpha'$ this is clear by definition, for $\beta'$ consider the following calculation:

$$\omega \in \mathbb{R}_+ \Leftrightarrow \left(\sum_{k=1}^{N} (\bar{y}_k - h_k)^2\right) + \kappa\mu^2 + 2\beta > \frac{1}{N+\kappa}\left(\left(\sum_{k=1}^{N} \bar{y}_k - h_k\right) + \kappa\mu\right)^2$$

As $\beta > 0$ it is sufficient to show that

$$\left(\sum_{k=1}^{N} (\bar{y}_k - h_k)^2\right) + \kappa\mu^2 \geq \frac{1}{N+\kappa}\left(\left(\sum_{k=1}^{N} \bar{y}_k - h_k\right) + \kappa\mu\right)^2$$

Following Cauchy–Schwarz inequality (7) with $a_k = \bar{y}_k - h_k$, $\rho_k = 1$ for $1 \leq k \leq N$ and $a_{N+1} = \sqrt{\kappa}\mu$, $b_{N+1} = \sqrt{\kappa}$, it holds that

$$\begin{aligned} \left(\left(\sum_{k=1}^{N} \bar{y}_k - h_k\right) + \kappa\mu\right)^2 &= \left(\sum_{k=1}^{N} a_k \cdot \rho_k\right)^2 \\ &\leq \left(\sum_{k=1}^{N} a_k^2\right) \cdot \left(\sum_{k=1}^{N} \rho_k^2\right) \\ &= \left(\left(\sum_{k=1}^{N} (\bar{y}_k - h_k)^2\right) + \kappa\mu^2\right) \cdot (N+\kappa) \end{aligned}$$

By dividing through $N + \kappa$ we get the result.

**Sampling of the offset parameter**

For $b$ the integrand in (8) can be considered

$$
\begin{aligned}
b &\propto \exp\left(-\frac{\lambda}{2}\left((N+\kappa)b^2 - 2\left(\left(\sum_{k=1}^{N} \bar{y}_k - h_k\right) + \kappa\mu\right)b + \left(\sum_{k=1}^{N}(\bar{y}_k - h_k)^2\right) + \kappa\mu^2 + 2\beta\right)\right) \\
&\propto \exp\left(-\frac{\lambda(N+\kappa)}{2}\left(b - \frac{\left(\sum_{k=1}^{N} \bar{y}_k - h_k\right) + \kappa\mu}{N+\kappa}\right)^2\right) \\
&\propto \mathcal{N}\left(\mu' = \frac{\left(\sum_{k=1}^{N} \bar{y}_k - h_k\right) + \kappa\mu}{N+\kappa}, \hat{\lambda} = \lambda(N+\kappa)\right) \qquad (10)
\end{aligned}
$$

The distribution is well defined because $\mu' \in \mathbb{R}$ and $\lambda' \in \mathbb{R}_+$ by definition.

#### 2.3 Unknown noise, scaling and offset parameters

We consider the observation and noise model

$$\bar{y}_k = (s \cdot h_k + b) + \epsilon_k, \quad \epsilon_k \sim \mathcal{N}(0, 1/\lambda),$$

with unknown scaling parameter $s$, unknown offset $b$ and unknown noise parameter $\lambda$. For these unknown parameters we assume a joint Normal-Gamma prior,

$$
\begin{aligned}
p(s, b, \lambda) &= \mathcal{N}(s \mid z, 1/(\tau\lambda)) \cdot \mathcal{N}(b \mid \mu, 1/(\kappa\lambda)) \cdot \Gamma(\lambda \mid \alpha, \beta) \\
&= \frac{\beta^\alpha \sqrt{\kappa\tau}}{\Gamma(\alpha) 2\pi} \cdot \lambda^\alpha \cdot \exp\left(-\frac{\lambda}{2}(\kappa(b-\mu)^2 + \tau(s-z)^2 + 2\beta)\right)
\end{aligned}
$$

with $\mu$, $\alpha$, $\beta$, $\kappa$, $\tau$ and $z$ as hyper-parameters. $\Gamma(\cdot)$ denotes the Gamma function. The marginalized likelihood is denoted by

$$
\begin{aligned}
p(\mathcal{D} \mid \theta) &= \int_0^\infty \int_{-\infty}^\infty \int_{-\infty}^\infty p(\mathcal{D} \mid \theta, s, b, \lambda) p(s, b, \lambda) \,\mathrm{d}b \,\mathrm{d}s \,\mathrm{d}\lambda \\
&= \int_0^\infty \int_{-\infty}^\infty \int_{-\infty}^\infty \left(\frac{\lambda}{2\pi}\right)^{N/2} \exp\left(-\frac{\lambda}{2}\sum_{k=1}^{N}(\bar{y}_k - (s \cdot h_k + b))^2\right) \\
&\quad \cdot \frac{\beta^\alpha \sqrt{\kappa\tau}}{\Gamma(\alpha) 2\pi} \cdot \lambda^\alpha \cdot \exp\left(-\frac{\lambda}{2}(\kappa(b-\mu)^2 + \tau(s-z)^2 + 2\beta)\right) \mathrm{d}b \,\mathrm{d}s \,\mathrm{d}\lambda
\end{aligned}
$$

We reformulate the marginalized likelihood by removing constants from the integral and by reordering the terms under the integral. This yields

$$\begin{aligned}p(\mathcal{D} \mid \theta) =& \frac{\beta^\alpha \sqrt{\kappa\tau}}{\Gamma(\alpha)}(2\pi)^{-\frac{N+2}{2}} \int_0^\infty \lambda^{\frac{N}{2}+\alpha} \exp\left(-\frac{1}{2}\left(2\beta + \tau z^2 + k\mu^2 + \sum_{k=1}^N \bar{y}_k^2\right)\cdot\lambda\right) \\ &\cdot \int_{-\infty}^\infty \exp\left(-\frac{\lambda}{2}\left(\tau + \sum_{k=1}^N h_k^2\right)\cdot s^2 + \lambda\left(z\tau + \sum_{k=1}^N h_k\bar{y}_k\right)\cdot s\right) \\ &\cdot \int_{-\infty}^\infty \exp\left(-\frac{\lambda(N+\kappa)}{2}\cdot b^2 + \lambda\left(\kappa\mu + \sum_{k=1}^N \bar{y}_k - h_k\cdot s\right)\cdot b\right) \mathrm{d}b\,\mathrm{d}s\,\mathrm{d}\lambda\end{aligned}$$

Applying the Gauss integral (2) to integrate $b$, we get

$$\begin{aligned}p(\mathcal{D} \mid \theta) =& \frac{\beta^\alpha}{\Gamma(\alpha)(2\pi)^{\frac{N+1}{2}}} \cdot \sqrt{\frac{\kappa\tau}{N+\kappa}} \\ &\cdot \int_0^\infty \lambda^{\alpha+\frac{N+1}{2}-1} \exp\left(-\frac{\lambda}{2}\cdot\left(\kappa\mu^2 + \tau z^2 + 2\beta + \sum_{k=1}^N \bar{y}_k^2 - \frac{\left(\kappa\mu + \sum_{k=1}^N \bar{y}_k\right)^2}{N+\kappa}\right)\right) \\ &\cdot \int_{-\infty}^\infty \exp\left(-\left(\frac{\lambda}{2}\cdot\frac{(N+\kappa)\left(\tau + \sum_{k=1}^N h_k^2\right) - \left(\sum_{k=1}^N h_k\right)^2}{N+\kappa}\right)\cdot s^2\right) \\ &\cdot \exp\left(-\left(\lambda\cdot\frac{\left(\kappa\mu + \sum_{k=1}^N \bar{y}_k\right)\left(\sum_{k=1}^N h_k\right) - (N+\kappa)\left(\tau z + \sum_{k=1}^N h_k\bar{y}_k\right)}{N+\kappa}\right)\cdot s\right) \mathrm{d}s\,\mathrm{d}\lambda\end{aligned} \tag{11}$$

Following Cauchy–Schwarz inequality (7), it holds that

$$\left(\sum_{k=1}^N 1\cdot h_k\right)^2 \le N\cdot\sum_{k=1}^N h_k^2,$$

thus, $s$ can be integrated by applying (2) and we obtain

$$\begin{aligned}p(\mathcal{D} \mid \theta) =& \frac{\beta^\alpha}{\Gamma(\alpha)(2\pi)^{\frac{N}{2}}} \cdot \sqrt{\kappa\tau} \cdot \left((N+\kappa)\left(\tau + \sum_{k=1}^N h_k^2\right) - \left(\sum_{k=1}^N h_k\right)^2\right)^{-1/2} \\ &\cdot \int_0^\infty \lambda^{\alpha+\frac{N}{2}-1} \exp\left(-\lambda\cdot\omega\right)\,\mathrm{d}\lambda.\end{aligned} \tag{12}$$

with

$$\omega := \frac{1}{2} \cdot \left( \kappa\mu^2 + \tau z^2 + 2\beta + \sum_{k=1}^{N} \bar{y}_k^2 - \frac{\left(\kappa\mu + \sum_{k=1}^{N} \bar{y}_k\right)^2}{N+\kappa} \right) - \frac{1}{2} \frac{\left(\left(\kappa\mu + \sum_{k=1}^{N} \bar{y}_k\right)\left(\sum_{k=1}^{N} h_k\right) - (N+\kappa)\left(\tau z + \sum_{k=1}^{N} h_k \bar{y}_k\right)\right)^2}{(N+\kappa)\left((N+\kappa)\left(\tau + \sum_{k=1}^{N} h_k^2\right) - \left(\sum_{k=1}^{N} h_k\right)^2\right)}$$

Using the transformation $z = \lambda\omega$ and using the definition of the Gamma function (5), we obtain a closed-form solution for the marginalized likelihood

$$\begin{aligned} p(\mathcal{D} \mid \theta) =& \frac{\beta^\alpha}{\Gamma(\alpha)(2\pi)^{\frac{N}{2}}} \cdot \sqrt{\kappa\tau} \cdot \left( (N+\kappa)\left(\tau + \sum_{k=1}^{N} h_k^2\right) - \left(\sum_{k=1}^{N} h_k\right)^2 \right)^{-1/2} \\ &\cdot \omega^{-\left(\alpha + \frac{N}{2}\right)} \int_0^\infty z^{\left(\alpha + \frac{N}{2} - 1\right)} \exp(-z)\,\mathrm{d}z \\ =& \frac{(\beta/\omega)^\alpha}{\Gamma(\alpha)(2\pi\omega)^{\frac{N}{2}}} \cdot \sqrt{\kappa\tau} \cdot \left( (N+\kappa)\left(\tau + \sum_{k=1}^{N} h_k^2\right) - \left(\sum_{k=1}^{N} h_k\right)^2 \right)^{-1/2} \Gamma\left(\alpha + \frac{N}{2}\right) \end{aligned}$$

##### 2.3.1 Retrieving the marginalized parameters

The conditional probability of the marginalized noise, scaling and offset parameters can be derived after obtaining $p(\theta \mid \mathcal{D})$ given values of $\theta$.

##### Sampling of the noise parameter

The integrand in (12) shows that the precision $\lambda$ has (up to a constant) the distribution of a Gamma distribution, mainly

$$\lambda \propto \Gamma\left(\alpha' = \alpha + \frac{N}{2}, \beta' = \omega\right)$$

but with the necessary condition that $\omega > 0$.

To show the necessary condition for $\omega$ the marginalised integral has to be finite:

$$\begin{aligned} p(D \mid \theta) &= \int_0^\infty \int_{-\infty}^\infty \int_{-\infty}^\infty \underbrace{p(\bar{y}_k \mid \theta, s, b, \lambda)}_{\leq (\lambda/2\pi)^{N/2}} p(s,b,\lambda)\,\mathrm{d}b\,\mathrm{d}s\,\mathrm{d}\lambda \\ &\leq \left(\frac{\lambda}{2\pi}\right)^{N/2} \cdot \underbrace{\int_0^\infty \int_{-\infty}^\infty \int_{-\infty}^\infty p(s,b,\lambda)\,\mathrm{d}b\,\mathrm{d}s\,\mathrm{d}\lambda}_{=1} = \left(\frac{\lambda}{2\pi}\right)^{N/2} < \infty \end{aligned}$$

Then the integral $\int_0^\infty \lambda^{\alpha+\frac{N}{2}-1} \exp\left(-\lambda \cdot \omega\right) \mathrm{d}\lambda$ can be considered. If we can show that for $\omega \leq 0$ the integral is infinite we have a contradiction and receive the conclusion $\omega > 0$.

- $\omega = 0$
We deal with the integral $\int_0^\infty \lambda^{\alpha+\frac{N}{2}-1} \mathrm{d}\lambda$. Because $\alpha + N/2 > 0$ we have $\alpha + \frac{N}{2} - 1 \geq -1$ and therefore $\int_1^\infty \lambda^{\alpha+\frac{N}{2}-1} \mathrm{d}\lambda = \infty$.

- $\omega < 0, N \geq 2$
In this case we have $\lambda^{\alpha+\frac{N}{2}-1} \geq 1$ on $[1, \infty)$ and therefore $\int_1^\infty \lambda^{\alpha+\frac{N}{2}-1} \exp\left(-\lambda \cdot \omega\right) \mathrm{d}\lambda \geq \int_1^\infty \exp\left(-\lambda \cdot \omega\right) \mathrm{d}\lambda \geq \int_1^\infty 1 \,\mathrm{d}\lambda = \infty$

- $\omega < 0, N = 1$
For this case we refer to the first case as $\exp(-\lambda\omega) > 1$ on $[1, \infty)$. $\int_1^\infty \lambda^{\alpha+\frac{N}{2}-1} \exp\left(-\lambda \cdot \omega\right) \mathrm{d}\lambda \geq \int_1^\infty \lambda^{\alpha+\frac{N}{2}-1} \mathrm{d}\lambda = \infty$.

Therefore we can conclude that $\omega > 0$.

##### Sampling of the scaling parameter

For $s$ the integrand in (11) can be considered

$$
\begin{aligned}
s \propto \exp &\left(-\left(\frac{\lambda}{2} \cdot \frac{(N+\kappa)\left(\tau + \sum_{k=1}^N h_k^2\right) - \left(\sum_{k=1}^N h_k\right)^2}{N+\kappa}\right) \cdot s^2\right) \\
&\cdot \exp\left(-\left(\lambda \cdot \frac{\left(\kappa\mu + \sum_{k=1}^N \bar{y}_k\right)\left(\sum_{k=1}^N h_k\right) - (N+\kappa)\left(\tau z + \sum_{k=1}^N h_k \bar{y}_k\right)}{N+\kappa}\right) \cdot s\right)
\end{aligned}
$$

We define

$$
s \propto \exp\left(-\frac{\lambda}{2} \cdot \frac{\widehat{A}}{N+\kappa} \cdot s^2 - \lambda \cdot \frac{\widehat{B}}{N+\kappa} \cdot s\right)
$$

with

$$
\begin{aligned}
\widehat{A} &:= (N+\kappa)\left(\tau + \sum_{k=1}^N h_k^2\right) - \left(\sum_{k=1}^N h_k\right)^2 \\
\widehat{B} &:= \left(\kappa\mu + \sum_{k=1}^N \bar{y}_k\right)\left(\sum_{k=1}^N h_k\right) - (N+\kappa)\left(\tau z + \sum_{k=1}^N h_k \bar{y}_k\right)
\end{aligned}
$$

Then we obtain

$$
\begin{aligned}
s &\propto \exp\left(-\frac{\lambda}{2}\left(\frac{\widehat{A}}{N+\kappa} \cdot s^2 + \frac{2\widehat{B}}{N+\kappa} \cdot s\right)\right) \\
s &\propto \exp\left(-\frac{\lambda\widehat{A}}{2\,(N+\kappa)}\left(s^2 - \left(-\frac{\widehat{B}}{\widehat{A}}\right)\right)^2\right) \\
s &\propto \mathcal{N}\left(\mu' = -\frac{\widehat{B}}{\widehat{A}}, \lambda' = \frac{\lambda\widehat{A}}{(N+\kappa)}\right) \qquad (13)
\end{aligned}
$$

$\widehat{A} > 0$ holds because by Cauchy–Schwarz inequality $\left(\sum_{k=1}^{N} h_k \cdot 1\right)^2 \leq N \cdot \sum_{k=1}^{N} h_k^2$ and therefore $\widehat{A} \geq (N+\kappa)\tau + \kappa \sum_{k=1}^{N} h_k^2 > 0$.

##### Sampling of the offset parameter

For this, the previously obtained expression in (10) can be used, therefore

$$b \propto \mathcal{N}\left(\mu' = \frac{\left(\sum_{k=1}^{N} \bar{y}_k - s \cdot h_k\right) + \kappa\mu}{N+\kappa}, \hat{\lambda} = \lambda(N+\kappa)\right).$$

#### 2.4 Experimentally measured noise variance and unknown scaling parameters

In the case that the variance of the noise is known (i.e. measured), the scaling parameters $s$ still need to inferred from the experimental data. We present here the corresponding marginalization.

We consider the observation and noise model

$$\bar{y}_k = s \cdot h_k + \epsilon_k, \quad \epsilon_k \sim \mathcal{N}(0, 1/\lambda),$$

with unknown scaling parameters $s$ and experimentally measured $\lambda$. For these unknown parameters we assume a Gaussian prior,

$$p(s) = \frac{1}{\sqrt{2\pi\hat{\sigma}^2}} \cdot \exp\left(-\frac{1}{2}\frac{(s-\mu)^2}{\hat{\sigma}^2}\right)$$

with $\mu$ and $\hat{\sigma}^2$ as hyper-parameters. The marginalized likelihood is denoted by

$$\begin{aligned} p(\mathcal{D} \mid \theta) &= \int_{-\infty}^{\infty} p(\mathcal{D} \mid \theta, s) p(s) \, \mathrm{d}s \\ &= \int_{-\infty}^{\infty} \left(\frac{\lambda}{2\pi}\right)^{N/2} \exp\left(-\frac{\lambda}{2}\sum_{k=1}^{N}(\bar{y}_k - s \cdot h_k)^2\right) \frac{1}{\sqrt{2\pi\hat{\sigma}^2}} \cdot \exp\left(-\frac{1}{2}\frac{(s-\mu)^2}{\hat{\sigma}^2}\right) \mathrm{d}s. \end{aligned}$$

We reformulate the marginalized likelihood by removing constants from the integral and by reordering the terms under the integral. This yields

$$p(\mathcal{D} \mid \theta) = \left(\frac{\lambda}{2\pi}\right)^{N/2} \underbrace{\int_{-\infty}^{\infty} \frac{1}{\sqrt{2\pi\hat{\sigma}^2}} \cdot \exp\left(-\frac{\lambda}{2}\sum_{k=1}^{N}(\bar{y}_k - s \cdot h_k)^2\right) \cdot \exp\left(-\frac{1}{2}\frac{(s-\mu)^2}{\hat{\sigma}^2}\right) \mathrm{d}s}_{(*)}$$

The integral $(*)$ with respect to $s$ can be written as

$$\begin{aligned} (*) &= \frac{1}{\sqrt{2\pi\hat{\sigma}^2}} \int_{-\infty}^{\infty} \exp\left(-\frac{\lambda}{2}\left(s^2\sum_{k=1}^{N} h_k^2 - 2s\sum_{k=1}^{N}\bar{y}_k h_k + \sum_{k=1}^{N}\bar{y}_k^2 + \frac{s^2 - 2s\mu + \mu^2}{\lambda\hat{\sigma}^2}\right)\right) \mathrm{d}s \\ &= \frac{1}{\sqrt{2\pi\hat{\sigma}^2}} \int_{-\infty}^{\infty} \exp\left(-\frac{1}{2}\left(\lambda\sum_{k=1}^{N} h_k^2 + \frac{1}{\hat{\sigma}^2}\right)s^2 + \left(\lambda\sum_{k=1}^{N}\bar{y}_k h_k + \frac{\mu}{\hat{\sigma}^2}\right)s - \frac{\lambda}{2}\sum_{k=1}^{N}\bar{y}_k^2 - \frac{\mu^2}{2\hat{\sigma}^2}\right) \mathrm{d}s \end{aligned}$$

To simplify the notation, we define $\hat{\lambda} := \frac{1}{\hat{\sigma}^2}$ and we get

$$(*) = \left(\frac{\hat{\lambda}}{2\pi}\right)^{1/2} \int_{-\infty}^{\infty} \exp\left(-\frac{1}{2}\left(\lambda \sum_{k=1}^{N} h_k^2 + \hat{\lambda}\right) s^2 + \left(\lambda \sum_{k=1}^{N} \bar{y}_k h_k + \hat{\lambda}\mu\right) s - \frac{1}{2}\left(\lambda \sum_{k=1}^{N} \bar{y}_k^2 + \hat{\lambda}\mu^2\right)\right) \, \mathrm{d}s \tag{14}$$

and with constants

$$A := \frac{1}{2}\left(\lambda \sum_{k=1}^{N} h_k^2 + \hat{\lambda}\right), \qquad B := \left(\lambda \sum_{k=1}^{N} \bar{y}_k h_k + \hat{\lambda}\mu\right), \qquad C := -\frac{1}{2}\left(\lambda \sum_{k=1}^{N} \bar{y}_k^2 + \hat{\lambda}\mu^2\right)$$

The integral (14) can be solved using (2). We obtain

$$\begin{aligned}
(*) &= \left(\frac{\hat{\lambda}}{2\pi}\right)^{1/2} \sqrt{\frac{\pi}{\frac{1}{2}\left(\lambda \sum_{k=1}^{N} h_k^2 + \hat{\lambda}\right)}} \exp\left\{ \frac{\left(\lambda \sum_{k=1}^{N} \bar{y}_k h_k + \hat{\lambda}\mu\right)^2}{4\frac{1}{2}\left(\lambda \sum_{k=1}^{N} h_k^2 + \hat{\lambda}\right)} - \frac{1}{2}\left(\lambda \sum_{k=1}^{N} \bar{y}_k^2 + \hat{\lambda}\mu^2\right) \right\} \\
&= \left(\frac{\hat{\lambda}}{2\pi}\right)^{1/2} \sqrt{\frac{2\pi}{\left(\lambda \sum_{k=1}^{N} h_k^2 + \hat{\lambda}\right)}} \exp\left\{ \frac{\left(\lambda \sum_{k=1}^{N} \bar{y}_k h_k + \hat{\lambda}\mu\right)^2}{2\left(\lambda \sum_{k=1}^{N} h_k^2 + \hat{\lambda}\right)} - \frac{1}{2}\left(\lambda \sum_{k=1}^{N} \bar{y}_k^2 + \hat{\lambda}\mu^2\right) \right\} \\
&= \left(\frac{\hat{\lambda}}{\lambda \sum_{k=1}^{N} h_k^2 + \hat{\lambda}}\right)^{1/2} \exp\left\{ \frac{1}{2}\left( \frac{\left(\lambda \sum_{k=1}^{N} \bar{y}_k h_k + \hat{\lambda}\mu\right)^2}{\lambda \sum_{k=1}^{N} h_k^2 + \hat{\lambda}} - \lambda \sum_{k=1}^{N} \bar{y}_k^2 - \hat{\lambda}\mu^2 \right) \right\}
\end{aligned}$$

To simplify the notation, we define $\omega := \lambda \sum_{k=1}^{N} h_k^2 + \hat{\lambda}$ and we get

$$(*) = \left(\frac{\hat{\lambda}}{\omega}\right)^{1/2} \exp\left\{ \frac{1}{2}\left( \frac{\left(\lambda \sum_{k=1}^{N} \bar{y}_k h_k + \hat{\lambda}\mu\right)^2}{\omega} - \lambda \sum_{k=1}^{N} \bar{y}_k^2 - \hat{\lambda}\mu^2 \right) \right\}.$$

The substitution of this result for the integral over $s$ in the integral formulation yields a closed-form solution for the marginalized likelihood

$$p(\mathcal{D} \mid \theta) = \left(\frac{\lambda}{2\pi}\right)^{N/2} \left(\frac{\hat{\lambda}}{\omega}\right)^{1/2} \exp\left\{ \frac{1}{2}\left( \frac{\left(\lambda \sum_{k=1}^{N} \bar{y}_k h_k + \hat{\lambda}\mu\right)^2}{\omega} - \lambda \sum_{k=1}^{N} \bar{y}_k^2 - \hat{\lambda}\mu^2 \right) \right\}.$$

##### 2.4.1 Retrieving the marginalized parameters

The conditional probability of the marginalized scaling parameters can be derived after obtaining $p(\theta \mid \mathcal{D})$ given values of $\theta$.

##### Sampling of the scaling parameter

For $s$ the integrand in (14) can be considered

$$\begin{aligned}
s &\propto \exp\left(-\frac{1}{2}\left(\lambda\sum_{k=1}^{N}h_k^2+\hat{\lambda}\right)\left(s^2-\frac{2\left(\lambda\sum_{k=1}^{N}\bar{y}_k h_k+\hat{\lambda}\mu\right)}{\left(\lambda\sum_{k=1}^{N}h_k^2+\hat{\lambda}\right)}s\right)\right)\\
&\propto \exp\left(-\frac{1}{2}\frac{\left(s-\frac{\left(\lambda\sum_{k=1}^{N}\bar{y}_k h_k+\hat{\lambda}\mu\right)}{\left(\lambda\sum_{k=1}^{N}h_k^2+\hat{\lambda}\right)}\right)^2}{1/\left(\lambda\sum_{k=1}^{N}h_k^2+\hat{\lambda}\right)}\right)\\
&\propto \mathcal{N}\left(\mu'=\frac{\left(\lambda\sum_{k=1}^{N}\bar{y}_k h_k+\hat{\lambda}\mu\right)}{\left(\lambda\sum_{k=1}^{N}h_k^2+\hat{\lambda}\right)},(\sigma')^2=\left(\lambda\sum_{k=1}^{N}h_k^2+\hat{\lambda}\right)^{-1}\right)
\end{aligned}$$

#### 2.5 Experimentally measured noise variance and unknown offset parameters

In the case that the variance of the noise is known (i.e. measured), the offset parameters $b$ still need to inferred from the experimental data. We present here the corresponding marginalization.

We consider the observation and noise model

$$\bar{y}_k = h_k + b + \epsilon_k, \quad \epsilon_k \sim \mathcal{N}(0, 1/\lambda),$$

with unknown offset parameters $b$ and experimentally measured $\lambda$. For these unknown parameters we assume a Gaussian prior,

$$p(b) = \sqrt{\frac{\hat{\lambda}}{2\pi}} \cdot \exp\left(-\frac{\hat{\lambda}}{2}(b-\mu)^2\right)$$

with $\mu$ and $\hat{\lambda}$ as hyper-parameters. The marginalized likelihood is denoted by

$$\begin{aligned}
p(\mathcal{D} \mid \theta) &= \int_{-\infty}^{\infty} p(\mathcal{D} \mid \theta, b)p(b)\ \mathrm{d}b\\
&= \int_{-\infty}^{\infty}\left(\frac{\lambda}{2\pi}\right)^{N/2}\exp\left(-\frac{\lambda}{2}\sum_{k=1}^{N}(\bar{y}_k-(b+h_k))^2\right)\sqrt{\frac{\hat{\lambda}}{2\pi}}\cdot\exp\left(-\frac{\hat{\lambda}}{2}(b-\mu)^2\right)\mathrm{d}b\\
&= \left(\frac{\lambda}{2\pi}\right)^{N/2}\sqrt{\frac{\hat{\lambda}}{2\pi}}\underbrace{\int_{-\infty}^{\infty}\exp\left(-\frac{\lambda}{2}\sum_{k=1}^{N}((\bar{y}_k-h_k)-b)^2-\frac{\hat{\lambda}}{2}(b-\mu)^2\right)\mathrm{d}b}_{(*)}
\end{aligned}$$

The integral $(*)$ with respect to $b$ can be written as

$$
\begin{aligned}
(*) &= \int_{-\infty}^{\infty} \exp\left(-\frac{\lambda}{2}\sum_{k=1}^{N}((\bar{y}_k - h_k) - b)^2 - \frac{\hat{\lambda}}{2}(b-\mu)^2\right)\,\mathrm{d}b \\
&= \int_{-\infty}^{\infty} \exp\left(-\frac{1}{2}\left(\lambda N + \hat{\lambda}\right)b^2 + \left(\lambda\sum_{k=1}^{N}(\bar{y}_k - h_k) + \hat{\lambda}\mu\right)b - \frac{1}{2}\left(\lambda\sum_{k=1}^{N}\left(\bar{y}_k - h_k\right)^2 + \hat{\lambda}\mu^2\right)\right)\,\mathrm{d}b
\end{aligned} \tag{15}
$$

with constants

$$
A := \frac{1}{2}\left(\lambda N + \hat{\lambda}\right), \qquad B := \left(\lambda\sum_{k=1}^{N}(\bar{y}_k - h_k) + \hat{\lambda}\mu\right), \qquad C := -\frac{1}{2}\left(\lambda\sum_{k=1}^{N}\left(\bar{y}_k - h_k\right)^2 + \hat{\lambda}\mu^2\right).
$$

As $\lambda, \hat{\lambda}, N > 0$, $A > 0$ can be assured, we can apply (2) and we obtain

$$
\begin{aligned}
(*) &= \sqrt{\frac{\pi}{\frac{1}{2}\left(\lambda N + \hat{\lambda}\right)}} \cdot \exp\left(\frac{\left(\lambda\sum_{k=1}^{N}(\bar{y}_k - h_k) + \hat{\lambda}\mu\right)^2}{4\frac{1}{2}\left(\lambda N + \hat{\lambda}\right)} - \frac{1}{2}\left(\lambda\sum_{k=1}^{N}\left(\bar{y}_k - h_k\right)^2 + \hat{\lambda}\mu^2\right)\right) \\
&= \sqrt{\frac{2\pi}{\lambda N + \hat{\lambda}}} \cdot \exp\left(\frac{1}{2}\left(\frac{\left(\lambda\sum_{k=1}^{N}(\bar{y}_k - h_k) + \hat{\lambda}\mu\right)^2}{\lambda N + \hat{\lambda}} - \lambda\sum_{k=1}^{N}\left(\bar{y}_k - h_k\right)^2 - \hat{\lambda}\mu^2\right)\right)
\end{aligned}
$$

The substitution of this result for the integral over $b$ in the integral formulation yields a closed-form solution for the marginalized likelihood

$$
\begin{aligned}
p(D \mid \theta) &= \left(\frac{\lambda}{2\pi}\right)^{N/2}\sqrt{\frac{\hat{\lambda}}{2\pi}}\sqrt{\frac{2\pi}{\lambda N + \hat{\lambda}}} \cdot \exp\left(\frac{1}{2}\left(\frac{\left(\lambda\sum_{k=1}^{N}(\bar{y}_k - h_k) + \hat{\lambda}\mu\right)^2}{\lambda N + \hat{\lambda}} - \lambda\sum_{k=1}^{N}\left(\bar{y}_k - h_k\right)^2 - \hat{\lambda}\mu^2\right)\right) \\
&= \left(\frac{\lambda}{2\pi}\right)^{N/2}\sqrt{\frac{\hat{\lambda}}{\lambda N + \hat{\lambda}}} \cdot \exp\left(\frac{1}{2}\left(\frac{\left(\lambda\sum_{k=1}^{N}(\bar{y}_k - h_k) + \hat{\lambda}\mu\right)^2}{\lambda N + \hat{\lambda}} - \lambda\sum_{k=1}^{N}\left(\bar{y}_k - h_k\right)^2 - \hat{\lambda}\mu^2\right)\right)
\end{aligned}
$$

##### 2.5.1 Retrieving the marginalized parameters

The conditional probability of the marginalized offset parameters can be derived after obtaining $p(\theta \mid \mathcal{D})$ given values of $\theta$.

**Sampling of the offset parameter**

For $b$ the integrand in (15) can be considered

$$b \propto \exp\left(-\frac{1}{2}\left(\lambda N + \hat{\lambda}\right) b^2 + \left(\lambda \sum_{k=1}^{N} (\bar{y}_k - h_k) + \hat{\lambda}\mu\right) b\right)$$

$$\propto \exp\left(-\frac{1}{2} \frac{\left(b - \frac{\left(\lambda \sum_{k=1}^{N} (\bar{y}_k - h_k) + \hat{\lambda}\mu\right)}{\left(\lambda N + \hat{\lambda}\right)}\right)^2}{1/\left(\lambda N + \hat{\lambda}\right)}\right)$$

$$\propto \mathcal{N}\left(\mu' = \frac{\left(\lambda \sum_{k=1}^{N} (\bar{y}_k - h_k) + \hat{\lambda}\mu\right)}{\left(\lambda N + \hat{\lambda}\right)}, (\sigma')^2 = \left(\lambda N + \hat{\lambda}\right)^{-1}\right) \tag{16}$$

#### 2.6 Experimentally measured noise variance, unknown scaling and offset parameters

In the case that the variance of the noise is known (i.e. measured), the scaling $s$ and offset parameters $b$ still need to inferred from the experimental data. We present here the corresponding marginalization. We consider the observation and noise model

$$\bar{y}_k = (s \cdot h_k + b) + \epsilon_k, \quad \epsilon_k \sim \mathcal{N}(0, 1/\lambda),$$

with unknown scaling parameters $s$, offset parameters $b$ and experimentally measured $\lambda$. For these unknown parameters we assume a Gaussian prior,

$$p(s,b) = \sqrt{\frac{\tau}{2\pi}} \cdot \exp\left(-\frac{\tau}{2}(s-\nu)^2\right) \cdot \sqrt{\frac{\hat{\lambda}}{2\pi}} \cdot \exp\left(-\frac{\hat{\lambda}}{2}(b-\mu)^2\right)$$

with $\mu, \hat{\lambda}, \nu, \tau$ as hyper-parameters. The marginalized likelihood is denoted by

$$p(\mathcal{D} \mid \theta) = \int_{-\infty}^{\infty} \int_{-\infty}^{\infty} p(\mathcal{D} \mid \theta, s, b) p(s,b) \, \mathrm{d}b \, \mathrm{d}s$$
$$= \int_{-\infty}^{\infty} \int_{-\infty}^{\infty} \left(\frac{\lambda}{2\pi}\right)^{N/2} \exp\left(-\frac{\lambda}{2} \sum_{k=1}^{N} (\bar{y}_k - (b + s \cdot h_k))^2\right)$$
$$\cdot \sqrt{\frac{\hat{\lambda}}{2\pi}} \cdot \exp\left(-\frac{\hat{\lambda}}{2}(b-\mu)^2\right) \cdot \sqrt{\frac{\tau}{2\pi}} \cdot \exp\left(-\frac{\tau}{2}(s-\nu)^2\right) \mathrm{d}b \, \mathrm{d}s$$
$$= \left(\frac{\lambda}{2\pi}\right)^{N/2} \sqrt{\frac{\hat{\lambda}}{2\pi}} \sqrt{\frac{\tau}{2\pi}} \int_{-\infty}^{\infty} \int_{-\infty}^{\infty} \exp\left(-\frac{\lambda}{2} \sum_{k=1}^{N} ((\bar{y}_k - s \cdot h_k) - b)^2 - \frac{\hat{\lambda}}{2}(b-\mu)^2 - \frac{\tau}{2}(s-\nu)^2\right) \mathrm{d}b \, \mathrm{d}s$$

Applying the Gauss integral (2) to integrate $b$, we get

$$\begin{aligned}
p(\mathcal{D} \mid \theta) &= \left(\frac{\lambda}{2\pi}\right)^{N/2} \sqrt{\frac{\hat{\lambda}}{\lambda N + \hat{\lambda}}} \sqrt{\frac{\tau}{2\pi}} \\
&\quad \cdot \int_{-\infty}^{\infty} \exp\left(\frac{1}{2}\left(\frac{\left(\lambda \sum_{k=1}^{N}(\bar{y}_k - s \cdot h_k) + \hat{\lambda}\mu\right)^2}{\lambda N + \hat{\lambda}} - \lambda \sum_{k=1}^{N} (\bar{y}_k - s \cdot h_k)^2 - \hat{\lambda}\mu^2\right)\right) \cdot \exp\Big(-\frac{\tau}{2}(s-\nu)^2\Big) \,\mathrm{d}s \\
&= \left(\frac{\lambda}{2\pi}\right)^{N/2} \sqrt{\frac{\hat{\lambda}}{\lambda N + \hat{\lambda}}} \sqrt{\frac{\tau}{2\pi}} \\
&\quad \cdot \underbrace{\int_{-\infty}^{\infty} \exp\left(\frac{1}{2}\left(\frac{\left(\lambda \sum_{k=1}^{N}(\bar{y}_k - s \cdot h_k) + \hat{\lambda}\mu\right)^2}{\lambda N + \hat{\lambda}} - \lambda \sum_{k=1}^{N} (\bar{y}_k - s \cdot h_k)^2 - \hat{\lambda}\mu^2 - \tau(s-\nu)^2\right)\right) \mathrm{d}s}_{(*)}
\end{aligned}$$

The integral $(*)$ with respect to $s$ can be written as

$$\begin{aligned}
(*) = \int_{-\infty}^{\infty} \exp\Bigg(\frac{1}{2}\Bigg(&\frac{\lambda^2 \left(\sum_{k=1}^{N} \bar{y}_k - \sum_{k=1}^{N} h_k \cdot s\right)^2 + \hat{\lambda}^2\mu^2 + 2\hat{\lambda}\mu\lambda \sum_{k=1}^{N}(\bar{y}_k - s \cdot h_k)}{\lambda N + \hat{\lambda}} \\
&- \lambda \sum_{k=1}^{N} (\bar{y}_k - s \cdot h_k)^2 - \hat{\lambda}\mu^2 - \tau(s^2 - 2\nu s + \nu^2)\Bigg)\Bigg) \,\mathrm{d}s \\
= \int_{-\infty}^{\infty} \exp\Bigg(&-\frac{\lambda}{2}\left(\frac{\left(\lambda N + \hat{\lambda}\right)\left(\sum_{k=1}^{N} h_k^2 + \frac{\tau}{\lambda}\right) - \lambda \left(\sum_{k=1}^{N} h_k\right)^2}{\lambda N + \hat{\lambda}}\right) s^2 \\
&- \lambda \left(\frac{\left(\lambda \sum_{k=1}^{N} \bar{y}_k + \hat{\lambda}\mu\right)\left(\sum_{k=1}^{N} h_k\right) - \left(\lambda N + \hat{\lambda}\right)\left(\sum_{k=1}^{N} \bar{y}_k h_k + \frac{\tau}{\lambda}\nu\right)}{\lambda N + \hat{\lambda}}\right) s \\
&- \frac{1}{2}\left(\lambda \sum_{k=1}^{N} \bar{y}_k^2 + \hat{\lambda}\mu^2 + \tau\nu^2 - \frac{\lambda^2\left(\sum_{k=1}^{N} \bar{y}_k\right)^2 + \hat{\lambda}^2\mu^2 + 2\hat{\lambda}\mu\lambda \sum_{k=1}^{N} \bar{y}_k}{\lambda N + \hat{\lambda}}\right)\Bigg) \,\mathrm{d}s
\end{aligned}$$

with constants

$$\begin{aligned}
A &:= \frac{\lambda}{2}\left(\frac{\left(\lambda N + \hat{\lambda}\right)\left(\sum_{k=1}^{N} h_k^2 + \frac{\tau}{\lambda}\right) - \lambda\left(\sum_{k=1}^{N} h_k\right)^2}{\lambda N + \hat{\lambda}}\right), \\
B &:= -\lambda \left(\frac{\left(\lambda \sum_{k=1}^{N} \bar{y}_k + \hat{\lambda}\mu\right)\left(\sum_{k=1}^{N} h_k\right) - \left(\lambda N + \hat{\lambda}\right)\left(\sum_{k=1}^{N} \bar{y}_k h_k + \frac{\tau}{\lambda}\nu\right)}{\lambda N + \hat{\lambda}}\right), \text{ and} \\
C &:= -\frac{1}{2}\left(\lambda \sum_{k=1}^{N} \bar{y}_k^2 + \hat{\lambda}\mu^2 + \tau\nu^2 - \frac{\lambda^2\left(\sum_{k=1}^{N} \bar{y}_k\right)^2 + \hat{\lambda}^2\mu^2 + 2\hat{\lambda}\mu\lambda \sum_{k=1}^{N} \bar{y}_k}{\lambda N + \hat{\lambda}}\right).
\end{aligned}$$

Applying the Gauss integral (2) to integrate $s$, we get

$$
\begin{aligned}
(*) &= \sqrt{\frac{\pi}{\frac{\lambda}{2}\left(\frac{(\lambda N+\hat{\lambda})(\sum_{k=1}^{N} h_k^2+\frac{\tau}{\lambda})-\lambda(\sum_{k=1}^{N} h_k)^2}{\lambda N+\hat{\lambda}}\right)}} \\
&\cdot \exp\left(\frac{\lambda^2\left(\frac{(\lambda\sum_{k=1}^{N}\bar{y}_k+\hat{\lambda}\mu)(\sum_{k=1}^{N} h_k)-(\lambda N+\hat{\lambda})(\sum_{k=1}^{N}\bar{y}_k h_k+\frac{\tau}{\lambda}\nu)}{\lambda N+\hat{\lambda}}\right)^2}{4\frac{\lambda}{2}\left(\frac{(\lambda N+\hat{\lambda})(\sum_{k=1}^{N} h_k^2+\frac{\tau}{\lambda})-\lambda(\sum_{k=1}^{N} h_k)^2}{\lambda N+\hat{\lambda}}\right)}\right) \\
&\cdot \exp\left(-\frac{1}{2}\left(\lambda\sum_{k=1}^{N}\bar{y}_k^2+\hat{\lambda}\mu^2+\tau\nu^2-\frac{\lambda^2\left(\sum_{k=1}^{N}\bar{y}_k\right)^2+\hat{\lambda}^2\mu^2+2\hat{\lambda}\mu\lambda\sum_{k=1}^{N}\bar{y}_k}{\lambda N+\hat{\lambda}}\right)\right) \\
&= \sqrt{\frac{2\pi\left(\lambda N+\hat{\lambda}\right)}{\lambda\left(\left(\lambda N+\hat{\lambda}\right)\left(\sum_{k=1}^{N} h_k^2+\frac{\tau}{\lambda}\right)-\lambda\left(\sum_{k=1}^{N} h_k\right)^2\right)}} \\
&\cdot \exp\left(\frac{\lambda}{2}\cdot\frac{\left(\left(\lambda\sum_{k=1}^{N}\bar{y}_k+\hat{\lambda}\mu\right)\left(\sum_{k=1}^{N} h_k\right)-\left(\lambda N+\hat{\lambda}\right)\left(\sum_{k=1}^{N}\bar{y}_k h_k+\frac{\tau}{\lambda}\nu\right)\right)^2}{\left(\lambda N+\hat{\lambda}\right)\left(\left(\lambda N+\hat{\lambda}\right)\left(\sum_{k=1}^{N} h_k^2+\frac{\tau}{\lambda}\right)-\lambda\left(\sum_{k=1}^{N} h_k\right)^2\right)}\right) \\
&\cdot \exp\left(-\frac{1}{2}\left(\lambda\sum_{k=1}^{N}\bar{y}_k^2+\hat{\lambda}\mu^2+\tau\nu^2-\frac{\lambda^2\left(\sum_{k=1}^{N}\bar{y}_k\right)^2+\hat{\lambda}^2\mu^2+2\hat{\lambda}\mu\lambda\sum_{k=1}^{N}\bar{y}_k}{\lambda N+\hat{\lambda}}\right)\right)
\end{aligned}
$$

The substitution of this result for the integral over $s$ in the integral formulation yields a closed-form solution for the marginalized likelihood

$$
\begin{aligned}
p(\mathcal{D}\mid\theta) &= \left(\frac{\lambda}{2\pi}\right)^{N/2}\cdot\sqrt{\frac{\tau\hat{\lambda}}{\lambda\left(\left(\lambda N+\hat{\lambda}\right)\left(\sum_{k=1}^{N} h_k^2+\frac{\tau}{\lambda}\right)-\lambda\left(\sum_{k=1}^{N} h_k\right)^2\right)}} \\
&\cdot \exp\left(\frac{1}{2}\cdot\frac{\left(\left(\lambda\sum_{k=1}^{N}\bar{y}_k+\hat{\lambda}\mu\right)\left(\lambda\sum_{k=1}^{N} h_k\right)-\left(\lambda N+\hat{\lambda}\right)\left(\lambda\sum_{k=1}^{N}\bar{y}_k h_k+\tau\nu\right)\right)^2}{\left(\lambda N+\hat{\lambda}\right)\left(\left(\lambda N+\hat{\lambda}\right)\left(\lambda\sum_{k=1}^{N} h_k^2+\tau\right)-\left(\lambda\sum_{k=1}^{N} h_k\right)^2\right)}\right) \\
&\cdot \exp\left(-\frac{1}{2}\left(\lambda\sum_{k=1}^{N}\bar{y}_k^2+\hat{\lambda}\mu^2+\tau\nu^2-\frac{\left(\hat{\lambda}\mu+\lambda\sum_{k=1}^{N}\bar{y}_k\right)^2}{\lambda N+\hat{\lambda}}\right)\right)
\end{aligned}
$$

##### 2.6.1 Retrieving the marginalized parameters

The conditional probability of the marginalized scaling parameters can be derived after obtaining $p(\theta\mid\mathcal{D})$ given values of $\theta$.

###### Sampling of the scaling parameter

For $s$ the integrand in (11) can be considered and similarly to (13) we obtain

$$s \propto \mathcal{N}\left(\mu' = \frac{\left(\hat{\lambda} + \lambda N\right)\left(\tau\nu + \lambda \sum_{k=1}^{N} \bar{y}_k h_k\right) - \left(\hat{\lambda}\mu + \lambda \sum_{k=1}^{N} \bar{y}_k\right)\left(\lambda \sum_{k=1}^{N} h_k\right)}{\left(\hat{\lambda} + \lambda N\right)\left(\tau + \lambda \sum_{k=1}^{N} h_k^2\right) - \left(\lambda \sum_{k=1}^{N} h_k\right)^2},\right.$$
$$\left.(\sigma^2)' = \left(\tau + \lambda \sum_{k=1}^{N} h_k^2 - \frac{\left(\lambda \sum_{k=1}^{N} h_k\right)^2}{\hat{\lambda} + \lambda N}\right)^{-1}\right)$$

###### Sampling of the offset parameter

For this, the previously obtained expression in (16) can be used, therefore

$$b \propto \mathcal{N}\left(\mu' = \frac{\left(\lambda \sum_{k=1}^{N} (\bar{y}_k - h_k) + \hat{\lambda}\mu\right)}{\left(\lambda N + \hat{\lambda}\right)}, (\sigma')^2 = \left(\lambda N + \hat{\lambda}\right)^{-1}\right)$$

#### 2.7 Unknown noise (absolute measurement data)

We consider the observation and noise model

$$\bar{y}_k = h_k + \epsilon_k, \quad \epsilon_k \sim \mathcal{N}(0, 1/\lambda),$$

with unknown noise parameters $\lambda$. For these unknown parameters we assume a Gamma distribution as prior

$$p(\lambda) = \frac{\beta^\alpha}{\Gamma(\alpha)} \lambda^{\alpha-1} \exp(-\beta\lambda)$$

with $\alpha, \beta$ as hyper-parameters. $\Gamma(\cdot)$ denotes the Gamma function. The marginalized likelihood is denoted by

$$\begin{aligned} p(\mathcal{D} \mid \theta) &= \int_0^\infty p(\mathcal{D} \mid \theta, \lambda) p(\lambda) \; \mathrm{d}\lambda \\ &= \int_0^\infty \left(\frac{\lambda}{2\pi}\right)^{N/2} \exp\left(-\frac{\lambda}{2} \sum_{k=1}^{N} (\bar{y}_k - h_k)^2\right) \frac{\beta^\alpha}{\Gamma(\alpha)} \lambda^{\alpha-1} \exp(-\beta\lambda) \; \mathrm{d}\lambda \end{aligned}$$

We reformulate the marginalized likelihood by removing constants from the integral and by reordering the terms under the integral. This yields

$$\begin{aligned} p(\mathcal{D} \mid \theta) &= \left(\frac{1}{2\pi}\right)^{N/2} \frac{\beta^\alpha}{\Gamma(\alpha)} \int_0^\infty \lambda^{(N/2)+\alpha-1} \exp\left(-\lambda\left(\beta + \frac{1}{2} \sum_{k=1}^{N} (\bar{y}_k - h_k)^2\right)\right) \; \mathrm{d}\lambda \\ &= \left(\frac{1}{2\pi}\right)^{N/2} \frac{\beta^\alpha}{\Gamma(\alpha)} \int_0^\infty \lambda^{(N/2)+\alpha-1} \exp(-\lambda\omega) \; \mathrm{d}\lambda \end{aligned} \tag{17}$$

with

$$\omega := \beta + \frac{1}{2}\sum_{k=1}^{N}\left(\bar{y}_k - h_k\right)^2.$$

Using the transformation $z = \lambda\omega$, we obtain for the marginalized likelihood

$$\begin{aligned}
p(\mathcal{D} \mid \theta) &= \left(\frac{1}{2\pi}\right)^{N/2} \frac{\beta^\alpha}{\Gamma(\alpha)} \int_0^\infty \left(\frac{z}{\omega}\right)^{(N/2)+\alpha-1} \exp\left(-z\right) \frac{1}{\omega}\,\mathrm{d}z \\
&= \left(\frac{1}{2\pi}\right)^{N/2} \frac{\beta^\alpha}{\Gamma(\alpha)} \left(\frac{1}{\omega}\right)^{\delta} \int_0^\infty z^{\delta-1} \exp\left(-z\right)\ \mathrm{d}z \qquad (18)
\end{aligned}$$

with $\delta := \alpha + N/2$. Using the definition of the Gamma function (5), we obtain a closed-form solution for the marginalized likelihood

$$\begin{aligned}
p(\mathcal{D} \mid \theta) &= \left(\frac{1}{2\pi}\right)^{N/2} \frac{\beta^\alpha}{\Gamma(\alpha)} \left(\frac{1}{\omega}\right)^{\alpha+N/2} \cdot \Gamma\left(\alpha + \frac{N}{2}\right) \\
&= \frac{(\beta/\omega)^\alpha}{\Gamma(\alpha)(2\pi\omega)^{N/2}} \cdot \Gamma\left(\alpha + \frac{N}{2}\right).
\end{aligned}$$

##### 2.7.1 Retrieving the marginalized parameters

The conditional probability of the marginalized noise parameters can be derived after obtaining $p(\theta \mid \mathcal{D})$ given values of $\theta$.

##### Sampling of the noise parameter

The integrand in (18) shows that the precision $\lambda$ has (up to a constant) the distribution of a Gamma distribution, mainly

$$\lambda \propto \text{Gamma}(\alpha' = \alpha + \frac{N}{2}, \beta' = \omega).$$

Similarly

$$\sigma^2 \propto \text{Inv-Gamma}(\alpha' = \alpha + \frac{N}{2}, \beta' = \omega).$$

### 3 Marginalization-based approach for multiplicative log-normal noise

In this section, we consider *multiplicative log-normal* measurement noise. The data set $\mathcal{D}$ is a collection of measurements

$$\bar{y}_k = (s \cdot h_k) \cdot \epsilon_k, \quad \epsilon_k \sim \mathcal{N}(0, \sigma^2), k = 1, \ldots, N.$$

For simplicity of notation, we write the problem in terms of precision $\lambda$, which is the inverse of the variance $\sigma^2$.

Bayes' theorem for unknown model parameter vector $\theta$, unknown scaling factor $s$ and unknown noise parameter $\lambda$ is given by

$$p(\theta, s, \lambda \mid \mathcal{D}) = \frac{p(\mathcal{D} \mid \theta, s, \lambda) p(\theta, s, \lambda)}{p(\mathcal{D})},$$

with prior $p(\theta, s, \lambda)$, marginal $p(\mathcal{D})$ and likelihood

$$p(\mathcal{D} \mid \theta, s, \lambda) = \left( \prod_{k=1}^{N} \frac{1}{\bar{y}_k} \right) \left( \frac{\lambda}{2\pi} \right)^{N/2} \exp\left( -\frac{\lambda}{2} \sum_{k=1}^{N} (\log(\bar{y}_k) - \log(s \cdot h_k))^2 \right)$$

In the following, we show different combinations of known and unknown parameters.

`Note`: We marginalise with respect to the logarithm of the scaling ($s_{\log}$) and noise parameter ($\lambda$) to obtain an analytical solution.

#### 3.1 Unknown noise and scaling parameters

We consider the observation and noise model

$$\bar{y}_k = e^{s_{\log}} \cdot h_k \cdot \epsilon_k, \quad \epsilon_k \sim \mathcal{N}(0, 1/\lambda),$$

with unknown scaling parameter (in logarithmic scale) $s_{\log}$ and unknown noise parameter $\lambda$. For these unknown parameters we assume a Normal-Gamma prior,

$$\begin{aligned} p(s_{\log}, \lambda) &= \mathcal{N}(s_{\log} \mid \mu, 1/(\kappa\lambda)) \cdot \Gamma(\lambda \mid \alpha, \beta) \\ &= \frac{\beta^\alpha \sqrt{\kappa}}{\Gamma(\alpha)\sqrt{2\pi}} \lambda^{\alpha - 1/2} \exp\left( -\beta\lambda - \frac{\kappa\lambda(s_{\log} - \mu)^2}{2} \right) \end{aligned}$$

with $\mu$, $\alpha$, $\beta$, $\kappa$ as hyper-parameters. $\Gamma(\cdot)$ denotes the Gamma function. The marginalized likelihood is denoted by

$$\begin{aligned} p(\mathcal{D} \mid \theta) &= \int_0^\infty \int_{-\infty}^\infty p(\mathcal{D} \mid \theta, e^{s_{\log}}, \lambda) p(s_{\log}, \lambda) \, \mathrm{d}s_{\log} \, \mathrm{d}\lambda \\ &= \int_0^\infty \int_{-\infty}^\infty \left( \prod_{k=1}^{N} \frac{1}{\bar{y}_k} \right) \left( \frac{\lambda}{2\pi} \right)^{N/2} \exp\left( -\frac{\lambda}{2} \sum_{k=1}^{N} (\log(\bar{y}_k) - \log(e^{s_{\log}} \cdot h_k))^2 \right) \\ &\quad \cdot \frac{\beta^\alpha \sqrt{\kappa}}{\Gamma(\alpha)\sqrt{2\pi}} \lambda^{\alpha - 1/2} \exp\left( -\beta\lambda - \frac{\kappa\lambda(s_{\log} - \mu)^2}{2} \right) \, \mathrm{d}s_{\log} \, \mathrm{d}\lambda \end{aligned}$$

We reformulate the marginalized likelihood by removing constants from the integral and by reordering the terms under the integral. This yields

$$p(\mathcal{D} \mid \theta) = \left(\prod_{k=1}^{N} \frac{1}{\bar{y}_k}\right) \frac{\beta^{\alpha}\sqrt{\kappa}}{\Gamma(\alpha)(2\pi)^{(N+1)/2}} \int_0^{\infty} \lambda^{\alpha+(N-1)/2} \exp\left(-\beta\lambda\right) \\ \underbrace{\cdot \int_{-\infty}^{\infty} \exp\left(-\frac{\lambda}{2}\left(\sum_{k=1}^{N}(\log\left(\bar{y}_k/h_k\right) - s_{\log})^2 + \kappa(s_{\log} - \mu)^2\right)\right) \, \mathrm{d}s_{\log} \, \mathrm{d}\lambda}_{(*)}$$

The integral $(*)$ with respect to $s_{\log}$ can be written as

$$\begin{aligned}(*) &= \int_{-\infty}^{\infty} \exp\left(-\frac{\lambda}{2}\left(N s_{\log}^2 - 2 s_{\log}\sum_{k=1}^{N}\log\left(\bar{y}_k/h_k\right) + \sum_{k=1}^{N}\left(\log\left(\bar{y}_k/h_k\right)\right)^2 + \kappa s_{\log}^2 - 2\kappa\mu s_{\log} + \kappa\mu^2\right)\right) \, \mathrm{d}s_{\log} \\ &= \int_{-\infty}^{\infty} \exp\left(-\frac{\lambda}{2}\left(\kappa + N\right) s_{\log}^2 + \lambda\left(\sum_{k=1}^{N}\log\left(\bar{y}_k/h_k\right) + \kappa\mu\right) s_{\log} - \frac{\lambda}{2}\left(\sum_{k=1}^{N}\left(\log\left(\bar{y}_k/h_k\right)\right)^2 + \kappa\mu^2\right)\right) \, \mathrm{d}s_{\log}.\end{aligned} \tag{19}$$

This follows the structure of the Gaussian integral (2) with constants

$$A := \frac{\lambda}{2}\left(\kappa + N\right), \qquad B := \lambda\left(\sum_{k=1}^{N}\log\left(\bar{y}_k/h_k\right) + \kappa\mu\right), \qquad \text{and} \qquad C := -\frac{\lambda}{2}\left(\sum_{k=1}^{N}\left(\log\left(\bar{y}_k/h_k\right)\right)^2 + \kappa\mu^2\right).$$

As $\lambda, N, \kappa > 0, A > 0$ can be assured and we obtain

$$\begin{aligned}(*) &= \sqrt{\frac{\pi}{\frac{\lambda}{2}\left(\kappa + N\right)}} \exp\left(\frac{\left(\lambda\left(\sum_{k=1}^{N}\log\left(\bar{y}_k/h_k\right) + \kappa\mu\right)\right)^2}{4\frac{\lambda}{2}\left(\kappa + N\right)} - \frac{\lambda}{2}\left(\sum_{k=1}^{N}\left(\log\left(\bar{y}_k/h_k\right)\right)^2 + \kappa\mu^2\right)\right) \\ &= \sqrt{\frac{2\pi}{\lambda\left(\kappa + N\right)}} \exp\left(-\lambda\tilde{\omega}\right)\end{aligned}$$

with

$$\tilde{\omega} := -\frac{1}{2}\left(\frac{\left(\sum_{k=1}^{N}\log\left(\bar{y}_k/h_k\right) + \kappa\mu\right)^2}{\kappa + N} - \left(\sum_{k=1}^{N}\left(\log\left(\bar{y}_k/h_k\right)\right)^2 + \kappa\mu^2\right)\right).$$

The substitution of this result for the integral over $s_{\log}$ in the integral formulation yields

$$\begin{aligned}p(\mathcal{D} \mid \theta) &= \left(\prod_{k=1}^{N} \frac{1}{\bar{y}_k}\right) \frac{\beta^{\alpha}\sqrt{\kappa}}{\Gamma(\alpha)(2\pi)^{(N+1)/2}} \int_0^{\infty} \lambda^{\alpha+(N-1)/2} \exp\left(-\beta\lambda\right) \sqrt{\frac{2\pi}{\lambda\left(\kappa + N\right)}} \exp\left(-\lambda\tilde{\omega}\right) \, \mathrm{d}\lambda \\ &= \left(\prod_{k=1}^{N} \frac{1}{\bar{y}_k}\right) \frac{\beta^{\alpha}\sqrt{\kappa}}{\Gamma(\alpha)(2\pi)^{(N+1)/2}} \sqrt{\frac{2\pi}{\kappa + N}} \int_0^{\infty} \lambda^{\alpha+N/2-1} \exp\left(-\lambda\omega\right) \, \mathrm{d}\lambda\end{aligned} \tag{20}$$

with

$$\omega := \beta + \tilde{\omega} = \beta - \frac{1}{2}\left(\frac{\left(\sum_{k=1}^{N} \log\left(\bar{y}_k/h_k\right) + \kappa\mu\right)^2}{\kappa + N} - \left(\sum_{k=1}^{N} \left(\log\left(\bar{y}_k/h_k\right)\right)^2 + \kappa\mu^2\right)\right).$$

Using the transformation $z = \lambda\omega$, we obtain for the marginalized likelihood

$$\begin{aligned} p(\mathcal{D} \mid \theta) &= \left(\prod_{k=1}^{N} \frac{1}{\bar{y}_k}\right) \frac{\beta^\alpha \sqrt{\kappa}}{\Gamma(\alpha)(2\pi)^{(N+1)/2}} \sqrt{\frac{2\pi}{\kappa+N}} \int_0^\infty \left(\frac{z}{\omega}\right)^{\alpha+N/2-1} \exp\left(-z\right) \frac{1}{\omega} \,\mathrm{d}z \\ &= \left(\prod_{k=1}^{N} \frac{1}{\bar{y}_k}\right) \frac{\beta^\alpha \sqrt{\kappa}}{\Gamma(\alpha)(2\pi)^{(N+1)/2}} \sqrt{\frac{2\pi}{\kappa+N}} \cdot \left(\frac{1}{\omega}\right)^\delta \int_0^\infty z^{\delta-1} \exp\left(-z\right) \,\mathrm{d}z \end{aligned}$$

with $\delta := \alpha + N/2$. This integral can be solved analytically since it follows the structure of (5) yielding a closed-form solution for the marginalized likelihood,

$$\begin{aligned} p(\mathcal{D} \mid \theta) &= \left(\prod_{k=1}^{N} \frac{1}{\bar{y}_k}\right) \frac{\beta^\alpha \sqrt{\kappa}}{\Gamma(\alpha)(2\pi)^{(N+1)/2}} \sqrt{\frac{2\pi}{\kappa+N}} \cdot \left(\frac{1}{\omega}\right)^\delta \cdot \Gamma(\delta) \\ &= \left(\prod_{k=1}^{N} \frac{1}{\bar{y}_k}\right) \frac{\beta^\alpha \sqrt{\kappa}}{\Gamma(\alpha)(2\pi)^{(N+1)/2}} \sqrt{\frac{2\pi}{\kappa+N}} \cdot \left(\frac{1}{\omega}\right)^{\alpha+\frac{N}{2}} \cdot \Gamma\left(\alpha + \frac{N}{2}\right) \\ &= \left(\prod_{k=1}^{N} \frac{1}{\bar{y}_k}\right) \frac{(\beta/\omega)^\alpha}{\Gamma(\alpha)(2\pi\omega)^{N/2}} \cdot \sqrt{\frac{\kappa}{\kappa+N}} \cdot \Gamma\left(\alpha + \frac{N}{2}\right). \end{aligned}$$

##### 3.1.1 Retrieving the marginalized parameters

The conditional probability of the marginalized noise and scaling parameters can be derived after obtaining $p(\theta \mid \mathcal{D})$ given values of $\theta$.

###### Sampling of the noise parameter

The integrand in (20) shows that the precision $\lambda$ has (up to a constant) the distribution of a Gamma distribution, mainly

$$\lambda \propto \text{Gamma}(\alpha' = \alpha + N/2, \beta' = \omega).$$

Similarly

$$\sigma^2 \propto \text{Inv-Gamma}(\alpha' = \alpha + N/2, \beta' = \omega).$$

##### Sampling of the scaling parameter

For $s_{\log}$ the integrand in (19) can be considered

$$\begin{aligned}
s_{\log} &\propto \exp\left(-\frac{\lambda}{2}(\kappa+N)\left(s_{\log}^2 - \frac{2\left(\sum_{k=1}^N \log(\bar{y}_k/h_k)+\kappa\mu\right)}{\kappa+N}s_{\log}\right)\right)\\
&\propto \exp\left(-\frac{1}{2}\frac{\left(s_{\log}-\frac{\left(\sum_{k=1}^N \log(\bar{y}_k/h_k)+\kappa\mu\right)}{\kappa+N}\right)^2}{\sigma^2/(\kappa+N)}\right)\\
&\propto \mathcal{N}\left(\mu' = \frac{\left(\sum_{k=1}^N \log(\bar{y}_k/h_k)+\kappa\mu\right)}{\kappa+N}, (\sigma')^2 = \frac{\sigma^2}{\kappa+N}\right)
\end{aligned}$$

#### 3.2 Experimentally measured noise variance and unknown scaling parameters

In the case that the variance of the noise is known (i.e. measured), the scaling parameters $s$ still need to inferred from the experimental data. We present here the corresponding marginalization.

We consider the observation and noise model

$$\bar{y}_k = e^{s_{\log}} \cdot h_k + \epsilon_k, \quad \epsilon_k \sim \mathcal{N}(0, 1/\lambda),$$

with unknown scaling parameters (in logarithmic scale) $s_{\log}$ and experimentally measured $\lambda$. For these unknown parameters we assume a Gaussian prior,

$$p(s_{\log}) = \sqrt{\frac{\hat{\lambda}}{2\pi}} \cdot \exp\left(-\frac{\hat{\lambda}}{2}(s_{\log}-\mu)^2\right)$$

with $\mu, \hat{\lambda}$ as hyper-parameters. The marginalized likelihood is denoted by

$$\begin{aligned}
p(\mathcal{D} \mid \theta) &= \int_{-\infty}^{\infty} p(\mathcal{D} \mid \theta, e^{s_{\log}}) p(s_{\log})\ \mathrm{d}s_{\log}\\
&= \int_{-\infty}^{\infty} \left(\prod_{k=1}^N \frac{1}{\bar{y}_k}\right)\left(\frac{\lambda}{2\pi}\right)^{N/2} \exp\left(-\frac{\lambda}{2}\sum_{k=1}^N (\log(\bar{y}_k) - \log(e^{s_{\log}} \cdot h_k))^2\right)\\
&\cdot \sqrt{\frac{\hat{\lambda}}{2\pi}} \cdot \exp\left(-\frac{\hat{\lambda}}{2}(s_{\log}-\mu)^2\right)\ \mathrm{d}s_{\log}
\end{aligned}$$

We reformulate the marginalized likelihood by removing constants from the integral and by reordering the terms under the integral. This yields

$$p(\mathcal{D} \mid \theta) = \left(\prod_{k=1}^{N} \frac{1}{\bar{y}_k}\right) \left(\frac{\lambda}{2\pi}\right)^{N/2} \sqrt{\frac{\hat{\lambda}}{2\pi}} \underbrace{\int_{-\infty}^{\infty} \exp\left(-\frac{\lambda}{2}\sum_{k=1}^{N}(\log(\bar{y}_k/h_k) - s_{\log})^2 - \frac{\hat{\lambda}}{2}(s_{\log} - \mu)^2\right) \, \mathrm{d}s_{\log}}_{(*)}$$

The integral $(*)$ with respect to $s_{\log}$ can be written as

$$\begin{aligned}(*) &= \int_{-\infty}^{\infty} \exp\left(-\frac{\lambda}{2}\sum_{k=1}^{N}(\log(\bar{y}_k/h_k) - s_{\log})^2 - \frac{\hat{\lambda}}{2}(s_{\log} - \mu)^2\right) \, \mathrm{d}s_{\log} \\ &= \int_{-\infty}^{\infty} \exp\left(-\left(\frac{\lambda N + \hat{\lambda}}{2}\right) s_{\log}^2 + \left(\lambda \sum_{k=1}^{N} \log(\bar{y}_k/h_k) + \hat{\lambda}\mu\right) s_{\log} - \frac{\lambda}{2}\sum_{k=1}^{N}(\log(\bar{y}_k/h_k))^2 - \frac{\hat{\lambda}}{2}\mu^2\right) \, \mathrm{d}s_{\log} \end{aligned} \tag{21}$$

This integral can be solved analytically since it follows the structure of the Gaussian integral (2) with constants

$$A := \frac{\lambda N + \hat{\lambda}}{2}, \qquad B := \lambda \sum_{k=1}^{N} \log(\bar{y}_k/h_k) + \hat{\lambda}\mu, \qquad \text{and} \qquad C := -\frac{\lambda}{2}\sum_{k=1}^{N}(\log(\bar{y}_k/h_k))^2 - \frac{\hat{\lambda}}{2}\mu^2.$$

As $\lambda, N, \hat{\lambda} > 0, A > 0$ can be assured and we obtain

$$\begin{aligned}(*) &= \sqrt{\frac{\pi}{\frac{\lambda N + \hat{\lambda}}{2}}} \exp\left(\frac{\left(\lambda \sum_{k=1}^{N} \log(\bar{y}_k/h_k) + \hat{\lambda}\mu\right)^2}{4\frac{\lambda N + \hat{\lambda}}{2}} - \frac{\lambda}{2}\sum_{k=1}^{N}(\log(\bar{y}_k/h_k))^2 - \frac{\hat{\lambda}}{2}\mu^2\right) \\ &= \sqrt{\frac{2\pi}{\lambda N + \hat{\lambda}}} \exp\left(\frac{\left(\lambda \sum_{k=1}^{N} \log(\bar{y}_k/h_k) + \hat{\lambda}\mu\right)^2}{2\left(\lambda N + \hat{\lambda}\right)} - \frac{\lambda}{2}\sum_{k=1}^{N}(\log(\bar{y}_k/h_k))^2 - \frac{\hat{\lambda}}{2}\mu^2\right)\end{aligned}$$

The substitution of this result for the integral over $s_{\log}$ in the integral formulation yields a closed-form solution for the marginalized likelihood

$$\begin{aligned}p(\mathcal{D} \mid \theta) = &\left(\prod_{k=1}^{N} \frac{1}{\bar{y}_k}\right) \left(\frac{\lambda}{2\pi}\right)^{N/2} \sqrt{\frac{\hat{\lambda}}{\lambda N + \hat{\lambda}}} \\ &\cdot \exp\left(\frac{1}{2}\left(\frac{\left(\lambda \sum_{k=1}^{N} \log(\bar{y}_k/h_k) + \hat{\lambda}\mu\right)^2}{\lambda N + \hat{\lambda}} - \lambda \sum_{k=1}^{N}(\log(\bar{y}_k/h_k))^2 - \hat{\lambda}\mu^2\right)\right)\end{aligned}$$

##### 3.2.1 Retrieving the marginalized parameters

**Sampling of the scaling parameter**

For $s$ the integrand in (21) can be considered

$$\begin{aligned} s_{\log} &\propto \exp\left(-\left(\frac{\lambda N+\hat{\lambda}}{2}\right)s_{\log}^2+\left(\lambda\sum_{k=1}^{N}\log\left(\bar{y}_k/h_k\right)+\hat{\lambda}\mu\right)s_{\log}\right)\\ &\propto \exp\left(-\frac{1}{2}\frac{\left(s_{\log}-\frac{\left(\lambda\sum_{k=1}^{N}\log(\bar{y}_k/h_k)+\hat{\lambda}\mu\right)}{\left(\lambda N+\hat{\lambda}\right)}\right)^2}{1/\left(\lambda N+\hat{\lambda}\right)}\right)\\ &\propto \mathcal{N}\left(\mu'=\frac{\left(\lambda\sum_{k=1}^{N}\log\left(\bar{y}_k/h_k\right)+\hat{\lambda}\mu\right)}{\left(\lambda N+\hat{\lambda}\right)},(\sigma')^2=\left(\lambda N+\hat{\lambda}\right)^{-1}\right)\end{aligned}$$

#### 3.3 Unknown noise (absolute measurement data)

We consider the observation and noise model

$$\bar{y}_k = h_k\cdot\epsilon_k,\quad \epsilon_k\sim\mathcal{N}(0,1/\lambda),$$

with unknown noise parameters $\lambda$. For these unknown parameters we assume a Gamma distribution as prior

$$p(\lambda)=\frac{\beta^\alpha}{\Gamma(\alpha)}\lambda^{\alpha-1}\exp\left(-\beta\lambda\right)$$

with $\alpha,\beta$ as hyper-parameters. $\Gamma(\cdot)$ denotes the Gamma function. The marginalized likelihood is denoted by

$$p(\mathcal{D}\mid\theta)=\int_0^\infty\left(\prod_{k=1}^{N}\frac{1}{\bar{y}_k}\right)\left(\frac{\lambda}{2\pi}\right)^{N/2}\exp\left(-\frac{\lambda}{2}\sum_{k=1}^{N}(\log\left(\bar{y}_k\right)-\log\left(h_k\right))^2\right)\cdot\frac{\beta^\alpha}{\Gamma(\alpha)}\lambda^{\alpha-1}\cdot\exp\left(-\beta\lambda\right)\ \mathrm{d}\lambda$$

We reformulate the marginalized likelihood by removing constants from the integral and by reordering the terms under the integral. This yields

$$\begin{aligned} p(\mathcal{D}\mid\theta)&=\left(\prod_{k=1}^{N}\frac{1}{\bar{y}_k}\right)\left(\frac{1}{2\pi}\right)^{N/2}\frac{\beta^\alpha}{\Gamma(\alpha)}\int_0^\infty\lambda^{(N/2)+\alpha-1}\exp\left(-\lambda\left(\beta+\frac{1}{2}\sum_{k=1}^{N}(\log\left(\bar{y}_k\right)-\log\left(h_k\right))^2\right)\right)\ \mathrm{d}\lambda\\ &=\left(\prod_{k=1}^{N}\frac{1}{\bar{y}_k}\right)\left(\frac{1}{2\pi}\right)^{N/2}\frac{\beta^\alpha}{\Gamma(\alpha)}\int_0^\infty\lambda^{(N/2)+\alpha-1}\exp\left(-\lambda\omega\right)\ \mathrm{d}\lambda \qquad (22)\end{aligned}$$

with

$$\omega:=\beta+\frac{1}{2}\sum_{k=1}^{N}(\log\left(\bar{y}_k\right)-\log\left(h_k\right))^2.$$

Using the transformation $z = \lambda\omega$, we obtain for the marginalized likelihood

$$
\begin{aligned}
p(\mathcal{D} \mid \theta) &= \left(\prod_{k=1}^{N} \frac{1}{\bar{y}_k}\right) \left(\frac{1}{2\pi}\right)^{N/2} \frac{\beta^\alpha}{\Gamma(\alpha)} \int_0^\infty \left(\frac{z}{\omega}\right)^{(N/2)+\alpha-1} \exp\left(-z\right) \frac{1}{\omega}\, \mathrm{d}z \\
&= \left(\prod_{k=1}^{N} \frac{1}{\bar{y}_k}\right) \left(\frac{1}{2\pi}\right)^{N/2} \frac{\beta^\alpha}{\Gamma(\alpha)} \left(\frac{1}{\omega}\right)^{\delta} \int_0^\infty z^{\delta-1} \exp\left(-z\right)\, \mathrm{d}z
\end{aligned}
$$

with $\delta := \alpha + N/2$. This integral can be solved analytically since it follows the structure of (5) yielding a closed-form solution for the marginalized likelihood,

$$
\begin{aligned}
p(\mathcal{D} \mid \theta) &= \left(\prod_{k=1}^{N} \frac{1}{\bar{y}_k}\right) \left(\frac{1}{2\pi}\right)^{N/2} \frac{\beta^\alpha}{\Gamma(\alpha)} \left(\frac{1}{\omega}\right)^{\alpha+N/2} \cdot \Gamma\left(\alpha + \frac{N}{2}\right) \\
&= \left(\prod_{k=1}^{N} \frac{1}{\bar{y}_k}\right) \frac{(\beta/\omega)^\alpha}{\Gamma(\alpha)(2\pi\omega)^{N/2}} \cdot \Gamma\left(\alpha + \frac{N}{2}\right).
\end{aligned}
$$

##### 3.3.1 Retrieving the marginalized parameters

The conditional probability of the marginalized noise parameters can be derived after obtaining $p(\theta \mid \mathcal{D})$ given values of $\theta$.

###### Sampling of the noise parameter

The integrand in (22) shows that the precision $\lambda$ has (up to a constant) the distribution of a Gamma distribution, mainly

$$\lambda \propto \text{Gamma}\left(\alpha' = \alpha + \frac{N}{2}, \beta' = \omega\right).$$

Similarly

$$\sigma^2 \propto \text{Inv-Gamma}\left(\alpha' = \alpha + \frac{N}{2}, \beta' = \omega\right).$$

### 4 Marginalization-based approach for additive Laplace-distributed noise

In this section, we consider *additive Laplace-distributed* measurement noise. The data set $\mathcal{D}$ is a collection of measurements

$$\bar{y}_k = (s \cdot h_k + b) + \epsilon_k, \quad \epsilon_k \sim \text{Laplace}(0, \sigma), \quad \text{with } \sigma \in (0, \infty) \text{ and } k = 1, \ldots, N.$$

Bayes' theorem for unknown model parameter vector $\theta$, unknown scaling factor $s$, unknown offset $\theta$, and unknown noise parameter $\sigma$ is given by

$$p(\theta, s, b, \sigma \mid \mathcal{D}) = \frac{p(\mathcal{D} \mid \theta, s, b, \sigma) p(\theta, s, b, \sigma)}{p(\mathcal{D})},$$

with prior $p(\theta, s, b, \sigma^2)$, marginal $p(\mathcal{D})$ and likelihood

$$p(\mathcal{D} \mid \theta, s, b, \lambda) = \frac{1}{(2\sigma)^N} \cdot \exp\left(\sum_{k=1}^{N} -\frac{|\bar{y}_k - (s \cdot h_k + b)|}{\sigma}\right)$$

In the following, we show different combinations of known and unknown parameters.

#### 4.1 Unknown scaling parameters

We consider the observation and noise model

$$\bar{y}_k = s \cdot h_k + \epsilon_k, \quad \epsilon_k \sim \text{Laplace}(0, \sigma),$$

with unknown scaling parameters $s$. For these unknown parameters we assume a Gaussian prior,

$$\begin{aligned} p(s) &= \mathcal{N}(s \mid \mu, 1/\tau) \\ &= \sqrt{\frac{\tau}{2\pi}} \exp\left(-\frac{\tau}{2}(s - \mu)^2\right) \end{aligned}$$

with $\mu$, $\tau$ as hyper-parameters. The marginalized likelihood is denoted by

$$\begin{aligned} p(\theta \mid \mathcal{D}) &= \int_{-\infty}^{\infty} p(\mathcal{D} \mid \theta, s) p(s) \, \mathrm{d}s \\ &= \frac{1}{(2\sigma)^N} \int_{-\infty}^{\infty} \exp\left(-\frac{1}{\sigma} \sum_{k=1}^{N} |\bar{y}_k - s \cdot h_k|\right) p(s) \, \mathrm{d}s. \end{aligned}$$

The absolute value contains $\bar{y}_k - s \cdot h_k \geq 0 \Leftrightarrow \bar{y}_k / h_k \geq s$. Hence, $\bar{y}_k$ and $h_k$ need to be renumbered so that $\bar{y}_1 / h_1$ is the smallest value and $\bar{y}_N / h_N$ the largest value. Note that $h_k = 0$ is possible and in that case $|\bar{y}_k - s \cdot h_k| = |\bar{y}_k| = \bar{y}_k \geq 0$.

The summation is split up into $k$ such that $h_k \neq 0$ and $k$ such that $h_k = 0$. With this constraint the integral can be split up to remove the absolute value into $N' := \#\{k \in \{1, \ldots, N\} \mid h_k \neq 0\}$ pieces. Then we can **renumber** $\bar{y}_k$ and $h_k$ together in pairs such that $h_1, \ldots, h_{N'} \neq 0$ and $\bar{y}_i / h_i \leq \bar{y}_j / h_j$ for $1 \leq i < j \leq N'$. If $N' \neq N$ we have $h_{N'+1}, \ldots, h_N = 0$, they are ordered such that $\bar{y}_i \leq \bar{y}_j$ for all

$N' < i \leq j \leq N$. We also need to introduce $\rho_0 = -\infty$, $\rho_i = \bar{y}_i/h_i$ (for $i = 1, \dots, N'$), $\rho_{N'+1} = \infty$. Then we obtain

$$p(\theta \mid \mathcal{D}) = \frac{1}{(2\sigma)^N} \sum_{i=0}^{N'} \int_{\rho_i}^{\rho_{i+1}} p(s) \exp\left( -\frac{1}{\sigma} \sum_{k=1}^{N} |\bar{y}_k - s \cdot h_k| \right) \mathrm{d}s.$$

We introduce the index

$$r_{k,i} = \begin{cases} -1, & \text{if } k \leq i \\ 1, & \text{if } k > i \end{cases}$$

to remove the absolute value term

$$\begin{aligned} p(\theta \mid \mathcal{D}) &= \frac{1}{(2\sigma)^N} \sum_{i=0}^{N'} \int_{\rho_i}^{\rho_{i+1}} p(s) \exp\left( -\frac{1}{\sigma} \sum_{k=1}^{N} r_{k,i} \big( \bar{y}_k - s \cdot h_k \big) \right) \mathrm{d}s \\ &= \frac{1}{(2\sigma)^N} \sum_{i=0}^{N'} \int_{\rho_i}^{\rho_{i+1}} p(s) \exp\left( -\frac{1}{\sigma} \left( \left( \sum_{k=i+1}^{N} \bar{y}_k - s \cdot h_k \right) - \left( \sum_{k=1}^{i} \bar{y}_k - s \cdot h_k \right) \right) \right) \mathrm{d}s \\ &= \frac{1}{(2\sigma)^N} \sum_{i=0}^{N'} \int_{\rho_i}^{\rho_{i+1}} p(s) \exp\left( -\frac{1}{\sigma} \left( \left( \sum_{k=i+1}^{N} \bar{y}_k - \sum_{l=1}^{i} \bar{y}_l \right) + s \cdot \left( \sum_{k=1}^{i} h_k - \sum_{l=i+1}^{N} h_l \right) \right) \right) \mathrm{d}s \\ &= \frac{1}{(2\sigma)^N} \sum_{i=0}^{N'} \exp\left( \frac{1}{\sigma} \left( \sum_{l=1}^{i} \bar{y}_l - \sum_{k=i+1}^{N} \bar{y}_k \right) \right) \int_{\rho_i}^{\rho_{i+1}} p(s) \exp\left( \frac{s}{\sigma} \left( \sum_{l=i+1}^{N} h_l - \sum_{k=1}^{i} h_k \right) \right) \mathrm{d}s. \end{aligned}$$

To simplify the notation, we introduce the indexes

$$d_i := \left( \sum_{k=1}^{i} \bar{y}_k \right) - \left( \sum_{k=i+1}^{N} \bar{y}_k \right) \quad \text{for } i = 0, \dots, N'$$

and

$$q_i := \left( \sum_{k=i+1}^{N} h_k \right) - \left( \sum_{k=1}^{i} h_k \right) \quad \text{for } i = 0, \dots, N'.$$

We can insert the Gaussian prior into the integral

$$p(\theta \mid \mathcal{D}) = \frac{1}{(2\sigma)^N} \cdot \sqrt{\frac{\tau}{2\pi}} \sum_{i=0}^{N'} \exp\left( \frac{d_i + \mu q_i}{\sigma} + \frac{q_i^2}{2\sigma^2 \tau} \right) \underbrace{\int_{\rho_i}^{\rho_{i+1}} \exp\left( -\frac{\tau}{2} \left( s - \left( \mu + \frac{q_i}{\sigma\tau} \right) \right)^2 \right) \mathrm{d}s}_{(*)}.$$

Using substitution

$$f_i(z) = \frac{z - \left( \mu + \frac{q_i}{\tau\sigma} \right)}{\sqrt{2/\tau}}$$

in $(*)$, we get

$$(*) = \sqrt{\frac{2}{\tau}} \int_{f_i(\rho_i)}^{f_i(\rho_{i+1})} e^{-z^2} \, \mathrm{d}z.$$

Applying the error function

$$\operatorname{erf}(x) = \frac{2}{\sqrt{\pi}} \int_0^x e^{-z^2} \, \mathrm{d}z, \tag{23}$$

we obtain the following solutions

$$(*) = \begin{cases} \sqrt{\frac{\pi}{2\tau}}\Big(\frac{2}{\sqrt{\pi}}\int_0^{-f_i(\rho_i)} e^{-z^2}\,\mathrm{d}z - \frac{2}{\sqrt{\pi}}\int_0^{-f_i(\rho_{i+1})} e^{-z^2}\,\mathrm{d}z\Big) & \text{in case (i)} \\ \sqrt{\frac{\pi}{2\tau}}\Big(\frac{2}{\sqrt{\pi}}\int_0^{-f_i(\rho_i)} e^{-z^2}\,\mathrm{d}z + \frac{2}{\sqrt{\pi}}\int_0^{f_i(\rho_{i+1})} e^{-z^2}\,\mathrm{d}z\Big) & \text{in case (ii)} \\ \sqrt{\frac{\pi}{2\tau}}\Big(\frac{2}{\sqrt{\pi}}\int_0^{f_i(\rho_{i+1})} e^{-z^2}\,\mathrm{d}z - \frac{2}{\sqrt{\pi}}\int_0^{f_i(\rho_i)} e^{-z^2}\,\mathrm{d}z\Big) & \text{in case (iii)} \end{cases}$$

$$= \begin{cases} \sqrt{\frac{\pi}{2\tau}} \cdot \big(\operatorname{erf}(-f_i(\rho_i)) - \operatorname{erf}(-f_i(\rho_{i+1}))\big) & \text{in case (i)} \\ \sqrt{\frac{\pi}{2\tau}} \cdot \big(\operatorname{erf}(-f_i(\rho_i)) + \operatorname{erf}(f_i(\rho_{i+1}))\big) & \text{in case (ii)} \\ \sqrt{\frac{\pi}{2\tau}} \cdot \big(\operatorname{erf}(f_i(\rho_{i+1})) - \operatorname{erf}(f_i(\rho_i))\big) & \text{in case (iii)} \end{cases}$$

with cases

$$\begin{cases} \text{(i)} := & f_i(\rho_i) \le f_i(\rho_{i+1}) \le 0 \\ \text{(ii)} := & f_i(\rho_i) \le 0 \le f_i(\rho_{i+1}) \\ \text{(iii)} := & 0 \le f_i(\rho_i) \le f_i(\rho_{i+1}) \end{cases}. \tag{24}$$

The substitution of this result for the integral over $s$ in the integral formulation yields a closed-form solution for the marginalized likelihood,

$$p(\theta \mid \mathcal{D}) = \frac{1}{2(2\sigma)^N} \sum_{i=0}^{N'} \exp\left(\frac{d_i + \mu q_i}{\sigma} + \frac{q_i^2}{2\tau\sigma^2}\right) \cdot k_i,$$

with

$$k_i := \begin{cases} \operatorname{erf}(-f_i(\rho_i)) - \operatorname{erf}(-f_i(\rho_{i+1})), & \text{in case (i)} \\ \operatorname{erf}(-f_i(\rho_i)) + \operatorname{erf}(f_i(\rho_{i+1})), & \text{in case (ii)} \\ \operatorname{erf}(f_i(\rho_{i+1})) - \operatorname{erf}(f_i(\rho_i)), & \text{in case (iii)} \end{cases}. \tag{25}$$

##### 4.1.1 Retrieving the marginalized parameters

The conditional probability of the marginalized scaling parameters can be derived after obtaining $p(\theta \mid \mathcal{D})$ given values of $\theta$.

##### Sampling of the scaling parameter

The distribution for $s$ is (up to a constant)

$$s \propto \sum_{i=0}^{N'} \mathbb{1}_{(\rho_i, \rho_i+1]} \sqrt{\frac{\tau}{2\pi}} \exp\left(\frac{d_i + \mu q_i}{\sigma} + \frac{q_i^2}{2\sigma^2\tau}\right) \cdot \exp\left(-\frac{\tau}{2}\left(s - \left(\mu + \frac{q_i}{\sigma\tau}\right)\right)^2\right)$$
$$+ \mathbb{1}_{(b_{N'}, \infty)} \sqrt{\frac{\tau}{2\pi}} \exp\left(\frac{d_{N'} + \mu q_{N'}}{\sigma} + \frac{q_{N'}^2}{2\sigma^2\tau}\right) \cdot \exp\left(-\frac{\tau}{2}\left(s - \left(\mu + \frac{q_{N'}}{\sigma\tau}\right)\right)^2\right).$$

This is a piece-wise Gaussian distribution. To sample this type of distribution we have to calculate the value $I_i$ of each piece separately. We can use the calculation above and receive

$$I_i := \exp\left(\frac{d_i + \mu q_i}{\sigma} + \frac{q_i^2}{2\tau\sigma^2}\right) \cdot \frac{k_i}{2}$$

with $k_i$ defined as in (25). We can sample a uniformly distributed random variable of the interval $[0, \sum_{i=0}^{N'} I_i]$ and determine with this in which interval we will sample. It remains to derive the distribution of $s$ in every interval which can be done with the inverse-cdf. With the previous calculation we get that

$$\mathbb{P}[s \leq \delta] = \begin{cases} \dfrac{\operatorname{erf}(-f_i(\rho_i)) - \operatorname{erf}(-f_i(\delta))}{\operatorname{erf}(-f_i(\rho_i)) - \operatorname{erf}(-f_i(\rho_{i+1}))} & \text{in case (i)} \\ \dfrac{\operatorname{erf}(-f_i(\rho_i)) + \operatorname{erf}(f_i(\delta))}{\operatorname{erf}(-f_i(\rho_i)) + \operatorname{erf}(f_i(\rho_{i+1}))} & \text{in case (ii)} \\ \dfrac{\operatorname{erf}(f_i(\delta)) - \operatorname{erf}(f_i(\rho_i))}{\operatorname{erf}(f_i(\rho_{i+1})) - \operatorname{erf}(f_i(\rho_i))} & \text{in case (iii)} \end{cases}$$

with cases as in (24).

We can sample a uniformly distributed $u \sim U(0,1)$ and insert it into the inverse-cdf $g(u)$ for the three cases. We first notice that the inverse of $f_i$ is

$$f_i^{-1}(u) = u \cdot \sqrt{\frac{2}{\tau}} + \left(\mu + \frac{q_i}{\tau\sigma}\right)$$

while we have to use a numerical approximation to create an inverse of the error function $\operatorname{erf}^{-1}$. In total we can write the inverse-cdf as

$$g(u) := \begin{cases} f_i^{-1}\left(-\operatorname{erf}^{-1}\left(\operatorname{erf}(-f_i(\rho_i)) - u \cdot \left(\operatorname{erf}\left(-f_i(\rho_i)\right) - \operatorname{erf}\left(-f_i(\rho_{i+1})\right)\right)\right)\right) & \text{in case (i)} \\ f_i^{-1}\left(\operatorname{erf}^{-1}\left(u \cdot \left(\operatorname{erf}\left(-f_i(\rho_i)\right) + \operatorname{erf}\left(f_i(\rho_{i+1})\right)\right) - \operatorname{erf}(-f_i(\rho_i))\right)\right) & \text{in case (ii)} \\ f_i^{-1}\left(\operatorname{erf}^{-1}\left(u \cdot \left(\operatorname{erf}\left(-f_i(\rho_{i+1})\right) - \operatorname{erf}\left(f_i(\rho_i)\right)\right) + \operatorname{erf}(-f_i(\rho_i))\right)\right) & \text{in case (iii)} \end{cases} .$$

#### 4.2 Unknown offset parameters

We consider the observation and noise model

$$\bar{y}_k = h_k + b + \epsilon_k, \quad \epsilon_k \sim \text{Laplace}(0, \sigma),$$

with unknown offset parameters $b$. For these unknown parameters we assume a Gaussian prior,

$$\begin{aligned} p(b) &= \mathcal{N}(s \mid \mu, 1/\tau) \\ &= \sqrt{\frac{\tau}{2\pi}} \exp\left(-\frac{\tau}{2}(b-\mu)^2\right) \end{aligned}$$

with $\mu$, $\tau$ as hyper-parameters. The marginalized likelihood is denoted by

$$\begin{aligned} p(\theta \mid \mathcal{D}) &= \int_{-\infty}^{\infty} p(\mathcal{D} \mid \theta, b) p(b) \,\mathrm{d}b \\ &= \int_{-\infty}^{\infty} \left( \prod_{k=1}^{N} \frac{1}{2\sigma} \cdot \exp\left(-\frac{|b - (\bar{y}_k - h_k)|}{\sigma}\right)\right) p(b) \,\mathrm{d}b. \end{aligned}$$

For calculation-reasons we **renumber** $\bar{y}_k, h_k$ together in pairs so that $\bar{y}_k - h_k$ are ordered from smallest to largest, i.e. $\bar{y}_1 - h_1$ is the smallest number, $\bar{y}_N - h_N$ the largest. Then we choose $\rho_0 = -\infty, \rho_i = \bar{y}_i - h_i$ for $(i = 1, \ldots, N), \rho_{N+1} = \infty$. Then, we get

$$p(\theta \mid \mathcal{D}) = \frac{1}{(2\sigma)^N} \left( \sum_{i=0}^{N} \int_{\rho_i}^{\rho_{i+1}} \exp\left(-\frac{\sum_{k=1}^{N} |b - \rho_k|}{\sigma}\right) p(b) \,\mathrm{d}b \right).$$

We introduce the index

$$q_{k,i} = \begin{cases} 1, & \text{if } k \le i \\ -1, & \text{if } k > i \end{cases}$$

to remove the absolute value

$$\begin{aligned} p(\theta \mid \mathcal{D}) &= \frac{1}{(2\sigma)^N} \left( \sum_{i=0}^{N} \int_{\rho_i}^{\rho_{i+1}} \exp\left(-\frac{\sum_{k=1}^{N} q_{k,i}(b - \rho_k)}{\sigma}\right) p(b) \,\mathrm{d}b \right) \\ &= \frac{1}{(2\sigma)^N} \left( \sum_{i=0}^{N} \int_{\rho_i}^{\rho_{i+1}} \exp\left(\frac{b \cdot ((N-i) - i) + \left(\sum_{k=1}^{i} \rho_k\right) - \left(\sum_{k=i+1}^{N} \rho_k\right)}{\sigma}\right) p(b) \,\mathrm{d}b \right) \\ &= \frac{1}{(2\sigma)^N} \left( \sum_{i=0}^{N} \exp\left(\frac{\left(\sum_{k=1}^{i} \rho_k\right) - \left(\sum_{k=i+1}^{N} \rho_k\right)}{\sigma}\right) \int_{\rho_i}^{\rho_{i+1}} \exp\left(b \cdot \frac{N - 2i}{\sigma}\right) p(b) \,\mathrm{d}b \right). \end{aligned} \quad (26)$$

To simplify the notation, we introduce the index

$$l_i := \left( \sum_{k=1}^{i} \rho_k \right) - \left( \sum_{k=i+1}^{N} \rho_k \right) \quad \text{for } i = 0, \ldots, N.$$

We can insert the Gaussian prior into the integral

$$
\begin{aligned}
p(\theta \mid \mathcal{D}) =& \sqrt{\frac{\tau}{2\pi}} \frac{1}{(2\sigma)^N} \sum_{i=0}^{N} \exp\left(\frac{l_i}{\sigma}\right) \int_{\rho_i}^{\rho_{i+1}} \exp\left(b \cdot \frac{N-2i}{\sigma} - \frac{\tau}{2}(b-\mu)^2\right) \mathrm{d}b \\
=& \sqrt{\frac{\tau}{2\pi}} \frac{1}{(2\sigma)^N} \sum_{i=0}^{N} \exp\left(\frac{l_i}{\sigma}\right) \int_{\rho_i}^{\rho_{i+1}} \exp\left(-\frac{\tau}{2}\left(b^2 - 2\left(\frac{N-2i}{\tau\sigma} + \mu\right) b + \mu^2\right)\right) \mathrm{d}b \\
=& \sqrt{\frac{\tau}{2\pi}} \frac{1}{(2\sigma)^N} \sum_{i=0}^{N} \exp\left(\frac{l_i}{\sigma}\right) \exp\left(\frac{(N-2i)^2}{2\tau\sigma^2} + \frac{\mu(N-2i)}{\sigma}\right) \\
& \cdot \underbrace{\int_{\rho_i}^{\rho_{i+1}} \exp\left(-\frac{\tau}{2}\left(b - \left(\frac{N-2i}{\tau\sigma} + \mu\right)\right)^2\right) \mathrm{d}b}_{(*)}.
\end{aligned}
$$

Using substitution

$$
f_i(z) = \frac{z - \left(\frac{N-2i}{\tau\sigma} + \mu\right)}{\sqrt{2/\tau}}
$$

in $(*)$, we get

$$
(*) = \sqrt{\frac{2}{\tau}} \int_{f_i(\rho_i)}^{f_i(\rho_{i+1})} e^{-z^2} \, \mathrm{d}z.
$$

Applying the error function (23), we obtain the following solutions

$$
\begin{aligned}
(*) &= \begin{cases}
\sqrt{\frac{\pi}{2\tau}} \Big( \frac{2}{\sqrt{\pi}} \int_0^{-f_i(\rho_i)} e^{-z^2} \, \mathrm{d}z - \frac{2}{\sqrt{\pi}} \int_0^{-f_i(\rho_{i+1})} e^{-z^2} \, \mathrm{d}z \Big) & \text{in case (i)} \\
\sqrt{\frac{\pi}{2\tau}} \Big( \frac{2}{\sqrt{\pi}} \int_0^{-f_i(\rho_i)} e^{-z^2} \, \mathrm{d}z + \frac{2}{\sqrt{\pi}} \int_0^{f_i(\rho_{i+1})} e^{-z^2} \, \mathrm{d}z \Big) & \text{in case (ii)} \\
\sqrt{\frac{\pi}{2\tau}} \Big( \frac{2}{\sqrt{\pi}} \int_0^{f_i(\rho_{i+1})} e^{-z^2} \, \mathrm{d}z - \frac{2}{\sqrt{\pi}} \int_0^{f_i(\rho_i)} e^{-z^2} \, \mathrm{d}z \Big) & \text{in case (iii)}
\end{cases} \\
&= \begin{cases}
\sqrt{\frac{\pi}{2\tau}} \cdot \big( \mathrm{erf}(-f_i(\rho_i)) - \mathrm{erf}(-f_i(\rho_{i+1})) \big) & \text{in case (i)} \\
\sqrt{\frac{\pi}{2\tau}} \cdot \big( \mathrm{erf}(-f_i(\rho_i)) + \mathrm{erf}(f_i(\rho_{i+1})) \big) & \text{in case (ii)} \\
\sqrt{\frac{\pi}{2\tau}} \cdot \big( \mathrm{erf}(f_i(\rho_{i+1})) - \mathrm{erf}(f_i(\rho_i)) \big) & \text{in case (iii)}
\end{cases}
\end{aligned}
$$

with cases

$$
\begin{cases}
\text{(i)} := & f_i(\rho_i) \le f_i(\rho_{i+1}) \le 0 \\
\text{(ii)} := & f_i(\rho_i) \le 0 \le f_i(\rho_{i+1}) \\
\text{(iii)} := & 0 \le f_i(\rho_i) \le f_i(\rho_{i+1})
\end{cases} . \tag{27}
$$

The substitution of this result for the integral over $b$ in the integral formulation yields a closed-form solution for the marginalized likelihood,

$$
p(\theta \mid \mathcal{D}) = \frac{1}{2(2\sigma)^N} \sum_{i=0}^{N} \exp\left(\frac{(N-2i)^2}{2\tau\sigma^2} + \frac{\mu(N-2i) + l_i}{\sigma}\right) \cdot k_i,
$$

with

$$k_i := \begin{cases} \operatorname{erf}(-f_i(\rho_i)) - \operatorname{erf}(-f_i(\rho_{i+1})) & \text{in case (i)} \\ \operatorname{erf}(-f_i(\rho_i)) + \operatorname{erf}(f_i(\rho_{i+1})) & \text{in case (ii)} \\ \operatorname{erf}(f_i(\rho_{i+1})) - \operatorname{erf}(f_i(\rho_i)) & \text{in case (iii)} \end{cases} . \tag{28}$$

##### 4.2.1 Retrieving the marginalized parameters

The conditional probability of the marginalized offset parameters can be derived after obtaining $p(\theta \mid \mathcal{D})$ given values of $\theta$.

**Sampling of the offset parameter**

The distribution for $b$ is (up to a constant)

$$\begin{aligned} b \propto \sum_{i=0}^{N-1} \mathbb{1}_{(\rho_i,\rho_i+1]} \sqrt{\frac{\tau}{2\pi}} \exp\left(\frac{l_i}{\sigma} + \frac{(N-2i)^2}{2\tau\sigma^2} + \frac{\mu(N-2i)}{\sigma}\right) \cdot \exp\left(-\frac{\tau}{2}\left(b - \left(\frac{N-2i}{\tau\sigma} + \mu\right)\right)^2\right) \\ + \mathbb{1}_{(\rho_{N-1},\infty)} \sqrt{\frac{\tau}{2\pi}} \exp\left(\frac{l_{N-1}}{\sigma} + \frac{(-N+2)^2}{2\tau\sigma^2} + \frac{\mu(-N+2)}{\sigma}\right) \cdot \exp\left(-\frac{\tau}{2}\left(b - \left(\frac{-N+2}{\tau\sigma} + \mu\right)\right)^2\right). \end{aligned}$$

This is a piece-wise Gaussian distribution. To sample this type of distribution we first have to calculate the value $I_i$ of each piece separately. We can just use the calculation above and receive

$$I_i := \exp\left(\frac{(N-2i)^2}{2\tau\sigma^2} + \frac{\mu(N-2i) + l_i}{\sigma}\right) \cdot \frac{k_i}{2}$$

with $k_i$ defined as in (28). We can sample a uniformly distributed random variable of the interval $[0, \sum_{i=0}^{N} I_i]$ and determine with this in which interval we will sample. It remains to derive the distribution of $b$ in every interval. We can do this with the inverse-cdf. With the calculation we did above we get that

$$\mathbb{P}[b \leq \delta] = \begin{cases} \dfrac{\operatorname{erf}(-f_i(\rho_i)) - \operatorname{erf}(-f_i(\delta))}{\operatorname{erf}(-f_i(\rho_i)) - \operatorname{erf}(-f_i(\rho_{i+1}))} & \text{in case (i),} \\ \dfrac{\operatorname{erf}(-f_i(\rho_i)) + \operatorname{erf}(f_i(\delta))}{\operatorname{erf}(-f_i(\rho_i)) + \operatorname{erf}(f_i(\rho_{i+1}))} & \text{in case (ii),} \\ \dfrac{\operatorname{erf}(f_i(\delta)) - \operatorname{erf}(f_i(\rho_i))}{\operatorname{erf}(f_i(\rho_{i+1})) - \operatorname{erf}(f_i(\rho_i))} & \text{in case (iii).} \end{cases}$$

with cases as in (27).

We can sample a uniformly distributed $u \sim U(0,1)$ and insert it into the inverse-cdf $g(u)$ for the three cases. We first notice that the inverse of $f_i$ is

$$f_i^{-1}(u) = u \cdot \sqrt{\frac{2}{\tau}} + \left(\frac{N-2i}{\tau\sigma} + \mu\right)$$

while we have to use a numerical approximation to create an inverse of the error function $\operatorname{erf}^{-1}$. In total we can write the inverse-cdf as

$$g(u) := \begin{cases} f_i^{-1}\left(-\operatorname{erf}^{-1}\left(\operatorname{erf}(-f_i(\rho_i)) - u \cdot \left(\operatorname{erf}\left(-f_i(\rho_i)\right) - \operatorname{erf}\left(-f_i(\rho_{i+1})\right)\right)\right)\right) & \text{in case (i)} \\ f_i^{-1}\left(\operatorname{erf}^{-1}\left(u \cdot \left(\operatorname{erf}\left(-f_i(\rho_i)\right) + \operatorname{erf}\left(f_i(\rho_{i+1})\right)\right) - \operatorname{erf}(-f_i(\rho_i))\right)\right) & \text{in case (ii)} \\ f_i^{-1}\left(\operatorname{erf}^{-1}\left(u \cdot \left(\operatorname{erf}\left(-f_i(\rho_{i+1})\right) - \operatorname{erf}\left(f_i(\rho_i)\right)\right) + \operatorname{erf}(-f_i(\rho_i))\right)\right) & \text{in case (iii)} \end{cases} .$$

### 5 Benchmark models

#### Conversion reaction model

The model for the conversion reaction describes a reversible chemical reaction, which converts a biochemical species $A$ to a species $B$ with rate $\theta_1$, and $B$ to $A$ with rate $\theta_2$ . The corresponding ODEs are

$$\dot{x}_1 = -\theta_1 x_1 + \theta_2 x_2$$
$$\dot{x}_2 = \theta_1 x_1 - \theta_2 x_2$$

for which the state vector $x = (x_1, x_2)^T$ consists of the concentrations of A and B, respectively. We assumed that $x_2$ is measured yielding the observation model $y(t) = s \cdot h(x(t,\theta),\theta) + b = s \cdot x_2(t) + b$. For the evaluation of the proposed method, we generated one artificial dataset. The dataset was generated with initial conditions $x_0 = (1, 0.01)^T$, model parameters $\theta = (\theta_1, \theta_2)^T = (10^{-0.3}, 10^{-0.5})^T$, scaling parameter $s = 10^{0.3}$, offset parameter $b = 10^{0.17}$, and additive Gaussian noise with variance $\sigma^2 = 10^{-3.0}$.

#### Model of mRNA transfection

The M3 model presented in the main manuscript describes an mRNA transfection process in which cells are transfected with GFP-mRNA. The corresponding ODEs are

$$\dot{x}_1 = -\delta \cdot x_1 \qquad x_1(t_0) = m_0$$
$$\dot{x}_2 = k_{TL} \cdot x_1 - \beta \cdot x_2 \qquad x_2(t_0) = 0$$

where $t_0$ denotes the onset time, $k_{TL}$ the translation rate of mRNA to protein, $m_0$ the initial number of effectively translated mRNA molecules, and $\delta$ and $\beta$ the degradation rates of mRNA and protein, respectively. The state vector $x = (x_1, x_2)^T$ consists of the concentrations of mRNA and protein, respectively. The observable is the protein concentration, which is given by

$$y(t) = s \cdot x_2(t) = s \cdot k_{TL} \cdot m_0 \cdot \frac{e^{-\beta(t-t_0)} - e^{-\delta(t-t_0)}}{\delta - \beta}.$$

The posterior distribution of this model is bimodal as the exchange of the degradation rates of mRNA $\delta$ and protein $\beta$ results in the same dynamics for the observable. We considered the product $\hat{s} = s \cdot k_{TL} \cdot m_0$ as a single parameter and treated $\hat{s}$ as a scaling parameter, therefore marginalized together with the noise parameter. Consequently, in our setting this model consists of 5 unknown parameters in total, of which we have 3 model parameters $\theta = (t_0, \beta, \delta)$, one scaling parameter $\hat{s}$, and one noise parameter (main manuscript Table 1, M3).
